# Understanding Cognitive Impairment in Mood Disorders: Mediation Analyses in the UK Biobank Cohort

**DOI:** 10.1101/655290

**Authors:** Breda Cullen, Daniel J. Smith, Ian J. Deary, Jill P. Pell, Katherine M. Keyes, Jonathan J. Evans

**Affiliations:** Lecturer, Institute of Health and Wellbeing, University of Glasgow, Glasgow, UK; Professor, Institute of Health and Wellbeing, University of Glasgow, Glasgow, UK; Professor, Department of Psychology and Centre for Cognitive Ageing and Cognitive Epidemiology, University of Edinburgh, Edinburgh, UK; Henry Mechan Professor of Public Health and Director, Institute of Health and Wellbeing, University of Glasgow, Glasgow, UK; Associate Professor, Department of Epidemiology, Mailman School of Public Health, Columbia University, New York, NY, USA

## Abstract

**Background:** Cognitive impairment is strongly linked with persistent disability in people with mood disorders, but the factors that explain cognitive impairment in this population are unclear.

**Aims:** We aimed to estimate the total effect of (i) bipolar disorder (BD) and (ii) major depression on cognitive function, and the magnitude of the effect that was explained by potentially modifiable intermediate factors.

**Method:** Cross-sectional study using baseline data from the UK Biobank cohort. Participants were categorised as BD (N=2,709), major depression (N=50,975), or no mood disorder (N=102,931 to 105,284). The outcomes were computerised tests of reasoning, reaction time and memory. The potential mediators were cardiometabolic disease and psychotropic medication. Analyses were informed by graphical methods, and controlled for confounding using regression, propensity score-based methods, and G-computation.

**Results:** Group differences of small magnitude were found on a visuospatial memory test. Z-score differences for BD were in the range −0.23 to −0.17 (95% CI range −0.39 to −0.03) across different estimation methods, and approximately −0.07 (95% CI −0.10 to −0.03) for major depression. One-quarter of the effect was mediated via psychotropic medication in the BD group (−0.05; 95% CI −0.09 to −0.01). No evidence was found for mediation via cardiometabolic disease.

**Conclusions:** In a large community-based sample in middle to early old age, BD and depression were associated with lower visuospatial memory performance, in part potentially due to psychotropic medication use. Mood disorders and their treatments will have increasing importance for population cognitive health as the proportion of older adults continues to grow.

## Introduction

Cognitive impairment is a key determinant of occupational and social outcomes and quality of life in people with mood disorders.^1,2^ Up to 57% of adults with bipolar disorder (BD) and up to one-half of adults with major depression show clinically significant levels of cognitive impairment, even in euthymia.^3,4^ In BD, impairment is typically found on tests of attention, working and episodic memory, processing speed, and executive function, with group differences of medium to large effect size compared with adults without a history of psychiatric illness.^5,6^ The profile of impairment is similar in major depression, but with effect sizes of smaller magnitude.^7^ The causal nature of this relationship is unclear, however, because approaches to accounting for confounding influences have been inconsistent in the literature. Moving towards causal explanations requires careful modelling of a range of potential confounding, mediating and moderating factors, acknowledging the complexity of their inter-relationships with mood disorder exposure, cognitive outcome, and each other. The application of novel confounder control techniques in psychiatric epidemiology is not yet widespread. The counterfactual approach, in which causal effects are conceptualized as alternative ‘potential outcomes’ of an exposure, can be linked with graphical notation in the form of directed acyclic graphs (DAG) to systematically identify causal effects in complex systems.^8,9^ This, in turn, informs rigorous statistical analyses and clarifies the assumptions needed to interpret estimates as causal. In this study, we applied a graphical approach to understand the structure of confounding and mediation with regard to cognitive performance in UK Biobank^10^ participants with a history of BD or major depression. We estimated the total effect of BD and major depression on cognitive function, and the magnitude of the effect that was transmitted through potentially modifiable intermediate factors.

## Method

### Participants

UK Biobank recruited adults from the general population across 22 centres in Great Britain between 2006 and 2010. The target age range was 40 to 69 years and no other exclusion criteria were applied. Postal invitation lists were generated from National Health Service (NHS) registers, with a response rate of approximately 6%. Cross-sectional data from the full cohort at baseline (N=502,618) were used. Participants were included in the analysis if they had sufficient data to classify their BD or major depression exposure status (see below) and had data on at least one cognitive outcome measure. The authors assert that all procedures contributing to this work comply with the ethical standards of the relevant national and institutional committees on human experimentation and with the Helsinki Declaration of 1975, as revised in 2008. All procedures involving human participants were approved by the North West - Haydock NHS Research Ethics Committee (reference 16/NW/0274 and 11/NW/0382). Written informed consent was obtained from all participants.

### Mood Disorder Status

Three sources of information were available regarding history of mood disorder: self-reported doctor diagnosis; probable classification based on self-reported lifetime mood disorder symptoms and help-seeking;^11^ and linked NHS hospital inpatient and day case records. The definitions used in these sources are provided in the eMethods in the Supplement. To permit consistency across each information source, single manic episode and BD were analysed as one exposure (mania/BD), and the major depression exposure included single episode and recurrent illness.

Participants were classified as exposed if they were positive for mania/BD or major depression in at least one information source. Mania/BD and major depression were then separated hierarchically into two mutually exclusive exposure groups. Participants with a record of both mania/BD and major depression were classified in the mania/BD group only. The unexposed comparison group comprised participants who had complete self-reported data that did not indicate mania/BD or major depression, and whose hospital records had no primary or secondary diagnosis of mania/BD or major depression. Furthermore, because misclassification is common between mania/BD and schizophrenia spectrum disorders,^12^ participants with a self-reported diagnosis or hospital record of schizophrenia (ICD-10 code F20x) were excluded from all exposed and unexposed groups. In summary, the final groups for analysis were: mania/BD (excluding participants with only major depression, and those with schizophrenia); major depression (excluding participants with mania/BD, and those with schizophrenia); and unexposed (complete data indicating no mania/BD or major depression or schizophrenia). Participants who did not meet the above criteria for either the exposed or unexposed groups were not further analysed (e.g. those with incomplete data, preventing inclusion in the unexposed group).

### Cognitive Outcome Measures

The cognitive measures analysed were reasoning, reaction time, numeric memory, visuospatial memory and prospective memory, as described in detail elsewhere.^13^ All were administered via a touchscreen computer. The psychometric properties of these tests have been reported previously.^14^ The data were provided by UK Biobank as raw scores, and for the purposes of the present analysis were standardized within five-year age strata, using all available data in the cohort at baseline. Five-year bands were deemed appropriate in light of the typical rate of age-related change in cognitive performance in middle to older adulthood.^15^ To address skew in the raw data distributions, the scores were first transformed into percentiles and then into z-scores (mean = 0 and standard deviation [SD] = 1). The scores for reaction time and visuospatial memory were reflected so that higher scores represent better performance, in line with the other tests. It was not possible to standardize the prospective memory data in this way because responses were dichotomized (correct response at the first attempt or not), and so the raw data were used in the analyses involving this test.

### Covariates

#### Sociodemographic, environmental, lifestyle and physical measures

Details of these measures are provided in the eMethods in the Supplement. Briefly, the sociodemographic variables were age, gender, ethnic background, country of birth, educational attainment, and neighbourhood deprivation level. Local environment measures comprised population density, proximity to the nearest major road, and air pollutants (particulate matter and nitrogen dioxide). Lifestyle and physical measures comprised smoking, alcohol consumption, insomnia, physical activity, and body mass index (BMI).

#### Medical and family history

A dichotomous indicator was created for history of any cardiometabolic disease (self-reported diagnosis of angina, hypertension or non-gestational diabetes, or adjudicated diagnosis of myocardial infarction or stroke; see eMethods in the Supplement). A dichotomous indicator was also created for history of any neurological or psychiatric condition (apart from mood disorder or schizophrenia) in the self-reported or hospital records data; the conditions included are listed in the eMethods in the Supplement. Family history of certain illnesses in biological parents and siblings was included in the baseline questionnaire, and for the present analyses dichotomous indicators were generated for history of psychiatric or neurological conditions (dementia, Parkinson’s disease, or severe depression, coded separately) in any parent or sibling. Participants also self-reported whether their mothers had smoked regularly around the time of their birth.

#### Mental health and psychotropic medication

In addition to the self-reported and hospital records data regarding psychiatric diagnoses, participants also provided self-reported information at baseline about depressive symptoms in the past two weeks, and current psychotropic medications (dichotomous indicator for any mood stabilizer, antidepressant, antipsychotic, sedative or hypnotic), as detailed in the eMethods in the Supplement. Self-reported information regarding number of episodes of depressed mood or anhedonia was collected at baseline and in a web-based follow-up questionnaire in 2016, and data regarding childhood trauma experiences were also collected in the web-based questionnaire (see eMethods).

#### Genome-wide polygenic scores

Genome-wide polygenic scores (GPS) were generated for cognitive ability, BD, and major depression, based on summary statistics from previous genome-wide association studies (GWAS). Full details of the genotyping data, GPS methods, and optimum scores used in the present analyses are provided in the eMethods in the Supplement.

### Statistical Analyses

#### Graphical models

The analyses were informed by a graphical model in the form of a DAG, which is used to visually represent qualitative causal assumptions.^16^ The structural nature of these assumptions permits the detection of implied patterns of dependency and independency among variables, which can then be tested with data. Structural analysis of the DAG allows confounders, mediators and colliders to be distinguished when planning multivariable analyses.^16^

A DAG was constructed to represent plausible causal assumptions about the relationship between lifetime history of mania/BD and cognitive performance, in the context of possible confounding factors and intermediate pathways. This was done before any data were analysed. The nodes in the DAG and the assumed directional relationships between them were determined from previous systematic reviews of cognitive function in BD,^3,5,17-24^ as well as general background knowledge and assumptions regarding other shared causes that were necessary to depict in order for the DAG to have a causal interpretation.^16^ The fit of the DAG to the data was then evaluated by estimating partial correlation coefficients for each pair of nodes that were predicted to be independent. Detection of a correlation between nodes that were predicted to be independent may indicate that the DAG has been misspecified. Where the results indicated lack of independence (i.e. partial correlation coefficient >|0.1|^25^), follow-up regression models were conducted to obtain further detail. Modifications to the structure of the DAG were then considered. A detailed account of the construction and evaluation of the DAG is provided in the eMethods in the Supplement.

The DAG used in the major depression analyses was based on the final DAG used in the mania/BD analyses, with an arrow added from gender to major depression; studies have consistently shown higher prevalence of depression in women,^26^ and it was assumed in the DAG that this relationship was at least partly causal. No arrows were removed compared with the mania/BD DAG, on the assumption that similar causal relationships might be operating to explain cognitive impairment in both disorders.

#### Total effects

The total effect of each mood disorder exposure on cognitive performance was firstly identified in the DAGs using DAGitty software.^27^ The DAGitty algorithm applies ‘d-separation’ rules^8,28^ to find all the confounding paths between the exposure and the outcome, and ascertains if there is a set of nodes which, if conditioned on in the analysis, would ‘block’ these confounding paths. This information was then used to plan regression- and matching-based analyses to estimate the effect in the dataset. This was estimated separately for each of the five cognitive outcome measures, to allow for the possibility of task-specific variation in the results. Results are reported as standardized mean differences or risk differences with 95% confidence intervals (CI). Further details of the estimation methods are given in the eMethods in the Supplement.

#### Mediation analyses

The DAGs were used to assess whether indirect effects via various mediators of interest could be identified. This required all confounding paths between the exposure and the mediator and between the mediator and the outcome to be blocked, as well as those between the exposure and the outcome. Where this requirement was satisfied (i.e. covariate adjustment sets could be found), G-computation was used to estimate the natural direct and indirect effects.^29^ This was implemented using the Stata package `gformula`;^30^ `gformula` permits mediation analysis in the presence of intermediate confounding, whereby a mediator-outcome confounder is itself caused by the exposure. Results are reported as standardized mean differences or risk differences with 95% CI.

#### Sensitivity analyses

Additional analyses were conducted to evaluate the sensitivity of the results to various potential sources of bias, including residual confounding, missing data, and exposure misclassification (see eMethods in the Supplement). These informed our interpretation with regard to key threats to the validity of the analytic framework.

Reporting follows STROBE guidelines.^31^

## Results

### Cognitive Impairment in Mania/BD

#### Characteristics of the sample

eFig. 2 in the Supplement shows a flowchart of exclusions leading to the final analysis sample, which comprised 2,709 participants with mania/BD and 105,284 comparison participants. The descriptive results indicated worse cognitive performance and less favourable covariate characteristics in the mania/BD group, although they were younger on average and more likely to have a degree (Table 1 and eTable 5).

**Table 1.** Summary of Cognitive Outcome Measures in the Mania/Bipolar Disorder and Comparison Groups

|  | All available data |  | Complete covariate data<sup>a</sup> |  |
| --- | --- | --- | --- | --- |
|  | Mania/BD | Comparison | Mania/BD | Comparison |
| N | 2,709 | 105,284 | 504 | 26,997 |
| Reasoning z-score |  |  |  |  |
| N (%) missing<sup>b</sup> | 31 (1.8) | 1,627 (1.6) | 1 (0.3) | 89 (0.3) |
| Mean (SD) | -0.35 (1.01) | -0.20 (0.97) | 0.05 (0.96) | 0.12 (0.92) |
| Cronbach's $\alpha$ | 0.71 | 0.70 | 0.70 | 0.68 |
| Reaction time z-score |  |  |  |  |
| N (%) missing | 51 (1.9) | 1,167 (1.1) | 1 (0.2) | 72 (0.3) |
| Mean (SD) | -0.19 (1.01) | -0.03 (0.98) | 0.03 (0.94) | 0.07 (0.96) |
| Cronbach's $\alpha$ | 0.82 | 0.82 | 0.72 | 0.82 |
| Numeric memory z-score |  |  |  |  |
| N (%) missing<sup>b</sup> | 18 (3.9) | 755 (2.4) | 0 (0.0) | 51 (0.7) |
| Mean (SD) | -0.52 (1.00) | -0.35 (0.94) | -0.23 (1.08) | -0.16 (0.93) |
| Visuospatial memory z-score |  |  |  |  |
| N (%) missing | 157 (5.8) | 2,896 (2.8) | 10 (1.2) | 212 (0.8) |
| Mean (SD) | 0.07 (1.07) | 0.26 (1.05) | 0.20 (1.09) | 0.37 (1.03) |
| Prospective memory<sup>b,c</sup> |  |  |  |  |
| N (%) correct | 1,264 (72.1) | 80,502 (76.9) | 286 (80.3) | 23,112 (85.7) |

BD, bipolar disorder; GPS, genome-wide polygenic score; SD, standard deviation.
- a. Participants with complete data on all the covariates that were entered into the maximally-adjusted total effects models (age, gender, white British genetic ancestry, English-speaking country of birth, degree, comorbid neurological/psychiatric condition, family history of dementia, family history of Parkinson's disease, family history of severe depression, maternal smoking around birth, childhood trauma, education/cognition GPS, bipolar disorder GPS). - b. Missing data refers only to the period when this measure was included in the battery. - c. No missing data.

#### Evaluation of the graphical model

The original DAG is shown and explained in the eMethods in the Supplement. The different predicted independencies implied by alternative plausible specifications of this DAG were tested (see eResults in the Supplement for details). eFig. 3 in the Supplement shows the best fitting DAG, which was used as the basis for the analysis models.

#### Total effects

Only the visuospatial memory test (Fig. 1) and, more equivocally, the prospective memory test (eFig. 4 in the Supplement) indicated a detrimental effect of mania/BD that remained evident in the multivariable models. The effect sizes were small: the mania/BD group scored approximately 0.2 SD lower than the unexposed comparison group on the visuospatial memory test, and the proportion of the mania/BD group succeeding on the prospective memory task was lower by approximately 5 percentage points (approximately 82% in the mania/BD group versus 87% in the unexposed group). The visuospatial and prospective memory estimates showed little change between the unadjusted and adjusted/matched models, whereas the estimates for the other three cognitive measures generally attenuated towards the null (see eFigs. 5 to 7 in the Supplement).

**Fig. 1.**
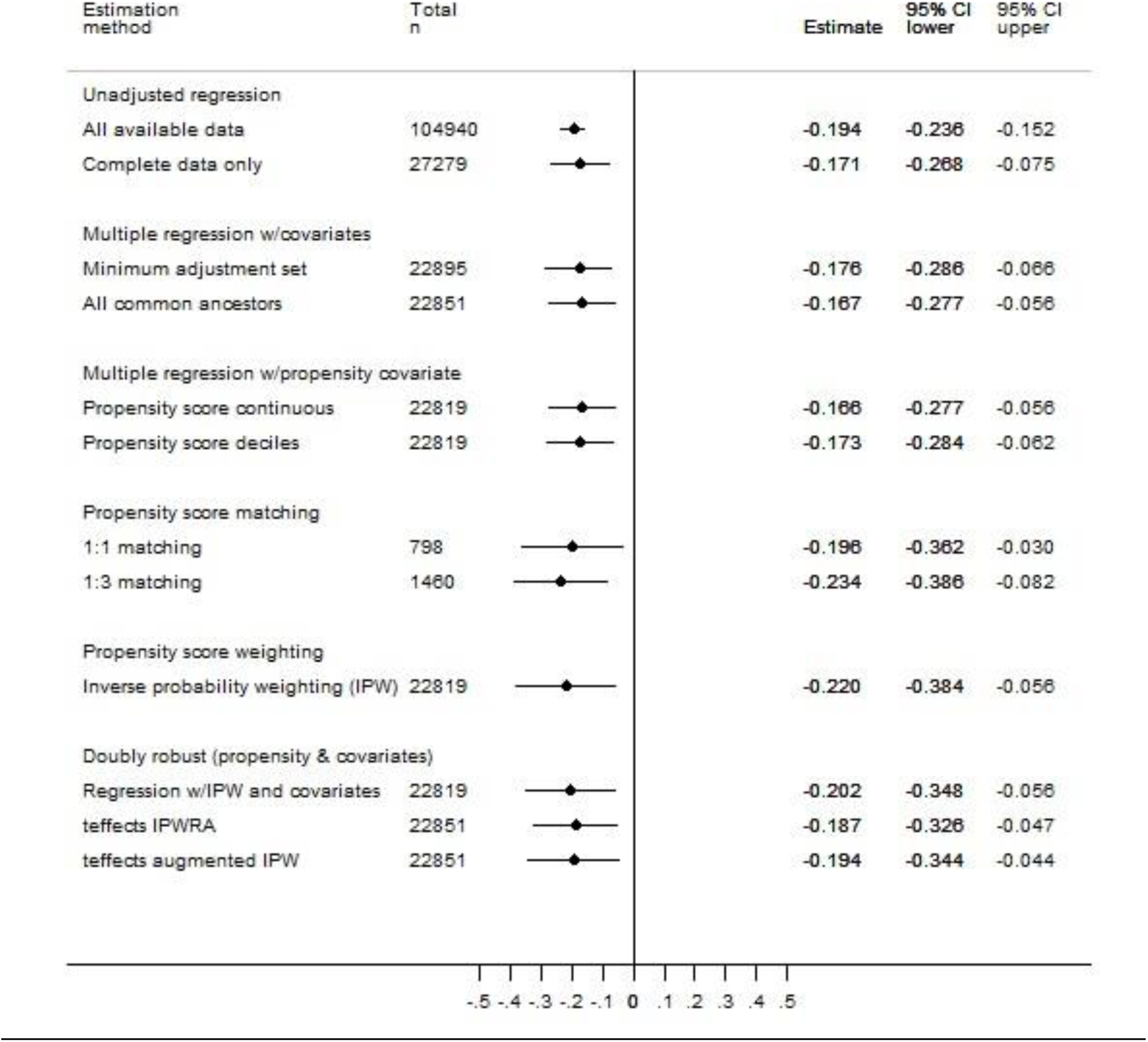
Total Effect of Mania/Bipolar Disorder on Visuospatial Memory. CI, confidence interval; GPS, genome-wide polygenic score; IPW, inverse probability weighting; IPWRA, inverse probability weighting with regression adjustment; teffects, Stata `teffects` package. Estimates are in z-score units and can be interpreted as standardized mean differences. The minimum sufficient adjustment set comprised gender, educational attainment, English-speaking birth country, ethnicity, education/cognition GPS, bipolar disorder GPS, family history of dementia or Parkinson’s disease, maternal smoking around birth, childhood trauma, and comorbid psychiatric/neurological conditions. The extended adjustment set (‘all common ancestors’) also included age and family history of depression. This extended adjustment set was used as the predictor set for the propensity score model. Ethnicity was accounted for in all the multivariable analyses and in the propensity score estimation by restricting these to participants of white British genetic ancestry. The GPS scores were residualized as described in the eMethods, and were entered as deciles, based on the distribution in the full analysis sample.

#### Mediation analyses

Structural analysis of the DAG indicated that direct and indirect effects could be decomposed for two potentially modifiable mediators: cardiometabolic disease and psychotropic medication. Further details of these models are provided in the eResults in the Supplement.

There was no evidence of substantive indirect effects via cardiometabolic disease in any of the models (eTable 8 in the Supplement).

There was evidence that the previously-noted detrimental effect of mania/BD on visuospatial memory was indirectly transmitted via psychotropic medication (eTable 9 in the Supplement). Of the estimated total effect of −0.19 SD units (95% CI −0.31 to −0.08), approximately one quarter was mediated via psychotropic medication (−0.05; 95% CI −0.09 to −0.01). Indirect effects were also evident in the reasoning, reaction time and prospective memory models, although the total effects estimates in these models did not show reliable decrements for mania/BD.

#### Sensitivity analyses

The results of the sensitivity analyses are provided in the eResults in the Supplement. Briefly, these indicated that: the total effects results for visuospatial memory are likely to be sensitive to exposure misclassification and would not be robust to an unmeasured confounder with even a weak association with exposure group membership (leading to minimally unbalanced odds of exposure, i.e. 45/55); the total effects results showed less attenuation when missing covariate data were imputed; the mediation results with missing covariate data imputation showed stronger evidence for an indirect effect via psychotropic medication but no evidence of an indirect effect via cardiometabolic disease; the DAGitty algorithm determined that there were six other DAGs that were equivalent to the DAG shown in eFig. 3 in the Supplement, but none of these alternative configurations was causally plausible.

### Cognitive Impairment in Major Depression

#### Characteristics of the sample

eFig. 9 in the Supplement shows that the analysis sample comprised 50,975 participants with major depression and 102,931 comparison participants. Table 2 summarizes their cognitive outcome data, and eTable 12 in the Supplement summarizes their covariate data.

**Table 2.** Summary of Cognitive Outcome Measures in the Major Depression and Comparison Groups

|  | All available data |  | Complete covariate data<sup>a</sup> |  |
| --- | --- | --- | --- | --- |
|  | Major depression | Comparison | Major depression | Comparison |
| N | 50,975 | 102,931 | 11,662 | 26,392 |
| Reasoning z-score |  |  |  |  |
| N (%) missing<sup>b</sup> | 421 (1.3) | 1,584 (1.6) | 25 (0.3) | 84 (0.3) |
| Mean (SD) | -0.20 (0.96) | -0.20 (0.97) | 0.10 (0.90) | 0.13 (0.92) |
| Cronbach's $\alpha$ | 0.69 | 0.70 | 0.66 | 0.68 |
| Reaction time z-score |  |  |  |  |
| N (%) missing | 534 (1.1) | 1,139 (1.1) | 34 (0.3) | 68 (0.3) |
| Mean (SD) | -0.08 (0.98) | -0.03 (0.98) | 0.04 (0.94) | 0.07 (0.95) |
| Cronbach's $\alpha$ | 0.82 | 0.82 | 0.82 | 0.82 |
| Numeric memory z-score |  |  |  |  |
| N (%) missing<sup>b</sup> | 252 (2.6) | 733 (2.4) | 18 (0.8) | 49 (0.7) |
| Mean (SD) | -0.39 (0.93) | -0.35 (0.94) | -0.24 (0.90) | -0.17 (0.93) |
| Visuospatial memory z-score |  |  |  |  |
| N (%) missing | 1,653 (3.2) | 2,843 (2.8) | 106 (0.9) | 206 (0.8) |
| Mean (SD) | 0.19 (1.05) | 0.26 (1.05) | 0.30 (1.04) | 0.37 (1.03) |
| Prospective memory<sup>b,c</sup> |  |  |  |  |
| N (%) correct | 25,006 (78.0) | 78,673 (76.8) | 7,003 (85.6) | 22,591 (85.7) |

GPS, genome-wide polygenic score; SD, standard deviation.
a. Participants with complete data on all the covariates that were entered into the maximally-adjusted total effects models (age, gender, white British genetic ancestry, English-speaking country of birth, degree, comorbid neurological/psychiatric condition, family history of dementia, family history of Parkinson's disease, family history of severe depression, maternal smoking around birth, childhood trauma, education/cognition GPS, major depression GPS).
b. Missing data refers only to the period when this measure was included in the battery.
c. No missing data.

#### Evaluation of the graphical model

The different predicted independencies implied by alternative plausible specifications of the DAG were tested (see eResults in the Supplement). The best fitting DAG followed the same structure as that used in the mania/BD analysis, with the addition of a path between gender and major depression.

#### Total effects

Major depression was negatively associated with visuospatial memory performance in the multivariable models (Fig. 2). The effect size was very small, with the major depression group scoring approximately 0.07 SD lower than the comparison group. Group differences were not seen on the other cognitive measures (eFigs. 10 to 13 in the Supplement).

**Fig. 2.**
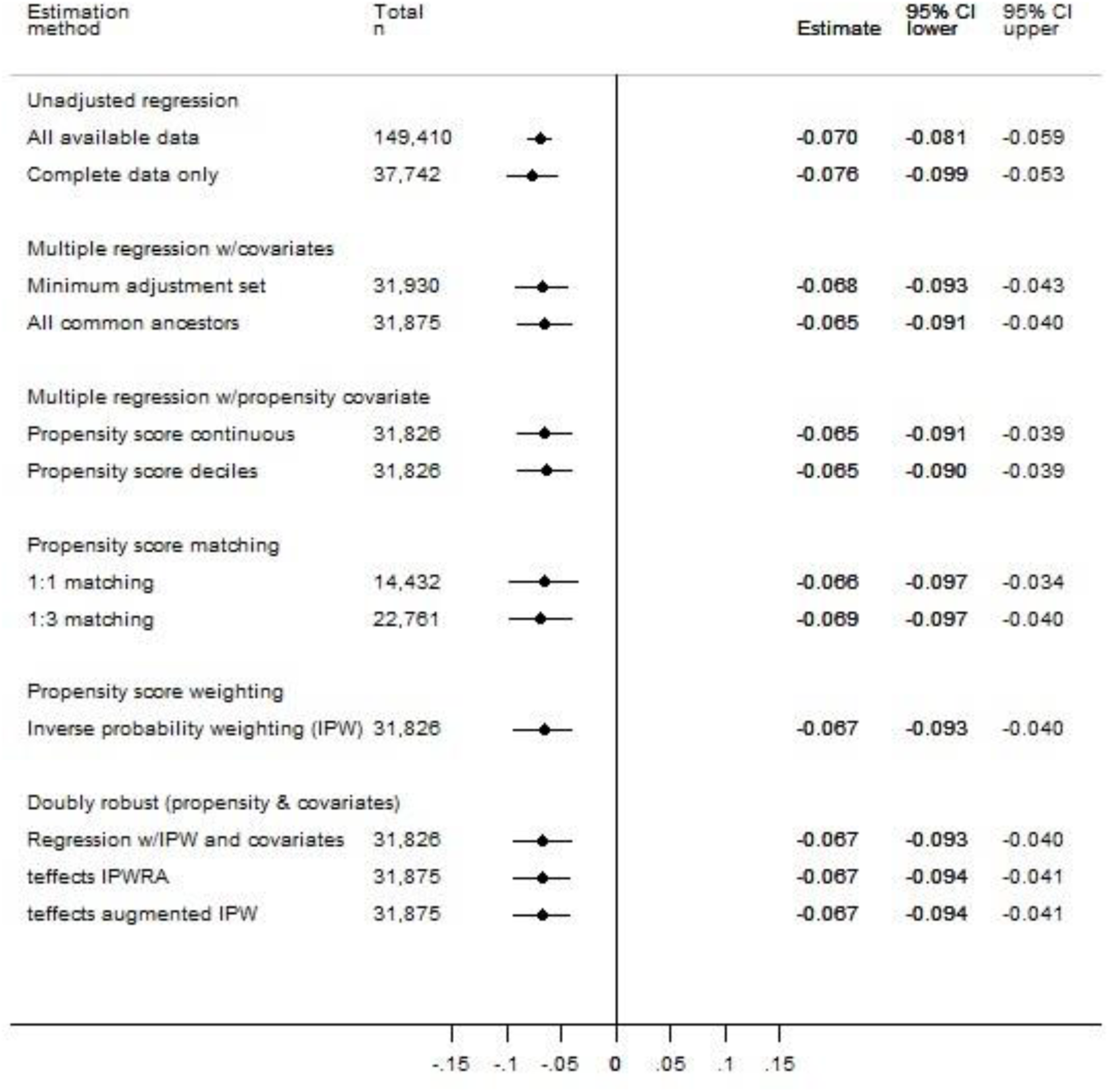
Total Effect of Major Depression on Visuospatial Memory. CI, confidence interval; GPS, genome-wide polygenic score; IPW, inverse probability weighting; IPWRA, inverse probability weighting with regression adjustment; teffects, Stata `teffects` package. Estimates are in z-score units and can be interpreted as standardized mean differences. The minimum sufficient adjustment set comprised gender, educational attainment, English-speaking birth country, ethnicity, education/cognition GPS, major depression GPS, family history of dementia or Parkinson’s disease, maternal smoking around birth, childhood trauma, and comorbid psychiatric/neurological conditions. The extended adjustment set (‘all common ancestors’) also included age and family history of depression. This extended adjustment set was used as the predictor set for the propensity score model. Ethnicity was accounted for in all the multivariable analyses and in the propensity score estimation by restricting these to participants of white British genetic ancestry. The GPS scores were residualized as described in the eMethods, and were entered as deciles, based on the distribution in the full analysis sample.

#### Mediation analyses

As with the mania/BD analyses, direct and indirect effects could be decomposed via cardiometabolic disease and via psychotropic medication (see eResults in the Supplement). There was no evidence of substantive indirect effects via cardiometabolic disease in any of the models (eTable 16 in the Supplement). There was little evidence of mediation via psychotropic medication (eTable 17): approximately one third of the total effect on visuospatial memory (itself of very small magnitude, at −0.058) was estimated to be indirect, but the confidence interval included the null (−0.019; 95% CI −0.040 to 0.003).

#### Sensitivity analyses

The sensitivity analyses (eResults in the Supplement) indicated that: the total effects results for visuospatial memory are likely to be sensitive to exposure misclassification and would not be robust to an unmeasured confounder with even a very weak association with exposure group membership; the total effects results for reaction time showed less attenuation when missing covariate data were imputed; the mediation results with missing covariate data imputation showed stronger evidence for an indirect effect via psychotropic medication but no evidence of an indirect effect via cardiometabolic disease; there were six structurally equivalent DAGs but none were causally plausible.

## Discussion

This study demonstrates small but robust associations between mood disorders and cognitive function in a large community-based sample, with rigorous confounder control based on extensively evaluated graphical models and analytical methods. A total effect of mania/BD on cognitive function was evident on a test of short-term visuospatial memory, but not on other tests. The magnitude of this effect was small, with the point estimates across the various matched/adjusted models being in the range −0.23 to −0.17 standard deviation units. There was evidence of an indirect pathway through psychotropic medication, accounting for approximately one-quarter of the total effect, but (perhaps surprisingly) not through cardiometabolic disease. The total effect of major depression on cognitive performance followed a similar pattern to that of mania/BD, though with an effect size of around one-third the magnitude. The proportion of the total effect mediated by psychotropic medication was similar to that in mania/BD, but the confidence interval was wide and the estimate was reliably different from the null only after imputation of missing data. The effect estimates for both mood disorders are likely to be sensitive to residual confounding and exposure misclassification, and they may be biased toward the null as a result of missing covariate data.

The complexity inherent in this area was acknowledged and addressed by developing and evaluating comprehensive graphical models and by incorporating a broad range of genetic, sociodemographic, environmental, lifestyle and clinical measures in the analyses. Model estimation was conducted in multiple ways, and quantitative and graphical sensitivity analyses were carried out to investigate the robustness of the results to key assumptions. The sample sizes were substantially larger than those used in previous studies in the field, allowing small effect sizes to be estimated with precision.

The observed gradation in severity of impairment on the visuospatial memory task across mania/BD and major depression is congruent with previous reports,^7^ although the magnitude of the difference compared with the non-mood-disorder group is notably smaller.^17-19,22^ The absence of group differences on the other cognitive tasks was surprising, in light of previous research showing multi-domain impairments, and it remains unclear to what extent this reflects insufficient adjustment for confounding in previous studies, the characteristics of the UK Biobank cohort, or the possibility that memory performance is a particularly sensitive marker of cognitive function in mood disorders.

The results also contribute to the evidence base on the relationship between psychotropic medication and cognitive impairment, which has been repeatedly highlighted in previous studies.^32^ Mediation analyses, taking account of intermediate confounders such as past depressive episodes, indicated that an appreciable proportion of the detrimental effect of mania/BD on visuospatial memory performance was accounted for by this. The interplay between reasons for prescribing—especially of antipsychotic medications—and affective remission in understanding this relationship is not yet understood, but the present results appear to confirm that psychotropic medications warrant closer study as potential modifiable causes of cognitive impairment in mood disorders.

### Limitations

The analyses were necessarily limited by the data collected in the UK Biobank resource: many of the measures were brief, key cognitive functions such as verbal memory were not assessed, and clinician diagnoses were unavailable. The reliability of some of the cognitive measures is suboptimal.^14^ This may cause imprecision in effect size estimates,^33^ but such imprecision is mitigated by the large sample size in the present study. However, imprecision that is not due to random error could result in underestimated magnitudes of associations. The cognitive tests administered in UK Biobank may be less sensitive than the neuropsychological assessments used in clinical studies, which may partly account for the small group differences observed here. Assumptions about the temporal order of the variables could not be verified empirically. Conducting estimation one mediator at a time may lead to erroneous conclusions about the contribution of each mediator to the overall effect.^34^ Possible collider stratification bias should be acknowledged;^35^ it is likely that people with less severe mood disorder history and better cognitive function will have joined UK Biobank, and this will have been amplified further in the patterns of missingness across the cognitive outcome measures and the covariates. What is unknown, however, is the magnitude of the bias arising from collider stratification, and how this compares with similar or opposing biases from residual confounding. The missing-at-random assumption that is required for multiple imputation is arguably not valid for some of the measures in these analyses, given the probability that, for example, missingness on mental health-related measures will be influenced by true mental health status. Finally, the UK Biobank cohort is not representative of the UK population; associations may be heterogeneous across other populations.^36^

### Conclusions and Future Directions

A small group difference in visuospatial memory performance was observed between mood disorder and comparison groups in this large general population cohort. Mediation analyses highlighted a potential causal pathway through psychotropic medication use. Our understanding of causal pathways towards cognitive impairment in psychiatric and neurological conditions will improve as the UK Biobank cohort is followed up over time, and other prospective cohorts such as the Avon Longitudinal Study of Parents and Children^37^ mature into adulthood. The availability of a fuller range of background and intermediate data, including early life factors, premorbid cognitive ability measures and brain imaging, will expand the kinds of causal effects that can be identified in models such as those proposed here. Linkage with prescribing data will permit more detailed investigation of the role of different classes of psychotropic medication, and combinations thereof, in explaining adverse cognitive outcomes.

## Author Contributions

Concept and design: All authors.

Acquisition, analysis, or interpretation of data: All authors.

Statistical analysis: B.C.

Drafting of the manuscript: B.C.

Critical revision of the manuscript for important intellectual content: All authors.

Approval of final version: All authors.

## Acknowledgements

This research has been conducted using the UK Biobank Resource under Application Number 11332 (PI: Cullen). UK Biobank was established by the Wellcome Trust medical charity, Medical Research Council, Department of Health, Scottish Government, and the Northwest Regional Development Agency. It has also had funding from the Welsh Government, British Heart Foundation, Cancer Research UK, and Diabetes UK. Data collection was funded by UK Biobank. We are grateful to Dr Carlos Celis Morales, Dr Nicholas Graham, Dr Daniel Mackay, Dr Daniel Martin, Dr Barbara Nicholl, and Mr Joey Ward, University of Glasgow, for contributions to the data analysis.

## Data Availability

The corresponding author had full access to the study data. UK Biobank is an open access resource. Data are available to bona fide scientists, undertaking health-related research that is in the public good. Access procedures are described at http://www.ukbiobank.ac.uk/using-the-resource/.

## References

1. Baune BT, Malhi GS. A review on the impact of cognitive dysfunction on social, occupational, and general functional outcomes in bipolar disorder. Bipolar Disord 2015; 17: 41–55.

2. Evans VC, Iverson GL, Yatham LN, Lam RW. The Relationship Between Neurocognitive and Psychosocial Functioning in Major Depressive Disorder: A Systematic Review. J Clin Psychiat 2014; 75(12): 1359–70.

3. Cullen B, Ward J, Graham NA, et al. Prevalence and correlates of cognitive impairment in euthymic adults with bipolar disorder: A systematic review. J Affect Disord 2016; 205: 165–81.

4. Rock PL, Roiser JP, Riedel WJ, Blackwell AD. Cognitive impairment in depression: a systematic review and meta-analysis. Psychol Med 2014; 44(10): 2029–40.

5. Bourne C, Aydemir O, Balanza-Martinez V, et al. Neuropsychological testing of cognitive impairment in euthymic bipolar disorder: an individual patient data meta-analysis. Acta Psychiatr Scand 2013; 128(3): 149–62.

6. Bortolato B, Miskowiak KW, Koehler CA, Vieta E, Carvalho AF. Cognitive dysfunction in bipolar disorder and schizophrenia: a systematic review of meta-analyses. Neuropsychiatric Disease and Treatment 2015; 11: 3111–25.

7. Szmulewicz AG, Valerio MP, Smith JM, Samame C, Martino DJ, Strejilevich SA. Neuropsychological profiles of major depressive disorder and bipolar disorder during euthymia. A systematic literature review of comparative studies. Psychiatry Research 2017; 248: 127–33.

8. Pearl J, Glymour M, Jewell NP. Causal Inference in Statistics: A Primer. Chichester, West Sussex: Wiley; 2016.

9. VanderWeele TJ. Mediation Analysis: A Practitioner’s Guide. Annu Rev Public Health 2016; 37: 17–32.

10. Sudlow C, Gallacher J, Allen N, et al. UK Biobank: An Open Access Resource for Identifying the Causes of a Wide Range of Complex Diseases of Middle and Old Age. PLoS Medicine 2015; 12(3): e1001779.

11. Smith DJ, Nicholl BI, Cullen B, et al. Prevalence and Characteristics of Probable Major Depression and Bipolar Disorder within UK Biobank: Cross-Sectional Study of 172,751 Participants. PLoS One 2013; 8(11): e75362.

12. Bromet EJ, Kotov R, Fochtmann LJ, et al. Diagnostic Shifts During the Decade Following First Admission for Psychosis. American Journal of Psychiatry 2011; 168(11): 1186–94.

13. Cullen B, Smith DJ, Deary IJ, Evans JJ, Pell JP. The ‘cognitive footprint’ of psychiatric and neurological conditions: cross-sectional study in the UK Biobank cohort. Acta Psychiatr Scand 2017; 135(6): 593–605.

14. Lyall DM, Cullen B, Allerhand M, et al. Cognitive Test Scores in UK Biobank: Data Reduction in 480,416 Participants and Longitudinal Stability in 20,346 Participants. PLoS One 2016; 11(4): e0154222.

15. Strauss E, Sherman EMS, Spreen O. A compendium of neuropsychological tests: Administration, norms, and commentary. 3rd ed. Oxford: Oxford University Press; 2006.

16. Elwert F. Graphical causal models. In: Morgan SL, ed. Handbook of Causal Analysis for Social Research. New York: Springer; 2013: 245–73.

17. Arts B, Jabben N, Krabbendam L, van Os J. Meta-analyses of cognitive functioning in euthymic bipolar patients and their first-degree relatives. Psychol Med 2008; 38(6): 771–85.

18. Bora E, Yucel M, Pantelis C. Cognitive endophenotypes of bipolar disorder: A meta-analysis of neuropsychological deficits in euthymic patients and their first-degree relatives. J Affect Disord 2009; 113(1-2): 1–20.

19. Bora E, Yucel M, Pantelis C, Berk M. Meta-analytic review of neurocognition in bipolar II disorder. Acta Psychiatr Scand 2011; 123(3): 165–74.

20. Camelo EVM, Velasques B, Ribeiro P, Netto T, Cheniaux E. Attention impairment in bipolar disorder: a systematic review. Psychology & Neuroscience 2013; 6(3): 299–309.

21. Lee RSC, Hermens DF, Scott J, et al. A meta-analysis of neuropsychological functioning in first-episode bipolar disorders. J Psychiatr Res 2014; 57: 1–11.

22. Mann-Wrobel MC, Carreno JT, Dickinson D. Meta-analysis of neuropsychological functioning in euthymic bipolar disorder: an update and investigation of moderator variables. Bipolar Disord 2011; 13(4): 334–42.

23. Robinson LJ, Thompson JM, Gallagher P, et al. A meta-analysis of cognitive deficits in euthymic patients with bipolar disorder. J Affect Disord 2006; 93(1-3): 105–15.

24. Samamé C, Martino DJ, Strejilevich SA. An individual task meta-analysis of social cognition in euthymic bipolar disorders. J Affect Disord 2015; 173: 146–53.

25. Kline RB. Principles and Practice of Structural Equation Modeling. 4th ed. New York: The Guilford Press; 2016.

26. Parker G, Brotchie H. Gender differences in depression. International Review of Psychiatry 2010; 22(5): 429–36.

27. Textor J, van der Zander B, Gilthorpe MS, Liskiewicz M, Ellison GTH. Robust causal inference using directed acyclic graphs: the R package ‘dagitty’. International Journal of Epidemiology 2016; 45(6): 1887–94.

28. Pearl J. Fusion, propagation, and structuring in belief networks. Artif Intell 1986; 29(3): 241–88.

29. De Stavola BL, Daniel RM, Ploubidis GB, Micali N. Mediation Analysis With Intermediate Confounding: Structural Equation Modeling Viewed Through the Causal Inference Lens. Am J Epidemiol 2015; 181(1): 64–80.

30. Daniel RM, De Stavola BL, Cousens SN. gformula: Estimating causal effects in the presence of time-varying confounding or mediation using the g-computation formula. Stata J 2011; 11(4): 479–517.

31. von Elm E, Altman DG, Egger M, et al. The Strengthening the Reporting of Observational Studies in Epidemiology (STROBE) statement: guidelines for reporting observational studies. Epidemiology 2007; 18(6): 800–4.

32. Balanzá-Martínez V, Selva G, Martinez-Aran A, et al. Neurocognition in bipolar disorders-A closer look at comorbidities and medications. European Journal of Pharmacology 2010; 626(1): 87–96.

33. Hutcheon JA, Chiolero A, Hanley JA. Random measurement error and regression dilution bias. British Medical Journal 2010; 340: c2289.

34. Vansteelandt S, Daniel RM. Interventional Effects for Mediation Analysis with Multiple Mediators. Epidemiology 2017; 28(2): 258–65.

35. Munafò MR, Tilling K, Taylor AE, Evans DE, Smith GD. Collider scope: when selection bias can substantially influence observed associations. International Journal of Epidemiology 2017; 47(1): 226–35.

36. Fry A, Littlejohns TJ, Sudlow C, et al. Comparison of Sociodemographic and Health-Related Characteristics of UK Biobank Participants with the General Population. Am J Epidemiol 2017; 186(9): 1026–34.

37. Boyd A, Golding J, Macleod J, et al. Cohort Profile: the ‘children of the 90s’--the index offspring of the Avon Longitudinal Study of Parents and Children. International Journal of Epidemiology 2013; 42(1): 111–27.

