## Supplement for "Understanding Cognitive Impairment in Mood Disorders: Mediation Analyses in the UK Biobank Cohort"

### Supplementary Information

### eMethods

#### Exposure Definitions

The mood disorder exposures (mania/BD and major depression) were classified hierarchically into mutually exclusive groups within each information source. Using the self-reported diagnosis data, the hierarchy order was: mania/BD (‘mania/bipolar/manic depression’); major depression (‘depression’ or ‘post-natal depression’). Using the hospital ICD-10 codes, the order was: mania/BD (F30x or F31x); major depression (F32x or F33x). Using the mood questionnaire data, the hierarchy was as described by Smith et al.:[1] mania/BD (BD type I and BD type II combined); major depression (single episode or recurrent). Owing to the limited detail in the self-reported diagnosis data and the mood questionnaire with regard to bipolar features, no distinction was made between single manic episode, bipolar disorder type I and bipolar disorder type II; the term ‘mania/BD’ is therefore used throughout.

#### Covariate Measures

##### Sociodemographic measures

Age was truncated to whole years, and was centred at 55 (approximating the cohort mean age at baseline) in the analyses. Gender was self-reported as male or female. Ethnic background was self-reported as white, Asian/Asian British, black/black British, Chinese, mixed or other. Participants who had self-reported a white British background were further grouped by similarity of genetic ancestry based on a principal components analysis of the genotypic data (http://biobank.ctsu.ox.ac.uk/crystal/field.cgi?id=22006). Participants self-reported their birth country, and these were grouped according to whether or not English was an official/first language (UK, Isle of Man, Channel Islands, Gibraltar, Ireland, Australia, New Zealand, USA, Canada, Anguilla, Antigua & Barbuda, Aruba, Bahamas, Barbados, Bermuda). Self-reported data regarding participants’ highest educational qualification were dichotomized as university/college degree or not. Neighbourhood deprivation level was recorded by UK Biobank prior to baseline using the Townsend Index,[2] and this was converted into quintiles in the whole cohort.

##### Local environment

The population density of each area of residence was classified categorically by UK Biobank, by combining participants’ residential postcodes with data generated from the 2001 census, using the GeoConvert tool provided by the UK Data Service Census Support (http://geoconvert.mimas.ac.uk/). Proximity to the nearest major road (traffic intensity >5,000 motor vehicles per 24 hours) was calculated by UK Biobank as the inverse distance (1/m) from the baseline address, using data for the year 2008 provided by the Department for Transport, and was converted to quintiles in the whole cohort. Neighbourhood air pollution data from a land use regression model and satellite-derived estimates[3] were linked by UK Biobank to participants’ baseline addresses; particulate matter of up to 10μm diameter ($PM_{10}$) and nitrogen dioxide ($NO_2$) were measured as annual average values in $\mu g/m^3$ (for the years 2007 and 2005 respectively) and were converted to quintiles in the whole cohort. Other air pollution data were also available but these were measured in later years, thus post-dating the cognitive assessment date for most participants, and so were not analysed.

##### Lifestyle and physical measures

Tobacco smoking status (current, former or never) was classified by UK Biobank using self-reported data. Self-reported frequency of alcohol consumption was categorized as daily/almost daily, 3-4 times per week, 1-2 times per week, 1-3 times per month, special occasions only, former drinker, or never drinker. Sleeplessness/insomnia was self-reported as never/rarely, sometimes, or usually; if participants were unsure how to respond to this item, they were prompted to answer in relation to the past four weeks. Physical activity (walking, moderate and vigorous) in a typical week was recorded using self-reported items from the International Physical Activity Questionnaire short form,[4] from which a single measure of total physical activity in metabolic equivalent of task (MET) hours per week was derived; this was converted into quintiles in the whole cohort. Body mass index (BMI; $kg/m^2$) was calculated from measures of height and weight taken by UK Biobank staff, and was categorized as underweight (<18.5), normal (18.5 to 24.9), overweight (25.0 to 29.9), obese class I (30.0 to 34.9), obese class II (35.0 to 39.9), and obese class III (≥40.0).

##### Medical and family history

Participants were asked to self-report any illnesses previously diagnosed by a doctor. These data were also combined with hospital records by UK Biobank analysts to generate ‘adjudicated’ classifications of myocardial infarction (http://biobank.ctsu.ox.ac.uk/crystal/docs/alg_outcome_mi.pdf) or stroke (http://biobank.ctsu.ox.ac.uk/crystal/docs/alg_outcome_stroke.pdf). The lists of neurological or psychiatric conditions (apart from mood disorder or schizophrenia) are provided in eTable 1 and eTable 2 below.

#### Mental health and psychotropic medication

Four questions were administered regarding frequency of depressive symptoms in the past two weeks: depressed mood or hopelessness; lack of interest or pleasure; tenseness or restlessness; and tiredness or low energy. These were based on items from the Patient Health Questionnaire.[5] Participants self-rated each symptom on a four-point scale from 'not at all' to 'nearly every day', summed to produce an overall score ranging from 0 to 12, with higher scores indicating more frequent depressive symptoms. Additional self-reported information was elicited using a web-based mental health questionnaire, which was administered in 2016. Information about the number of episodes of depressed mood or anhedonia experienced across the lifetime was collected both at baseline assessment and in the web-based questionnaire; for the present analyses this was coded ordinally (0, 1, 2, 3, 4, 5, 6, 7, 8, 9, 10, >10 and 'too many to count') using the baseline data if available, or the web-based data if the baseline data were missing. These data were available for participants regardless of their mood disorder exposure status (i.e. participants may have reported one or more such episodes without necessarily meeting the criteria for mood disorder). It was not possible to distinguish how many depression episodes preceded the baseline assessment date, in participants who only had web-based data. Participants were also asked five questions from the brief Childhood Trauma Questionnaire[6] within the web-based mental health questionnaire, representing examples of abuse (physical, emotional, sexual) and neglect (physical, emotional). Ordinal responses on each of the five questions were dichotomized using thresholds extrapolated from previous research,[7] and an overall dichotomous indicator was created to represent above-threshold responses on one or more of the five items. The list of psychotropic medications self-reported at baseline is shown in eTable 3 below.

*eTable 1 Psychiatric and neurological diagnoses in hospital records*

| ICD-10 code | ICD-10 description |
|---|---|
| A8x.x | Viral infections of the central nervous system |
| B22.0 | HIV disease resulting in encephalopathy |
| B90.0 | Sequelae of central nervous system tuberculosis |
| B94.1 | Sequelae of viral encephalitis |
| C70.0 | Malignant neoplasm of meninges (cerebral) |
| C71.x | Malignant neoplasm of brain |
| C72.8 | Overlapping lesion of brain and other parts of central nervous system |
| C75.1 | Malignant neoplasm of pituitary gland |
| C75.3 | Malignant neoplasm of pineal gland |
| C79.3 | Secondary malignant neoplasm of brain and cerebral meninges |
| D32.0 | Benign neoplasm of meninges (cerebral) |
| D33.0 | Benign neoplasm of brain, supratentorial |
| D33.1 | Benign neoplasm of brain, infratentorial |
| D33.2 | Benign neoplasm of brain, unspecified |
| D35.2 | Benign neoplasm of pituitary gland |
| D35.4 | Benign neoplasm of pineal gland |
| D42.0 | Neoplasm of uncertain or unknown behaviour of meninges (cerebral) |
| D43.0 | Neoplasm of uncertain or unknown behaviour of brain, supratentorial |
| D43.1 | Neoplasm of uncertain or unknown behaviour of brain, infratentorial |
| D43.2 | Neoplasm of uncertain or unknown behaviour of brain, unspecified |
| D44.3 | Neoplasm of uncertain or unknown behaviour of pituitary gland |
| D44.5 | Neoplasm of uncertain or unknown behaviour of pineal gland |
| Fxx.x | Mental and behavioural disorders |
| G0x.x | Inflammatory diseases of the central nervous system |
| G10 | Huntington disease |
| G11.x | Hereditary ataxia |
| G12.2 | Motor neuron disease |
| G13.1 | Other systemic atrophy primarily affecting central nervous system in neoplastic disease (Paraneoplastic limbic encephalopathy) |
| G2x.x | Extrapyramidal and movement disorders |
| G30.x | Alzheimer disease |
| G31.x | Other degenerative diseases of nervous system, not elsewhere classified |
| G32.8 | Other specified degenerative disorders of nervous system in diseases classified elsewhere |
| G35 | Multiple sclerosis |
| G36.x | Other acute disseminated demyelination |
| G37.x | Other demyelinating diseases of central nervous system |
| G4x.x | Episodic and paroxysmal disorders |
| G8x.x | Cerebral palsy and other paralytic syndromes |
| G90.3 | Multi-system degeneration |
| G91.x | Hydrocephalus |
| G92 | Toxic encephalopathy |
| G93.x | Other disorders of brain |
| G94.x | Other disorders of brain in diseases classified elsewhere |

| ICD-10 code | ICD-10 description |
|---|---|
| G96.x | Other disorders of central nervous system |
| G97.x | Postprocedural disorders of nervous system, not elsewhere classified |
| G98 | Other disorders of nervous system, not elsewhere classified |
| H47.6 | Disorders of visual cortex |
| I6x.x | Cerebrovascular diseases |
| Q0x.x | Congenital malformations of the nervous system |
| Q28.2 | Arteriovenous malformation of cerebral vessels |
| Q28.3 | Other malformations of cerebral vessels |
| Q9x.x | Chromosomal abnormalities, not elsewhere classified |
| R41.x | Other symptoms and signs involving cognitive functions and awareness |
| R90.0 | Intracranial space-occupying lesion |
| R94.0 | Abnormal results of function studies of central nervous system |
| S02.0x | Fracture of vault of skull |
| S02.1x | Fracture of base of skull |
| S06.x | Intracranial injury |
| S07.1 | Crushing injury of skull |
| S09.7 | Multiple injuries of head |
| T02.0x | Fractures involving head with neck |
| T04.0 | Crushing injuries involving head with neck |
| T06.0 | Injuries of brain and cranial nerves with injuries of nerves and spinal cord at neck level |
| T40.x | Poisoning by narcotics and psychodysleptics [hallucinogens] |
| T42.x | Poisoning by antiepileptic, sedative-hypnotic and antiparkinsonism drugs |
| T43.x | Poisoning by psychotropic drugs, not elsewhere classified |
| T51.x | Toxic effect of alcohol |
| T58 | Toxic effect of carbon monoxide |
| T90.2 | Sequelae of fracture of skull and facial bones |
| T90.5 | Sequelae of intracranial injury |

*eTable 2 Self-reported psychiatric and neurological diagnoses*

| UK Biobank data field | Diagnosis |
|---|---|
| 6150 (touchscreen - vascular) | Stroke |
| 20001 (interview - cancer) | Brain cancer/primary malignant tumour |
| " | Meningeal cancer/malignant meningioma |
| 20002 (interview - non-cancer) | Alcohol dependency |
| " | Anorexia/bulimia/other eating disorder |
| " | Anxiety/panic attacks |
| " | Benign/essential tremor |
| " | Brain haemorrhage |
| " | Brain/intracranial abscess |
| " | Cerebral aneurysm |
| " | Cerebral palsy |
| " | Chronic/degenerative neurological problem |
| " | Deliberate self-harm/suicide attempt |
| " | Dementia/Alzheimer's/cognitive impairment |
| " | Encephalitis |
| " | Epilepsy |
| " | Fracture skull/head |
| " | Head injury |
| " | Headaches (not migraine) |
| " | Infection of nervous system |
| " | Insomnia |
| " | Ischaemic stroke |
| " | Meningioma benign |
| " | Meningitis |
| " | Migraine |
| " | Motor neurone disease |
| " | Multiple sclerosis |
| " | Nervous breakdown |
| " | Neurological injury/trauma |
| " | Neuroma benign |
| " | Obsessive compulsive disorder (OCD) |
| " | Opioid dependency |
| " | Other demyelinating condition |
| " | Other neurological problem |
| " | Other substance abuse/dependency |
| " | Parkinson's disease |
| " | Post-traumatic stress disorder |
| " | Psychological/psychiatric problem |
| " | Spina bifida |
| " | Stress |
| " | Stroke |
| " | Subarachnoid haemorrhage |
| " | Subdural haematoma |
| " | Transient ischaemic attack |

*eTable 3* *Self-reported psychotropic medications from UK Biobank data field 20003*

| Mood stabilisers | Selective serotonin reuptake inhibitors | Other antidepressants | Traditional antipsychotics | Second generation antipsychotics | Sedatives & hypnotics |
|---|---|---|---|---|---|
| lithium product | paroxetine | mirtazapine | chlorpromazine | quetiapine | diazepam |
| Priadel (lithium) | Seroxat (paroxetine) | Zispin (mirtazapine) | cpz - chlorpromazine | Seroquel (quetiapine) | diazepam product |
| Camcolit (lithium) | fluoxetine | duloxetine | Largactil (chlorpromazine) | risperidone | Valium tablet (diazepam) |
| sodium valproate | Prozac (fluoxetine) | Cymbalta (duloxetine) | haloperidol | Risperdal (risperidone) | Valium syrup (diazepam) |
| Epilim (sodium valproate) | citalopram | Yentreve (duloxetine) | Haldol (haloperidol) | olanzapine | Valium supp (diazepam) |
| Depakote (semisodium valproate) | Cipramil (citalopram) | venlafaxine | Serenace (haloperidol) | Zyprexa (olanzapine) | temazepam |
| valproic acid | escitalopram | Efexor (venlafaxine) | fluphenazine decanoate | aripiprazole | Normison (temazepam) |
| carbamazepine product | Cipralex (escitalopram) | amitriptyline | fluphenazine | Abilify (aripiprazole) | Euhypnos (temazepam) |
| carbamazepine | sertraline | Elavil (amitriptyline) | Modecate (fluphenazine) | amisulpride | zopiclone |
| Tegretol (carbamazepine) | Lustral (sertraline) | Tryptizol (amitriptyline) | Moditen tablet (fluphenazine) | Solian (amisulpride) | Zimovane (zopiclone) |
| Teril (carbamazepine) | fluvoxamine | Lentizol (amitriptyline) | Moditen enanthate (fluphenazine) | clozapine | zaleplon |
| Teril retard (carbamazepine) | | amitriptyline+perphenazine | flupentixol | Clozaril (clozapine) | Sonata (zaleplon) |
| Timonil retard (carbamazepine) | | Triptafen (amitriptyline+perphenazine) | Flupenthixol (flupentixol) | | zolpidem |
| Epimaz (carbamazepine) | | amitriptyline+chlordiazepoxide | Depixol (flupentixol) | | Stilnoct (zolpidem) |
| lamotrigine | | Limbitrol 10 (amitriptyline+chlordiazepoxide) | Fluanxol (flupentixol) | | nitrazepam |
| Lamictal (lamotrigine) | | Limbitrol-5 (amitriptyline+chlordiazepoxide) | zuclopenthixol | | Mogadon (nitrazepam) |

| Mood stabilisers | Selective serotonin reuptake inhibitors | Other antidepressants | Traditional antipsychotics | Second generation antipsychotics | Sedatives & hypnotics |
|---|---|---|---|---|---|
| | | phenelzine | Clopixol (zuclopenthixol) | | Nitrados (nitrazepam) |
| | | maoi - phenelzine | loxapine | | Remnos (nitrazepam) |
| | | Nardil (phenelzine) | Loxapac (loxapine) | | Somnite (nitrazepam) |
| | | moclobemide | droperidol | | Noctesed (nitrazepam) |
| | | Manerix (moclobemide) | Droleptan (droperidol) | | Surem (nitrazepam) |
| | | imipramine | trifluoperazine | | Unisomnia (nitrazepam) |
| | | Tofranil (imipramine) | Stelazine (trifluoperazine) | | flunitrazepam |
| | | trimipramine | thioridazine | | Rohypnol (flunitrazepam) |
| | | Surmontil (trimipramine) | Melleril (thioridazine) | | triazolam |
| | | dothiepin | | | Halcion (triazolam) |
| | | dosulepin | | | |
| | | Prothiaden (dosulepin) | | | |
| | | Thaden (dosulepin) | | | |
| | | clomipramine | | | |
| | | Anafranil (clomipramine) | | | |
| | | lofepramine | | | |
| | | Gamanil (lofepramine) | | | |
| | | Lomont (lofepramine) | | | |
| | | mianserin | | | |
| | | Bolvidon (mianserin) | | | |
| | | Norval (mianserin) | | | |

#### Genome-wide polygenic scores

Participants provided a blood sample at the baseline assessment, and genotyping was carried out centrally by UK Biobank. Full details of the genotyping, imputation, and quality control processes used by UK Biobank are publicly available at http://biobank.ctsu.ox.ac.uk/crystal/label.cgi?id=100314 and in Bycroft et al.[8] Direct genotyping was performed using two custom Affymetrix arrays: approximately 50,000 participants were genotyped on the UK BiLEVE Axiom array, which was designed for the BiLEVE study of lung function (a partner study of UK Biobank), and the remainder were genotyped using the UK Biobank Axiom array. The two arrays are very similar, with over 95% common marker content. The arrays included more than 800,000 single nucleotide polymorphisms (SNP), chosen because of known or likely associations with a wide range of diseases and health-related phenotypes, as well as to provide good genome-wide coverage for imputation purposes in European populations across common (>5%) and low (1-5%) minor allele frequency (MAF) ranges. The directly genotyped data were imputed by UK Biobank to reference panels from the Haplotype Reference Consortium, UK10K Project and 1000 Genomes Project (Phase 3). Only the Haplotype Reference Consortium imputed data (approximately 40 million markers) were available at the time the present analyses were conducted, due to quality control problems with the UK10K and 1000 Genomes imputations.

The cognitive genome-wide polygenic score (GPS) used summary statistics from a large genome-wide association study (GWAS) of years of education;[9] years of education was used here as a proxy for general cognitive ability because results were unavailable from any similarly-sized GWAS of general cognitive ability that did not involve UK Biobank participants. Education and cognitive ability have a genetic correlation of approximately 0.8,[10] and current evidence suggests that a GPS based on the very large available GWAS of education has greater predictive power for observed cognitive ability than does a GPS based on smaller GWAS of cognitive ability itself.[11] To minimize sample overlap with UK Biobank participants, the education GWAS authors provided summary statistics from analyses that did not include UK cohorts; participants from the 23andMe data resource were also omitted due to data-sharing restrictions. The BD GPS used summary statistics from the Psychiatric Genomics Consortium (PGC);[12] this GWAS included UK cohorts and participant overlap is therefore possible with UK Biobank. The major depression GPS used summary statistics from the most recent PGC GWAS;[13] the GWAS authors provided reanalysed statistics that excluded UK Biobank and 23andMe participants.

The GPS were calculated using a bespoke script for R (https://www.r-project.org/) and PLINK (https://www.cog-genomics.org/plink2) software. They were generated from all available SNPs, applying the following quality control criteria: information score >0.8; Hardy-Weinberg equilibrium test $P > 1\text{x}10^{-6}$; MAF >0.01; linkage disequilibrium clumping $R^2 < 0.1$ using a 250kb window. GPS were created at various thresholds based on the $P$ values in the source GWAS ($5\text{x}10^{-8}$ to 0.9), and were weighted by the GWAS effect sizes at each SNP. The optimum GPS was chosen based on the magnitude of the variance explained ($R^2$) in the relevant phenotype measures in the UK Biobank data. These analyses were conducted in unrelated UK Biobank participants of white British genetic ancestry, after standard quality control exclusions for sex mismatch, sex chromosome aneuploidy, and outlying values of heterozygosity and missingness (http://biobank.ctsu.ox.ac.uk/crystal/field.cgi?id=22027). Each GPS was first regressed on variables indicating the genotyping array and batch, UK Biobank assessment centre, and the first 20 genetic principal components. The residuals from these models were then used as the independent variable in models to predict the relevant UK Biobank phenotype.

For each phenotype, the optimum GPS had a $P$ threshold of 0.5. The $R^2$ for the association between the optimum education/cognition GPS and having a degree in the UK Biobank cohort was 0.018, and was 0.013 for the raw reasoning test score. These results are similar to those previously reported in independent samples, e.g. $R^2 = 0.02$ for both educational attainment[14,15] and general cognitive ability.[16] They are notably lower, however, than more recent analyses which made use of the full education GWAS results including UK participants,[17] in which the $R^2$ for educational attainment at age 16 was 0.091 and for general cognitive ability was 0.036. The $R^2$ for the association between the optimum BD GPS and the mania/BD phenotype in the UK Biobank cohort was 0.01. This is lower than the variance explained in other independent samples, e.g. $R^2 = 0.024$,[18] although results may not be directly comparable due to different methods of calculating pseudo $R^2$ in logistic regression models. The $R^2$ for the optimum major depression GPS was 0.005. This is similar to that reported by the GWAS authors when using their core “anchor” cohort results alone to predict into independent samples, although $R^2$ values of up to 0.02 were obtained when additional cohort results (including UK Biobank and 23andMe) were included in GWAS meta-analyses.[13]

### Statistical Analysis

#### Graphical Models

##### *Construction of the directed acyclic graphs*

The DAG shown in eFigure 1 below represents the original assumptions made about the causal relationship between mania/BD and cognitive function. The presence of an arrow represents the weak assumption of a causal relationship for at least one member of the population, and the absence of an arrow represents the strong assumption of no causal relationship for any member of the population.[19] All nodes that are causally antecedent to a given node are known as ancestors; shared ancestors of the exposure and outcome are potential confounders, because they lie on non-causal 'back-door' paths.[20] Intermediate nodes were included in the DAG where these were of interest in the mediation analyses or were required as common parents of other pairs of nodes; these were not exhaustive, and so each individual arrow could in principle be shown in more detail as a chain of intermediate nodes that, for the purposes of the present study, are omitted or unknown. The node names, the constructs they are intended to represent, and the corresponding measures in UK Biobank, are listed in eTable 4 below.

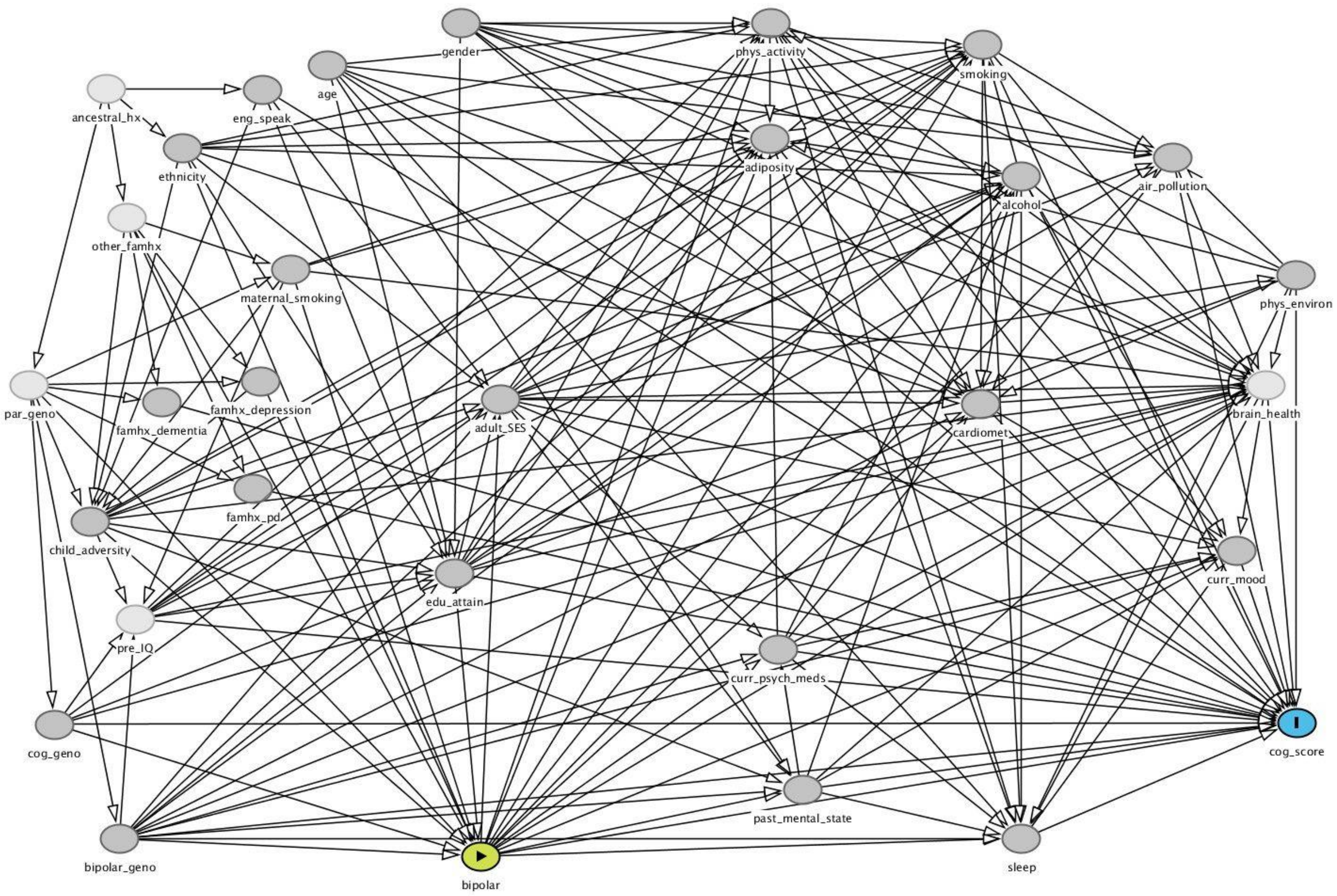

eFigure 1 Original DAG used in the mania/BD analyses

Cardiomet, cardiometabolic disease; cog, cognitive; curr, current; edu, educational; eng_speak, English speaking birth country; famhx, family history; geno, genotype; hx, history; par, parental; PD, Parkinson's disease; phys, physical; pre_IQ, premorbid intelligence; psych_meds, psychotropic medications; SES, socioeconomic status. Green node is the exposure and blue node is the outcome. Light nodes represent unmeasured constructs and darker nodes represent measured constructs.

eTable 4 DAG constructs

| Node name | Construct represented | Measurement in UK Biobank |
|---|---|---|
| **Exposure** | | |
| bipolar | Lifetime history of mania or bipolar disorder (prior to cognitive assessment) | Disorder exposure status (versus comparison group) |
| **Outcome** | | |
| cog_score | Performance on cognitive assessment | • Reasoning<br>• Reaction time<br>• Numeric memory<br>• Visuospatial memory<br>• Prospective memory |
| **Shared ancestors of exposure and outcome** | | |
| age | Age | Age in years |
| ancestral_hx | Various ancestral/migration factors that determine ethnicity, country of origin and family history (genetic and non-genetic) | *Unmeasured* |
| bipolar_geno | Genotype associated with bipolar disorder | Genome-wide polygenic score |
| child_adversity | Adverse experiences in childhood | Childhood abuse and neglect (self-reported as 'never true' to 'very often true'):<br>When I was growing up…<br>a) I felt loved<br>b) People in my family hit me so hard that it left me with bruises or marks<br>c) I felt that someone in my family hated me<br>d) Someone molested me (sexually)<br>e) There was someone to take me to the doctor if I needed it |
| cog_geno | Genotype associated with cognitive function | Genome-wide polygenic score (using education GWAS as proxy) |
| edu_attain | Educational attainment | Has a degree or not (self-reported) |
| eng_speak | Born in an English-speaking country | Born in an English-speaking country (self-reported) |
| ethnicity | Ethnic background | • Ethnic category (self-reported)<br>• Genetically-identified white British ancestry |
| famhx_depression | Parent/sibling with depression | Self-report of biological parent or sibling with 'severe depression' |
| gender | Gender | Self-reported male or female |
| maternal_smoking | Mother smoked around time of participant's birth | Participant's response to "Did your mother smoke regularly around the time when you were born?" |
| other_famhx | Other aspects of family history (non-genetic) and circumstances/environment | *Unmeasured* |
| par_geno | Genotype of parents | *Unmeasured* |
| pre_IQ | Premorbid intellectual ability | *Unmeasured* |
| **Intermediates between exposure and outcome** | | |
| adiposity | Body fat | Body mass index |
| adult_SES | Socioeconomic status or deprivation in adulthood | Townsend index score |

| Node name | Construct represented | Measurement in UK Biobank |
|---|---|---|
| air_pollution | Airborne toxic particles/gases | Neighbourhood measures of:<br>• Particulate matter<br>• Nitrogen dioxide |
| alcohol | Frequency/amount of alcohol consumption | Self-reported frequency of intake |
| brain_health | Structural/functional brain state | *Unmeasured (except for small subgroup)* |
| cardiomet | History of cardiometabolic disease | • Self-reported history of angina, hypertension or diabetes (non-gestational)<br>• Adjudicated history of myocardial infarction or stroke |
| curr_mood | Mood state at time of cognitive assessment | Patient Health Questionnaire (four self-reported items) |
| curr_psych_meds | Psychotropic medication at time of cognitive assessment | On any psychotropic medication (self-reported) |
| past_mental_state | Past psychiatric symptoms/illness course/duration/severity, over and above history of simply having exposure of interest or not | Number of depressed/unenthusiastic episodes (self-reported on touchscreen or web) |
| phys_activity | Level of physical activity | International Physical Activity Questionnaire (self-reported) |
| phys_environ | Physical aspects of the local environment | • Inverse distance to nearest major road<br>• Home area population density |
| sleep | Sleep pattern/quality/duration | Self-reported sleeplessness/insomnia (never/rarely; sometimes; usually) |
| smoking | Tobacco smoking history | Self-reported smoking status (never; former; current) |
| **Other ancestors of outcome (not descended from exposure)** | | |
| famhx_dementia | Parent/sibling with dementia | Self-report of biological parent or sibling with 'Alzheimer's/dementia' |
| famhx_pd | Parent/sibling with Parkinson's disease | Self-report of biological parent or sibling with 'Parkinson's disease' |

#### Shared ancestors of exposure and outcome

Older age increases the risk of cognitive impairment, although the trajectory and mechanisms are not fully understood.[21] Age was also assumed to have an indirect effect on mania/BD status, via educational attainment. Gender differences in average cognitive performance have often been reported, although again the causal mechanisms are not well understood.[22] Gender was assumed not to affect mania/BD status,[23,24] except through educational attainment. The direction of the relationship between educational attainment and mania/BD was uncertain, and the model was tested with the arrow as shown above and with it reversed. Given the temporal order of the measures in UK Biobank, educational attainment (past) was assumed to influence cognitive performance (current). Premorbid intellectual ability was not measured in UK Biobank, but was depicted as an antecedent of current cognitive performance and of other nodes such as educational attainment, socioeconomic status and health-related behaviours (e.g. smoking). No arrow was drawn between premorbid ability and mania/BD status, because it was assumed that any statistical association between them would be accounted for by their shared genetic and early life antecedents (see below), or by the indirect causal path through educational attainment.

Genotypes associated with bipolar disorder and with cognitive function were assumed to have shared effects on those respective phenotypes and on other outcomes.[25,26] These genotypes were depicted as descending from parental genotype (unmeasured). Parental genotype and other aspects of family history (unmeasured) were also assumed to affect parental behaviour (maternal smoking measure), childhood adversity, and family history of psychiatric and neurological conditions. A distal node representing ancestral history was conceptualised as giving rise to parental genotype and other aspects of family history, as well as ethnicity and English-speaking status. Ethnicity was assumed to be a possible antecedent of mania/BD,[24] and (through other nodes such as socioeconomic status) of cognitive performance. Being from a non-English-speaking country was assumed to influence educational attainment and childhood adversity (e.g. among individuals who had migrated at a young age), in turn influencing mania/BD status, and it was also assumed to affect cognitive performance.

Maternal smoking was assumed to affect mania/BD,[27] and to affect cognitive function indirectly through other nodes (e.g. brain health). Childhood adversity (conceptualised broadly when constructing the graph, although measured in UK Biobank solely as abuse and neglect history) was assumed to be a possible cause of mania/BD,[28] and of cognitive function via nodes such as educational attainment, socioeconomic status and health-related behaviours. Finally, family history of depression was depicted as a cause of mania/BD,[29] and of cognitive function via childhood adversity.

#### Intermediates between exposure and outcome

Lifetime history of mania/BD was assumed to affect multiple behaviour-related measures, namely physical activity, adiposity, alcohol consumption and smoking,[30] and these in turn were assumed to influence cognitive function,[31] including via their effects on brain structure. Cardiometabolic disease was also assumed to be influenced by mania/BD status,[32,33] and to affect cognitive outcome.[34,35] It was assumed that mania/BD might affect socioeconomic status in adulthood (e.g. via impact on occupational functioning);[36] this in turn might influence performance on cognitive tests,[37] although this relationship is complicated by shared genetic factors,[38] as shown in the graph. Note that early life socioeconomic status was not measured in UK Biobank, but this can be conceptualised as part of the childhood adversity node, and can thus be assumed to influence both mania/BD and adult socioeconomic status. With regard to local environment variables, mania/BD was assumed to influence exposure to air pollution, including via socioeconomic status, smoking and physical activity; the relationship between mania/BD and other aspects of the physical environment was assumed to arise indirectly via socioeconomic status. Physical environment exposures, including pollution, were assumed to affect cognitive function.[39,40]

It was assumed that mood state around the time of the cognitive assessment would be influenced by mania/BD status, and would in turn affect cognitive performance.[41,42] Mood state was here measured by items assessing current depressive symptoms only; manic mood state (unmeasured in UK Biobank) was not included as a separate node in the graph, as previous research had reported no association between residual mania and cognitive performance.[41] Sleeplessness was also depicted as an intermediate between mania/BD status and cognitive performance. Although sleep disturbance can be a trigger for relapse in BD, it was placed temporally downstream of mania/BD status in this graph because it represented recent sleep patterns (in the four-week period preceding the UK Biobank cognitive assessment), whereas the mania/BD node represented lifetime status. Sleeplessness is an ongoing problem for many people with BD,[43] and it may affect cognitive performance.[44,45]

A node representing past mental state was included as an intermediate between mania/BD status and cognitive score. Although the past mental state node and the mania/BD exposure node both represent past states (i.e. lifetime experiences up to the time of the cognitive assessment), mania/BD was placed first in the temporal order depicted in the graph, on the grounds that having mania/BD influences the severity of the illness experience over time (measured here as number of depressed episodes). It was not a requirement in the exposure definition used here that a participant had to have experienced more than one affective episode to be classified in the mania/BD group, and so it was deemed more plausible that mania/BD status would influence the number of episodes, rather than the reverse. Similarly, current psychotropic medications at the time of the cognitive assessment were assumed to be a consequence of lifetime mania/BD status, and being on such medications did not contribute to the exposure classification used here. The temporal order of the past mental state and current psychotropic medication nodes was depicted such that the former influenced the latter, although it is likely that there is a reciprocal relationship between these variables over time (i.e. a graph for a longitudinal analysis might show mental state at time 1 influencing medication at time 2, and medication at time 2 influencing mental state at time 3). Given the nature of the present cross-sectional analysis, however, it was considered plausible that cumulative lifetime affective episodes (past mental state) would be an antecedent of current medication status. Both past mental state and current psychotropic medications were assumed to affect cognitive performance.[41,46,47]

A node representing brain health (i.e. as potentially measured by structural volume and integrity, and functional activation and connectivity) was depicted as intermediate between mania/BD status and cognitive performance. This assumes that changes occur in the brain as a consequence of mania/BD, as shown in longitudinal studies,[48] although it may also be the case that some brain changes (e.g. reduced white matter tract integrity) are a marker of BD vulnerability that precedes illness onset, given that similar findings are evident in unaffected relatives of people with BD.[49] Structural and functional brain changes were assumed to cause cognitive impairment, although—as shown in the graph—premorbid cognitive ability and shared genetic antecedents likely contribute to this relationship.[21] It was not assumed that every causal path leading to cognitive outcome went through brain health, however, as other factors may be at play (such as confidence or test experience) that would affect performance on cognitive tests but are not necessarily mediated by brain structure or function. Since neuroimaging data were available only for a relatively small and non-representative sub-group (~2%) of the UK Biobank cohort, this node was tagged as unmeasured when planning the present analyses.

#### Other ancestors of outcome

Family history of neurodegenerative disease was assumed to be a potential additional cause of cognitive impairment, given that participants with this background may be at higher risk of cognitive decline arising from disease processes not necessarily captured by their own medical history data (e.g. individuals with unrecognised early-stage disease). This was represented in the graph by separate nodes for family history of dementia and of Parkinson's disease, although this could have been depicted equivalently using one node, because they were considered to share the same antecedents and consequences. Family history of neurodegenerative disease was assumed not to be a causal antecedent of mania/BD status.

##### *Testing the fit of the directed acyclic graphs*

The DAG was drawn using DAGitty software,[50] which automatically generated a list of all the testable independencies implied by its structure (ignoring any nodes that were tagged as being unmeasured). These were then tested in the dataset, by calculating partial correlation coefficients between each pair of nodes that were predicted to be independent, adjusting for other covariates if this was specified in the prediction. For example, a predicted conditional independency generated by DAGitty such as

age ⊥ physical_environment | deprivation educational_attainment

would be tested by calculating the partial correlation coefficient between age and a measure of the local physical environment, adjusted for the Townsend deprivation score and having a degree. These calculations were done using correlation or regression models, depending on the need to adjust for covariates. For simplicity, only continuous or dichotomous measures were used in the initial calculations. Where a node had more than one relevant available measure (e.g. physical_environment measured by population density or road proximity), the measure with the largest sample size was used, in order to minimize missing data bias.

#### Total Effects

For the purpose of comparison, estimation was conducted in several ways:

- Unadjusted regression model in all available participants;
- Unadjusted regression model only in participants who had complete data on all covariates that were to be used in the adjusted models;
- Multiple regression model adjusted for the minimum sufficient covariate set identified by DAGitty;
- Multiple regression model adjusted for the minimum sufficient set plus all other measured common antecedents of exposure and outcome;
- Multiple regression model adjusted for a propensity score created by regressing the mood disorder exposure on background covariates;
- Matched analyses (1:1 and 1:3) using the propensity score to form matched participant sets;
- Weighted regression model using inverse probability weights (IPW) derived from the propensity score;
- Doubly robust models (IPW-weighted regression with additional covariate adjustment or augmented weighting).

Where models included age as a covariate, age squared was also entered, to account for possible curvilinear relationships. The propensity score model was specified in three ways, and the score that resulted in the best covariate balance (evaluated by comparing descriptive statistics for each covariate between the propensity score-matched samples) was taken forward into the total effects analyses listed above. This decision was based solely on covariate balance, without reference to the cognitive outcome data. The first propensity score model regressed the mood disorder exposure status variable on all ancestors of the exposure and all ancestors of the outcome that were not descended from the exposure.[51] The second propensity score model used the same predictor variables as the first, but also included all pairwise interaction terms. The third approach used boosted regression modelling (a machine learning method) to find the optimum prediction specification,[52] again using the same predictor variables as the other two models. The propensity score was also converted to an inverse probability weight, which was rescaled to sum to 1.[53]

The analyses were conducted using Stata v15.[54] The propensity scores were estimated using `psmatch2` and `boost`. The total effects models were estimated using `regress` or `logistic`, `psmatch2` and `teffects`, and results were reported as standardized mean differences with 95% confidence intervals (CI) calculated from robust standard errors. For the logistic regression models (prospective memory outcome measure), `adjrr` was used to convert the odds ratio (OR) estimates into risk differences. The matched models were performed with replacement and used a caliper set at 0.2 SD of the logit of the propensity score.[55]

##### Mediation Analyses

The covariate adjustment sets in these models were the minimum sufficient adjustment sets to block all confounding paths, as determined by DAGitty. All outcome and intermediate confounder variables were entered in continuous or binary form, as required by `gformula`.

#### Sensitivity Analyses

##### Conditional Exchangeability

The assumption of conditional exchangeability implies that, within a matched pair of participants, each participant had equal odds of being exposed ('treated') and unexposed. Covariate balance checks allow this to be verified with respect to measured background factors, but cannot confirm that matched pairs are balanced (i.e. exchangeable) for unmeasured or unknown background variables. Sensitivity of treatment effect estimates or their *P* values to different potential magnitudes of departure from exchangeability can be evaluated quantitatively using 'bounds' methods developed by Rosenbaum.[56] Potential deviations from exchangeability are summarized in a parameter referred to as gamma (where $\Gamma = 1$ represents equal odds), and the value of gamma at which the effect estimate or *P* value crosses the null is ascertained using permutation methods. This was conducted following the propensity score matched models, using the Stata package `rbounds`[a].

##### Missing Data

Because the estimation methods used here involved adjustment and/or propensity score estimation for a large number of covariates, results were potentially sensitive to selection bias or reduced power, arising from missing data. Multiple imputation with chained equations was implemented using the `ice` package in Stata, and the regression models for total effects were repeated on the imputed datasets (25 imputations) using the `mi estimate` function. The cognitive outcome variables were included in the imputation model specification,[57] but their original (unimputed) values were analysed in the outcome models. A chained equations imputation option was also implemented in `gformula`, to allow a comparison of the results of the mediation models using raw versus imputed mediator and covariate data. These methods assume missingness-at-random.

##### Exposure Misclassification

The effect of different hypothetical levels of exposure misclassification on cognitive outcome was assessed using the Stata package `episens`. The outcome was dichotomized as impaired (z-score ≤ -1.645, i.e. 5$^{th}$ percentile) or not. The range of assumed sensitivity and specificity values for correct exposure classification was entered as a trapezoidal function, by specifying minimum and maximum values around a narrower range of equally probable values (e.g. minimum 0.7 and maximum 1.0, around a peak interval of 0.8 to 0.9). Differential misclassification was assumed, on the grounds that cognitively impaired participants would be more likely to be misclassified on the self-reported exposure data.

##### Equivalent Models

The final DAG that was used as the basis for the total effects and mediation models was analysed structurally in the DAGitty R package, to determine the number of alternative ways it could be drawn while retaining the same predicted

---

[a] The `rbounds` package conducts sensitivity analyses on the 'average treatment effect in the treated' (ATT). The ATT represents the average effect of treatment/exposure on outcome within the group that actually received the treatment/exposure.

conditional independencies. This ascertains whether there are other specifications of the model that would be statistically indistinguishable from the version that was analysed, i.e. an 'equivalence class'.[50]

### eResults

#### Cognitive Impairment in Mania/BD

##### Characteristics of the Sample

eFigure 2 below shows a flowchart of exclusions leading to the final analysis sample. A large number of participants were excluded due to missing data in at least one exposure information source, which meant they could not be classified in the comparison group. Where genotyping data indicated relatedness (third degree or closer), one member of each related set was chosen at random for analysis. Ethnic ancestry exclusions were applied only in the adjusted models.

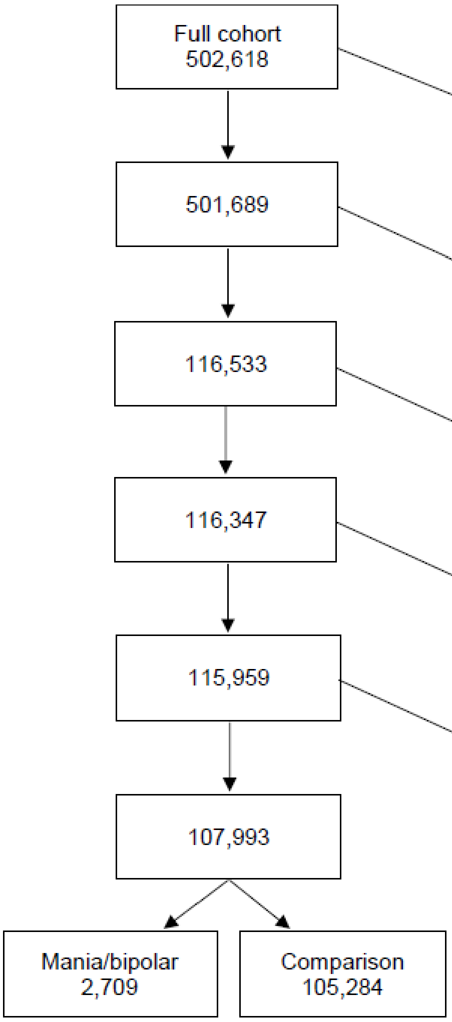

*eFigure 2 Mania/bipolar disorder analysis sample flowchart*

eTable 5 summarizes the covariate data in the mania/BD and comparison groups. Missingness was more common in the mania/BD group, apart from on the number of depressed episodes. Owing to the low response rate on the web-based mental health questionnaire (N = 157,366; 31.3% of the whole cohort), the proportion of missing data was highest for the childhood trauma variable. Missingness was also common on the family medical history, maternal smoking, current depressive symptoms and physical activity variables. eTable 5 indicates that the mania/BD group was younger on average than the comparison group, and had a higher proportion of women and of degree-holders. The participants with mania/BD also appeared to be more likely to live in urban and more deprived areas, and to be current smokers and former drinkers. The proportions with frequent sleeplessness, obesity, cardiometabolic disease, comorbid neurological/psychiatric conditions, family history of severe depression, current psychotropic medication, and history of childhood trauma were higher in the mania/BD group, and this group also reported more depressed episodes and a higher current depressive symptom score on average. The distribution of the education/cognition GPS score appeared to be somewhat different between the mania/BD and comparison groups, with both low (decile 1) and high (decile 10) GPS values being slightly over-represented in the mania/BD group. The distribution of the bipolar GPS score was skewed towards higher values in the mania/BD group. The subset of participants with complete covariate data appeared different from the full analysis sample, being on average younger, more highly educated and from less deprived areas, for example.

*eTable 5 Summary of covariates in the mania/bipolar and comparison groups*

| | All available data | | Complete covariate data[a] | |
|---|---|---|---|---|
| | **Mania/BD** | **Comparison** | **Mania/BD** | **Comparison** |
| N | 2,709 | 105,284 | 504 | 26,997 |
| **Sociodemographic** | | | | |
| Age (years)[b] | | | | |
| Mean (SD) | 55.0 (8.1) | 57.0 (8.2) | 54.3 (7.5) | 56.3 (7.9) |
| Gender[b] | | | | |
| N (%) female | 1,437 (53.1) | 52,730 (50.1) | 277 (55.0) | 14,414 (53.4) |
| Ethnic group | | | | |
| N (%) missing | 18 (0.7) | 411 (0.4) | 0 (0.0) | 73 (0.3) |
| White, N (%)[c] | 2,457 (91.3) | 95,463 (91.0) | 479 (95.0) | 25,708 (95.5) |
| Asian/Asian British | 74 (2.8) | 3,697 (3.5) | 16 (3.2) | 459 (1.7) |
| Black/Black British | 78 (2.9) | 3,182 (3.0) | 2 (0.4) | 322 (1.2) |
| Chinese | 4 (0.2) | 474 (0.5) | 1 (0.2) | 99 (0.4) |
| Mixed & other background | 78 (2.9) | 2,057 (2.0) | 6 (1.2) | 336 (1.3) |
| White British genetic ancestry | | | | |
| N (%) missing | 104 (3.8) | 3,419 (3.3) | 0 (0.0) | 0 (0.0) |
| N (%)[c] | 1,968 (75.6) | 81,183 (79.7) | 394 (78.2) | 22,606 (83.7) |
| English-speaking country of birth | | | | |
| N (%) missing | 6 (0.2) | 149 (0.1) | 0 (0.0) | 0 (0.0) |
| N (%)[c] | 2,432 (90.0) | 93,802 (89.2) | 462 (91.7) | 24,993 (92.6) |
| Has a degree | | | | |
| N (%) missing | 29 (1.1) | 1,045 (1.0) | 0 (0.0) | 0 (0.0) |
| N (%)[c] | 1,034 (38.6) | 36,797 (35.3) | 265 (52.6) | 13,210 (48.9) |
| Townsend quintile[d] | | | | |
| N (%) missing | 3 (0.1) | 162 (0.2) | 0 (0.0) | 34 (0.1) |
| Qu1 (least deprived), N (%)[c] | 334 (12.3) | 18,097 (17.2) | 78 (15.5) | 5,162 (19.1) |
| Qu2 | 361 (13.3) | 21,344 (20.3) | 70 (13.9) | 5,846 (21.7) |
| Qu3 | 461 (17.0) | 21,794 (20.7) | 93 (18.5) | 5,924 (22.0) |
| Qu4 | 607 (22.4) | 23,685 (22.5) | 124 (24.6) | 6,094 (22.6) |
| Qu5 (most deprived) | 943 (34.9) | 20,202 (19.2) | 139 (27.6) | 3,937 (14.6) |
| **Local environment** | | | | |
| Home area population density[e] | | | | |
| N (%) missing | 38 (1.4) | 968 (0.9) | 5 (1.0) | 271 (1.0) |
| England/Wales urban, N (%)[c] | 2,311 (86.5) | 90,610 (86.9) | 433 (86.8) | 22,766 (85.2) |
| England/Wales town | 129 (4.8) | 6,603 (6.3) | 25 (5.0) | 1,776 (6.7) |
| England/Wales village | 88 (3.3) | 4,967 (4.8) | 20 (4.0) | 1,533 (5.7) |
| England/Wales hamlet/isolated | 45 (1.7) | 2,136 (2.1) | 10 (2.0) | 651 (2.4) |
| Scotland large urban | 80 (3.0) | 0 (0.0) | 9 (1.8) | 0 (0.0) |
| Scotland other urban | 11 (0.4) | 0 (0.0) | 1 (0.2) | 0 (0.0) |
| Scotland small town | 5 (0.2) | 0 (0.0) | 1 (0.2) | 0 (0.0) |
| Scotland rural | 2 (0.1) | 0 (0.0) | 0 (0.00 | 0 (0.0) |
| Proximity to major road (1/m) | | | | |
| N (%) missing | 65 (2.4) | 1,299 (1.2) | 9 (1.8) | 348 (1.3) |
| Mean (SD) | 0.006 (0.021) | 0.006 (0.013) | 0.006 (0.010) | 0.005 (0.011) |
| Particulate matter ≤10μm (μg/m$^3$) | | | | |
| N (%) missing | 72 (2.7) | 1,679 (1.6) | 11 (2.2) | 437 (1.6) |
| Mean (SD) | 22.9 (3.1) | 22.8 (3.0) | 23.2 (3.2) | 22.8 (3.2) |

| | All available data | | Complete covariate data[a] | |
|---|---|---|---|---|
| | Mania/BD | Comparison | Mania/BD | Comparison |
| Nitrogen dioxide (µg/m$^3$) | | | | |
| N (%) missing | 65 (2.4) | 1,299 (1.2) | 9 (1.8) | 348 (1.3) |
| Mean (SD) | 32.9 (11.0) | 31.9 (10.6) | 33.1 (11.7) | 31.8 (11.0) |
| **Lifestyle and physical** | | | | |
| Smoking status | | | | |
| N (%) missing | 15 (0.6) | 380 (0.4) | 1 (0.2) | 47 (0.2) |
| Never, N (%)[c] | 1,185 (44.0) | 60,305 (57.5) | 251 (49.9) | 16,428 (61.0) |
| Former | 922 (34.2) | 35,436 (33.8) | 175 (34.8) | 8,885 (33.0) |
| Current | 587 (21.8) | 9,163 (8.7) | 77 (15.3) | 1,637 (6.1) |
| Alcohol frequency | | | | |
| N (%) missing | 12 (0.4) | 80 (0.1) | 1 (0.2) | 4 (0.01) |
| Daily/almost daily, N (%)[c] | 497 (18.4) | 22,179 (21.1) | 112 (22.3) | 6,514 (24.1) |
| 3-4 times per week | 451 (16.7) | 24,718 (23.5) | 98 (19.5) | 7,176 (26.6) |
| 1-2 times per week | 583 (21.6) | 26,774 (25.5) | 114 (22.7) | 6,627 (24.6) |
| 1-3 times per month | 312 (11.6) | 11,397 (10.8) | 63 (12.5) | 2,872 (10.6) |
| Special occasions only | 432 (16.0) | 11,824 (11.2) | 66 (13.1) | 2,387 (8.8) |
| Never (former drinker) | 261 (9.7) | 3,186 (3.0) | 36 (7.2) | 620 (2.3) |
| Never (not former drinker) | 161 (6.0) | 5,126 (4.9) | 14 (2.8) | 797 (3.0) |
| Sleeplessness | | | | |
| N (%) missing | 2 (0.1) | 79 (0.1) | 0 (0.0) | 14 (0.1) |
| Never/rarely, N (%)[c] | 535 (19.8) | 29,205 (27.8) | 113 (22.4) | 8, 019 (29.7) |
| Sometimes | 1,199 (44.3) | 50,293 (47.8) | 211 (41.9) | 12,802 (47.4) |
| Usually | 973 (35.9) | 25,707 (24.4) | 180 (35.7) | 6,162 (22.8) |
| Physical activity (MET h/week) | | | | |
| N (%) missing | 256 (9.5) | 6,861 (6.5) | 31 (6.2) | 1,054 (3.9) |
| Median (Q1, Q3) | 25.6 (11.6, 56.7) | 29.8 (13.7, 60.1) | 27.5 (12.3, 54.9) | 29.1 (14.2, 55.7) |
| Body mass index | | | | |
| N (%) missing | 27 (1.0) | 741 (0.7) | 1 (0.2) | 90 (0.3) |
| Underweight, N (%)[c] | 16 (0.6) | 514 (0.5) | 3 (0.6) | 156 (0.6) |
| Normal | 719 (26.8) | 34,893 (33.4) | 174 (34.6) | 10,751 (40.0) |
| Overweight | 1,043 (38.9) | 44,979 (43.0) | 210 (41.8) | 11,104 (41.3) |
| Obese class I | 610 (22.7) | 17,804 (17.0) | 87 (17.3) | 3,750 (13.9) |
| Obese class II | 205 (7.6) | 4,729 (4.5) | 20 (4.0) | 860 (3.2) |
| Obese class III | 89 (3.3) | 1,624 (1.6) | 9 (1.8) | 286 (1.1) |
| **Medical and family history** | | | | |
| Cardiometabolic disease | | | | |
| N (%) missing | 11 (0.4) | 178 (0.2) | 0 (0.0) | 22 (0.1) |
| N (%)[c] | 986 (36.6) | 32,120 (30.6) | 140 (27.8) | 6,310 (23.4) |
| Comorbid neurological or psychiatric condition[f] | | | | |
| N (%) | 814 (30.1) | 9,430 (9.0) | 120 (23.8) | 2,093 (7.8) |
| Family history of dementia | | | | |
| N (%) missing | 473 (17.5) | 14,799 (14.1) | 0 (0.0) | 0 (0.0) |
| N (%)[c] | 384 (17.2) | 15,755 (17.4) | 83 (16.5) | 4,661 (17.3) |
| Family history of Parkinson's disease | | | | |
| N (%) missing | 589 (21.8) | 16,406 (15.6) | 0 (0.0) | 0 (0.0) |
| N (%)[c] | 107 (5.1) | 4,250 (4.8) | 18 (3.6) | 1,276 (4.7) |

| | All available data | | Complete covariate data[a] | |
|---|---|---|---|---|
| | Mania/BD | Comparison | Mania/BD | Comparison |
| Family history of severe depression | | | | |
| N (%) missing | 466 (17.2) | 15,655 (14.9) | 0 (0.0) | 0 (0.0) |
| N (%)[c] | 874 (39.0) | 10,503 (11.7) | 170 (33.7) | 3,086 (11.4) |
| Maternal smoking around birth | | | | |
| N (%) missing | 390 (14.4) | 13,420 (12.7) | 0 (0.0) | 0 (0.0) |
| N (%)[c] | 716 (30.9) | 24,807 (27.0) | 151 (30.0) | 7,065 (26.2) |
| **Mental health** | | | | |
| Current depressive symptoms | | | | |
| N (%) missing | 265 (9.8) | 8,758 (8.3) | 27 (5.4) | 1,223 (4.5) |
| Mean (SD) | 3.5 (3.2) | 1.2 (1.7) | 2.8 (2.9) | 1.0 (1.4) |
| Any psychotropic medication | | | | |
| N (%) missing | 38 (1.4) | 1,247 (1.2) | 5 (1.0) | 286 (1.1) |
| N (%)[c] | 1,457 (54.6) | 2,647 (2.5) | 221 (44.3) | 503 (1.9) |
| Number of depressed episodes | | | | |
| N (%) missing | 127 (4.7) | 6,573 (6.2) | 23 (4.6) | 1,448 (5.4) |
| Median (Q1, Q3) | 1 (0, 6) | 0 (0, 1) | 4 (2, 11) | 0 (0, 1) |
| Any childhood trauma[g] | | | | |
| N (%) missing | 1,976 (72.9) | 68,715 (65.3) | 0 (0.0) | 0 (0.0) |
| N (%)[c] | 476 (64.9) | 16,015 (43.8) | 317 (62.9) | 11,240 (41.6) |
| **Genome-wide polygenic scores** | | | | |
| Education/cognition GPS decile[d] | | | | |
| N (%) missing | 104 (3.8) | 3,419 (3.3) | 0 (0.0) | 0 (0.0) |
| D1 (lowest), N (%)[c] | 291 (11.2) | 10,156 (10.0) | 36 (7.14) | 2,262 (8.4) |
| D2 | 259 (9.9) | 10,188 (10.0) | 57 (11.3) | 2,439 (9.0) |
| D3 | 281 (10.8) | 10,166 (10.0) | 51 (10.1) | 2,497 (9.3) |
| D4 | 234 (9.0) | 10,213 (10.0) | 55 (10.9) | 2,619 (9.7) |
| D5 | 245 (9.4) | 10,202 (10.0) | 51 (10.1) | 2,599 (9.6) |
| D6 | 244 (9.4) | 10,203 (10.0) | 48 (9.5) | 2,666 (9.9) |
| D7 | 251 (9.6) | 10,196 (10.0) | 42 (8.3) | 2,822 (10.5) |
| D8 | 257 (9.9) | 10,190 (10.0) | 45 (8.9) | 2,878 (10.7) |
| D9 | 246 (9.4) | 10,201 (10.0) | 54 (10.7) | 3,014 (11.2) |
| D10 (highest) | 297 (11.4) | 10,150 (10.0) | 65 (12.9) | 3,201 (11.9) |
| Bipolar disorder GPS decile[d] | | | | |
| N (%) missing | 104 (3.8) | 3,419 (3.3) | 0 (0.0) | 0 (0.0) |
| D1 (lowest), N (%)[c] | 199 (7.6) | 10,248 (10.1) | 39 (7.7) | 2,773 (10.3) |
| D2 | 225 (8.6) | 10,222 (10.0) | 45 (8.9) | 2,698 (10.0) |
| D3 | 232 (8.9) | 10,215 (10.0) | 47 (9.3) | 2,696 (10.0) |
| D4 | 228 (8.8) | 10,219 (10.0) | 47 (9.3) | 2,654 (9.8) |
| D5 | 228 (8.8) | 10,219 (10.0) | 36 (7.1) | 2,707 (10.0) |
| D6 | 264 (10.1) | 10,183 (10.0) | 46 (9.1) | 2,638 (9.8) |
| D7 | 274 (10.5) | 10,173 (10.0) | 49 (9.7) | 2,707 (10.0) |
| D8 | 262 (10.1) | 10,185 (10.0) | 48 (9.5) | 2,706 (10.0) |
| D9 | 306 (11.8) | 10,141 (10.0) | 63 (12.5) | 2,717 (10.1) |
| D10 (highest) | 387 (14.9) | 10,060 (9.9) | 84 (16.7) | 2,701 (10.0) |

BD, bipolar disorder; D, decile; GPS, genome-wide polygenic score; MET, metabolic equivalent of task; Q, quartile; Qu, quintile; SD, standard deviation.

a. Participants with complete data on all the covariates that were entered into the maximally-adjusted total effects models (age, gender, white British genetic ancestry, English-speaking country of birth, degree, comorbid neurological/psychiatric condition, family history of

dementia, family history of Parkinson's disease, family history of severe depression, maternal smoking around birth, childhood trauma, education/cognition GPS, bipolar disorder GPS).
b. No missing data.
c. Missing excluded from denominator.
d. Based on data distribution in the whole UK Biobank cohort.
e. Scottish psychiatric hospital records were unavailable, which meant no Scotland-based participants could be classified in the comparison group; therefore all locations for comparison participants are in England/Wales.
f. Apart from mood disorder or schizophrenia; not possible to distinguish between missing data and self-report of no condition, therefore both classified as 'No'.
g. From the web-based questionnaire, which was completed by 157,366 (31.3%) of the cohort.

#### Evaluation of the Graphical Model

The original DAG shown in eFigure 1 above postulated that educational attainment was a causal antecedent of mania/BD status, but a plausible alternative specification would show the arrow in reverse such that mania/BD causally influences educational attainment (for example, if illness onset occurs at a young age). The different predicted independencies implied by both specifications were tested. The results indicated poorer fit in the second specification, with a greater proportion of the partial correlation coefficients being above |0.1|. The first specification also showed poor fit involving certain nodes, particularly current psychotropic medication use. The reasons for poor fit (e.g. model misspecification, measurement error, selection bias) cannot be discerned from the data alone, but consideration was given to whether the graph should be modified by adding new nodes or paths.

It was deemed plausible that paths should be added between educational attainment and current psychotropic medication, and between gender and current psychotropic medication (reflecting possible influences of, for example, knowledge or attitudes on likelihood of seeking or accepting treatment). A new node was also added, representing other psychiatric or neurological conditions (apart from mood disorder or schizophrenia); this was in line with previous results,[58] which had indicated different patterns of cognitive impairment in participants with and without psychiatric or neurological comorbidities. The implied independencies of this modified DAG were then tested, firstly with educational attainment specified as a causal antecedent of both mania/BD and other psychiatric/neurological conditions, and then with educational attainment specified as a causal consequence of both.

The best fit was observed in the first version of the modified DAG, in which educational attainment was specified as a causal antecedent of both mania/BD and other psychiatric/neurological conditions. Twenty percent (28 of 138) of the partial correlation coefficients remained above |0.1|, but most of these were below |0.2| and the largest coefficient was |0.43|. The largest coefficients again involved the current psychotropic medication node. When evaluated further, however, regression models indicated wide variance in the estimated associations between the pairs of nodes, including the null in most instances (results not shown). No further modifications were made to the DAG; it was decided that the diagram broadly reflected the evidence-based assumptions drawn from the literature and expert knowledge, and additional data-driven modifications may have been misleading if they were in fact influenced by issues such as measurement error rather than model misspecification. The final DAG used in the analyses is shown in eFigure 3 below.

#### Total Effects

The best propensity score model, in terms of covariate balance, was the first model with no interaction terms. This was used in all the outcome models that involved propensity score adjustment or matching, and was used as the basis for the inverse probability weights. eTable 6 shows the degree of covariate balance, as illustrated within the matched samples used in the reaction time analyses.

eFigures 4 to 7 show the total effects results for prospective memory, reasoning, reaction time, and numeric memory.

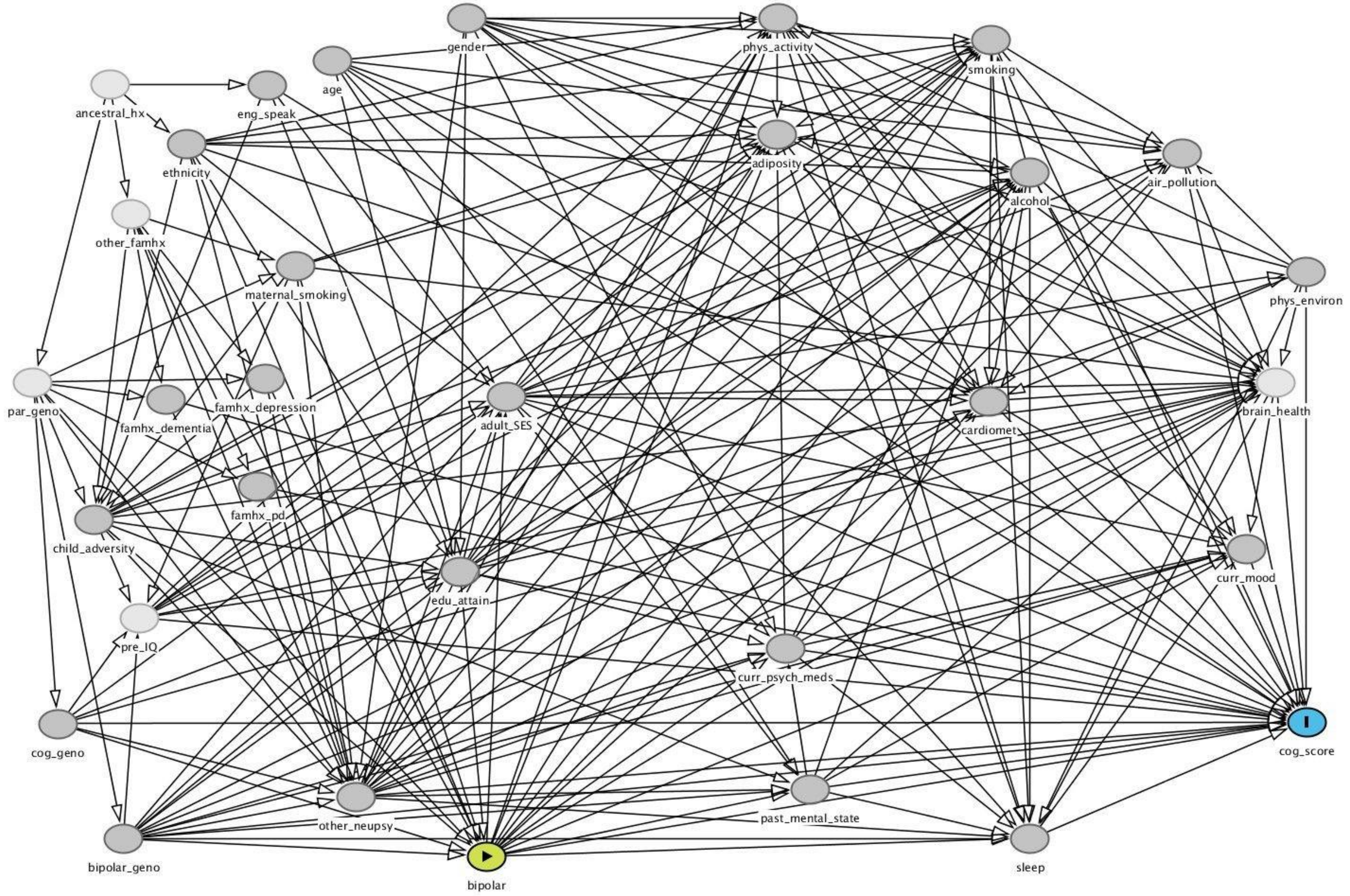

eFigure 3 Final DAG used in the mania/BD analyses

Cardiomet, cardiometabolic disease; cog, cognitive; curr, current; edu, educational; eng_speak, English speaking birth country; famhx, family history; geno, genotype; hx, history; other_neupsy, other comorbid neurological/psychiatric condition; par, parental; PD, Parkinson's disease; phys, physical; pre_IQ, premorbid intelligence; psych_meds, psychotropic medications; SES, socioeconomic status. Green node is the exposure and blue node is the outcome. Light nodes represent unmeasured constructs and darker nodes represent measured constructs. This graph and the underlying code are publicly accessible online at dagitty.net/mSTG_SM *eTable 6 Summary of covariates in matched mania/bipolar and comparison groups*

| | Mania/BD | Comparison |
|---|---|---|
| | Mean (SD) | |
| Age (years) | 54.9 (7.4) | 55.1 (7.4) |
| | % | |
| Female gender | 52.9 | 52.0 |
| English-speaking country of birth | 97.7 | 98.3 |
| Has a degree | 52.4 | 55.2 |
| Comorbid neurological or psychiatric condition | 25.2 | 25.1 |
| Family history of dementia | 18.6 | 17.5 |
| Family history of Parkinson's disease | 3.6 | 4.3 |
| Family history of severe depression | 34.6 | 36.2 |
| Maternal smoking around birth | 33.1 | 34.5 |
| Any childhood trauma | 60.8 | 60.2 |
| Education/cognition GPS | | |
| D1 (lowest) | 7.5 | 7.1 |
| D2 | 9.9 | 8.6 |
| D3 | 10.2 | 8.0 |
| D4 | 10.7 | 12.3 |
| D5 | 10.4 | 9.9 |
| D6 | 9.2 | 10.3 |
| D7 | 8.7 | 7.8 |
| D8 | 8.7 | 10.0 |
| D9 | 11.5 | 11.1 |
| D10 (highest) | 13.2 | 14.9 |
| Bipolar disorder GPS | | |
| D1 (lowest) | 7.8 | 6.6 |
| D2 | 9.7 | 10.3 |
| D3 | 7.6 | 8.5 |
| D4 | 9.7 | 9.3 |
| D5 | 7.1 | 7.8 |
| D6 | 8.9 | 9.9 |
| D7 | 10.2 | 10.2 |
| D8 | 9.4 | 9.5 |
| D9 | 11.5 | 11.4 |
| D10 (highest) | 18.1 | 16.5 |

BD, bipolar disorder; D, decile; GPS, genome-wide polygenic score; SD, standard deviation.
These results are from the propensity-score matched samples used in the 1:3 matched model for the total effect of mania/BD on reaction time (mania/BD N = 392; comparison N = 1,081).

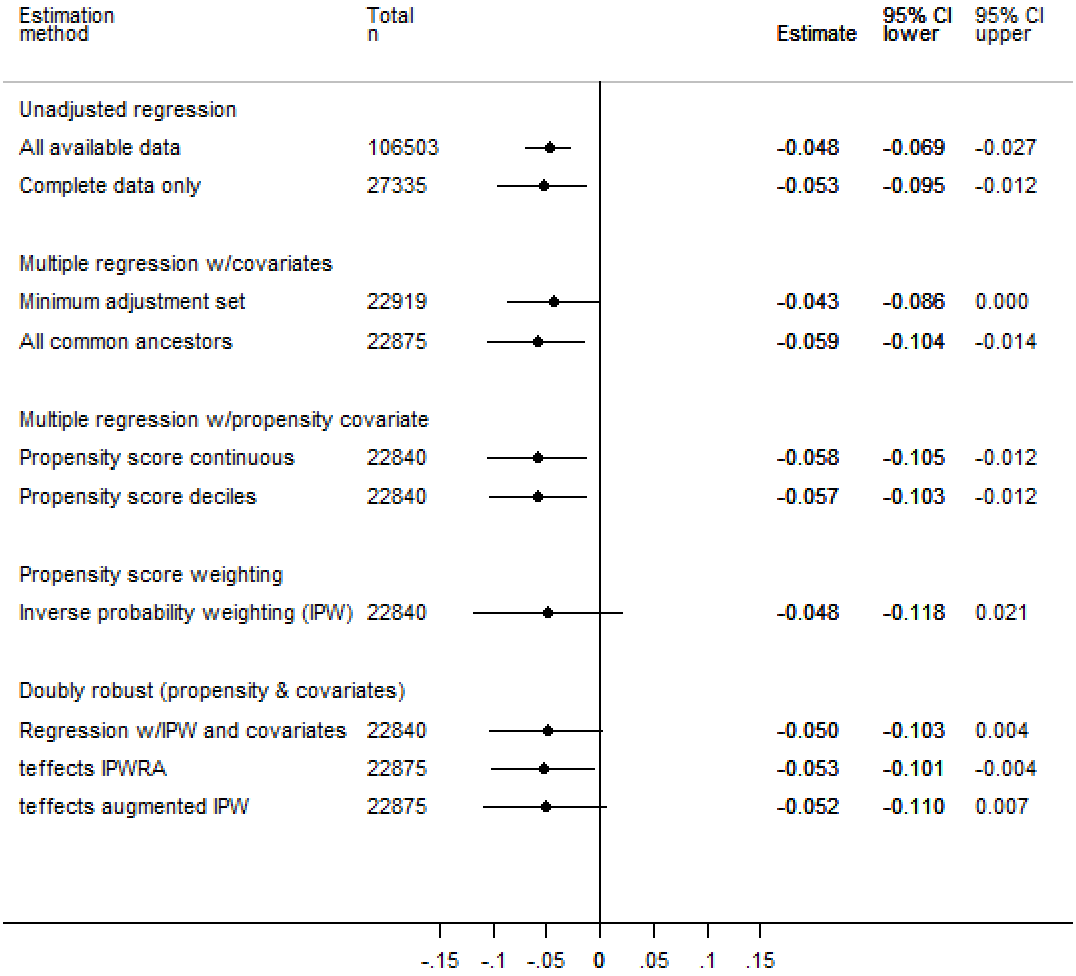

*eFigure 4 Total effect of mania/bipolar disorder on prospective memory*

CI, confidence interval; IPW, inverse probability weighting; IPWRA, inverse probability weighting with regression adjustment; teffects, Stata teffects package. Estimates are proportions and can be interpreted as risk differences. No estimates are provided from propensity-score matched models as it was not possible to express these as risk differences.

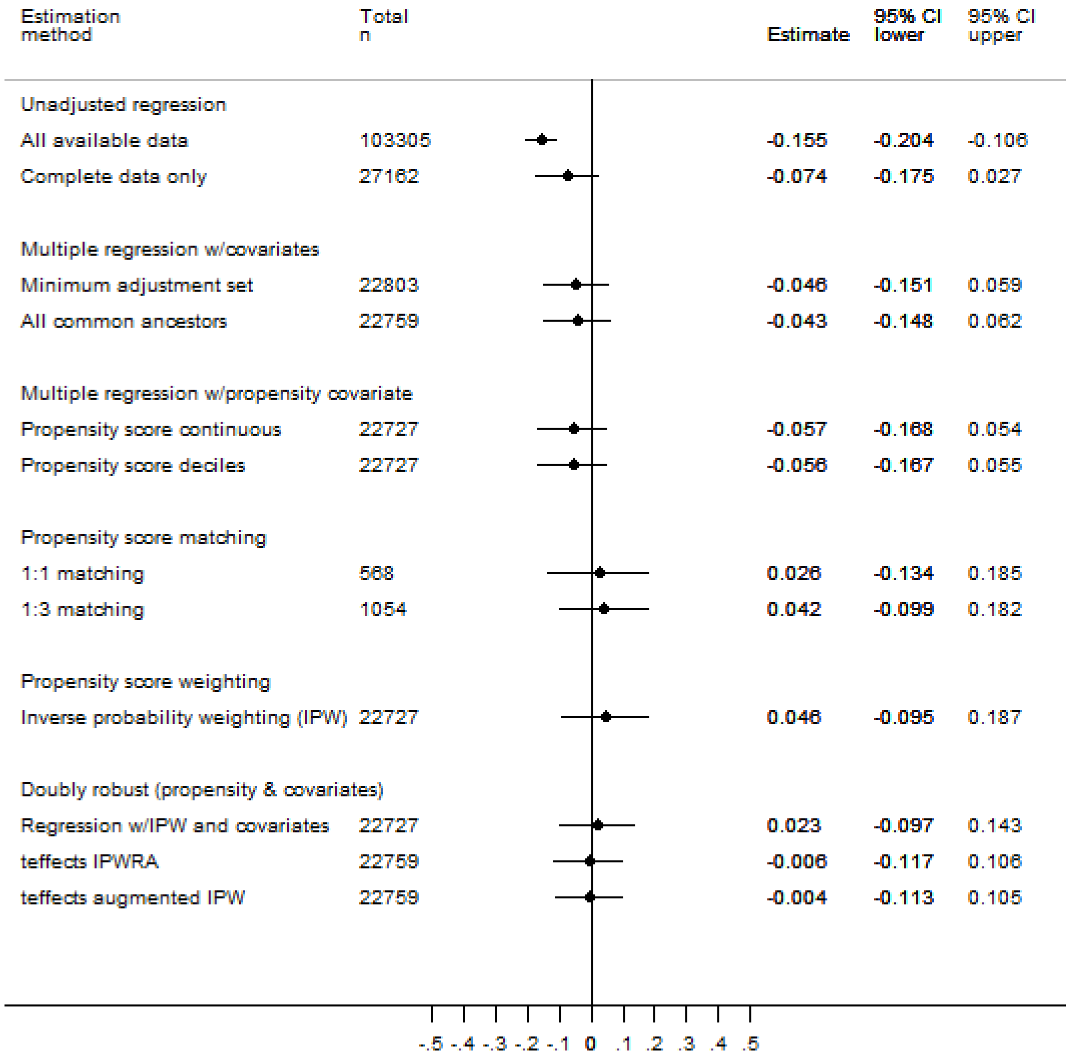

*eFigure 5* *Total effect of mania/bipolar disorder on reasoning*

CI, confidence interval; IPW, inverse probability weighting; IPWRA, inverse probability weighting with regression adjustment; teffects, Stata teffects package. Estimates are in z-score units and can be interpreted as standardized mean differences.

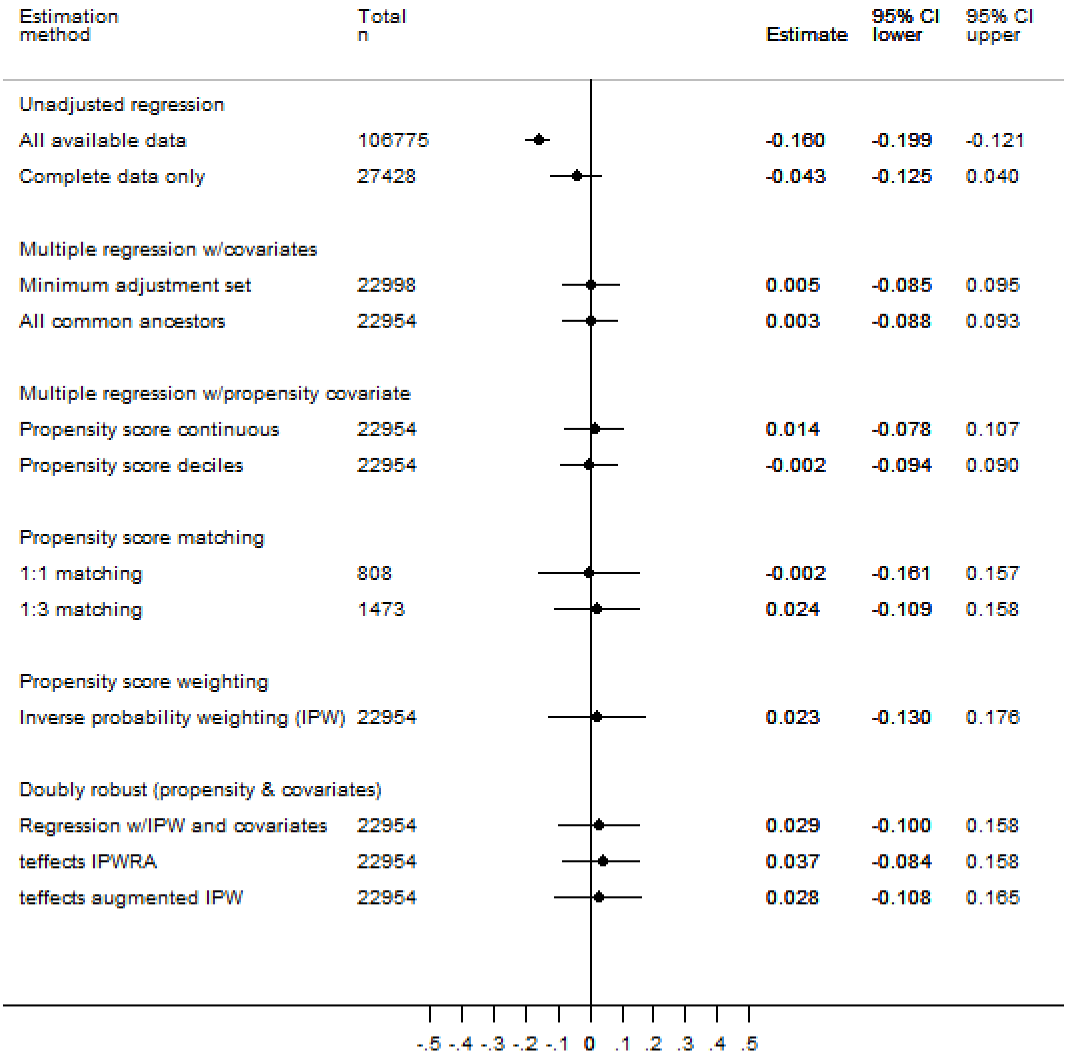

*eFigure 6 Total effect of mania/bipolar disorder on reaction time*

CI, confidence interval; IPW, inverse probability weighting; IPWRA, inverse probability weighting with regression adjustment; teffects, Stata teffects package. Estimates are in z-score units and can be interpreted as standardized mean differences.

| Estimation method | Total n | Estimate | 95% CI lower | 95% CI upper |
|---|---|---|---|---|
| Unadjusted regression | | | | |
| All available data | 30867 | -0.170 | -0.264 | -0.077 |
| Complete data only | 7853 | -0.063 | -0.280 | 0.153 |
| Multiple regression w/covariates | | | | |
| Minimum adjustment set | 7001 | 0.002 | -0.224 | 0.228 |
| All common ancestors | 6983 | 0.003 | -0.225 | 0.230 |
| Multiple regression w/propensity covariate | | | | |
| Propensity score continuous | 6972 | 0.005 | -0.228 | 0.238 |
| Propensity score deciles | 6972 | 0.014 | -0.219 | 0.248 |
| Propensity score matching | | | | |
| 1:1 matching | 158 | 0.179 | -0.126 | 0.484 |
| 1:3 matching | 308 | 0.123 | -0.145 | 0.392 |
| Propensity score weighting | | | | |
| Inverse probability weighting (IPW) | 6972 | 0.089 | -0.186 | 0.364 |
| Doubly robust (propensity & covariates) | | | | |
| Regression w/IPW and covariates | 6972 | 0.048 | -0.176 | 0.272 |
| teffects IPWRA | 6983 | -0.021 | -0.205 | 0.162 |
| teffects augmented IPW | 6983 | 0.001 | -0.256 | 0.259 |

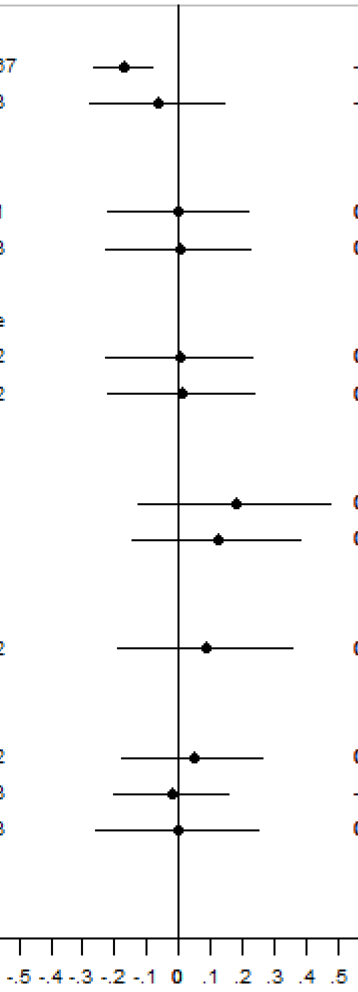

*eFigure 7 Total effect of mania/bipolar disorder on numeric memory*

CI, confidence interval; IPW, inverse probability weighting; IPWRA, inverse probability weighting with regression adjustment; teffects, Stata teffects package. Estimates are in z-score units and can be interpreted as standardized mean differences.

#### Mediation Analyses

Mediation analyses were conducted to quantify the proportion of the total effect of mania/BD on each cognitive outcome that was accounted for by (1) cardiometabolic disease and (2) psychotropic medication. Effects via other intermediate nodes of interest (e.g. current depressive symptoms) could not be identified, because no covariate adjustment set could be found for the relevant exposure-mediator and/or mediator-outcome paths.

In both sets of analyses, the indirect pathways were affected by intermediate confounding (i.e. the mediator-outcome path was confounded by at least one node that descended from mania/BD), and so the analyses were conducted under the identifying assumption of no interaction between mania/BD and either cardiometabolic disease or psychotropic medication.[59] This assumption was checked by conducting a regression model of each cognitive outcome on mania/BD exposure status, the mediator and all the covariates, including a product term for mania/BD * mediator; there was no evidence of interaction in any of the models (see eTable 7 below). The mediation model estimates are interpreted as 'randomized interventional analogs' of the natural direct and indirect effects.[60]

*eTable 7 Tests of interactions between exposure and mediators in the mania/BD analyses*

| | **Coefficient for mania/BD * mediator** | **95% CI** | ***P*** |
|---|---|---|---|
| **Mediator: Cardiometabolic disease** | | | |
| Reasoning[a] | 0.029 | -0.207, 0.264 | 0.812 |
| Reaction time[a] | 0.149 | -0.047, 0.345 | 0.137 |
| Numeric memory[a] | -0.124 | -0.652, 0.403 | 0.644 |
| Visuospatial memory[a] | 0.239 | -0.019, 0.497 | 0.070 |
| Prospective memory[b] | 0.991 | 0.475, 2.071 | 0.982 |
| **Mediator: Psychotropic medication** | | | |
| Reasoning[a] | 0.005 | -0.255, 0.264 | 0.973 |
| Reaction time[a] | 0.095 | -0.110, 0.301 | 0.364 |
| Numeric memory[a] | 0.311 | -0.294, 0.916 | 0.313 |
| Visuospatial memory[a] | -0.045 | -0.295, 0.205 | 0.725 |
| Prospective memory[b] | 0.883 | 0.435, 1.796 | 0.732 |

BD, bipolar disorder; CI, confidence interval.
All models included mania/BD, the mediator and their product, as well as all the covariates entered into the gformula mediation models.
a. Linear regression model with outcome measured in z-score units.
b. Logistic regression model with outcome measured as correct or not; estimate expressed as odds ratio.

*eTable 8 Mediation of the effect of mania/bipolar disorder on cognitive outcome via cardiometabolic disease*

| | N | Estimate | 95% CI[a] |
|---|---|---|---|
| **Reasoning**[b] | 21,043 | | |
| TE | | -0.049 | -0.157, 0.059 |
| CDE | | -0.048 | -0.156, 0.061 |
| NDE | | -0.045 | -0.153, 0.063 |
| NIE | | -0.004 | -0.020, 0.013 |
| **Reaction time**[b] | 21,213 | | |
| TE | | 0.014 | -0.082, 0.111 |
| CDE | | 0.034 | -0.063, 0.132 |
| NDE | | 0.007 | -0.090, 0.103 |
| NIE | | 0.008 | -0.011, 0.026 |
| **Numeric memory**[b] | 6,396 | | |
| TE | | 0.047 | -0.213, 0.306 |
| CDE | | 0.021 | -0.237, 0.280 |
| NDE | | 0.023 | -0.235, 0.281 |
| NIE | | 0.024 | -0.008, 0.055 |
| **Visuospatial memory**[b] | 21,124 | | |
| TE | | -0.166 | -0.285, -0.047 |
| CDE | | -0.160 | -0.277, -0.044 |
| NDE | | -0.164 | -0.282, -0.046 |
| NIE | | -0.002 | -0.022, 0.017 |
| **Prospective memory**[c] | 21,139 | | |
| TE | | -0.033 | -0.078, 0.012 |
| CDE | | -0.038 | -0.083, 0.006 |
| NDE | | -0.038 | -0.083, 0.007 |
| NIE | | 0.005 | -0.002, 0.012 |

CDE, controlled direct effect when cardiometabolic disease = 0; CI, confidence interval; GPS, genome-wide polygenic score; NDE, natural direct effect; NIE, natural indirect effect; $NO_2$, nitrogen dioxide; $PM_{10}$, particulate matter of up to 10μm diameter; TE, total effect. Models were restricted to participants of white British genetic ancestry, and were adjusted for age, gender, educational attainment, English-speaking birth country, education/cognition GPS, bipolar disorder GPS, family history of dementia, family history of Parkinson's disease, maternal smoking around birth, childhood trauma, other psychiatric/neurological conditions, deprivation, population density, road proximity, air pollution ($PM_{10}$ and $NO_2$), body mass index, alcohol frequency, smoking status, physical activity, and psychotropic medication.

a. Normal-based, from bootstrapped standard error (1000 replicates).

b. Estimate expressed as a standardized mean difference.

c. Estimate expressed as a risk difference for the probability of being correct.

*eTable 9 Mediation of the effect of mania/bipolar disorder on cognitive outcome via psychotropic medication*

| | N | Estimate | 95% CI[a] |
|---|---|---|---|
| **Reasoning**[b] | 21,339 | | |
| TE | | -0.065 | -0.176, 0.047 |
| CDE | | 0.016 | -0.098, 0.130 |
| NDE | | -0.015 | -0.130, 0.099 |
| NIE | | -0.049 | -0.077, -0.021 |
| **Reaction time**[b] | 21,518 | | |
| TE | | -0.005 | -0.103, 0.094 |
| CDE | | 0.073 | -0.036, 0.182 |
| NDE | | 0.036 | -0.072, 0.144 |
| NIE | | -0.040 | -0.075, -0.005 |
| **Numeric memory**[b] | 6,547 | | |
| TE | | 0.033 | -0.203, 0.270 |
| CDE | | 0.058 | -0.181, 0.296 |
| NDE | | 0.075 | -0.163, 0.312 |
| NIE | | -0.041 | -0.097, 0.015 |
| **Visuospatial memory**[b] | 21,424 | | |
| TE | | -0.194 | -0.311, -0.077 |
| CDE | | -0.098 | -0.220, 0.024 |
| NDE | | -0.140 | -0.263, -0.018 |
| NIE | | -0.054 | -0.094, -0.013 |
| **Prospective memory**[c] | 21,436 | | |
| TE | | -0.037 | -0.082, 0.009 |
| CDE | | -0.028 | -0.072, 0.016 |
| NDE | | -0.018 | -0.062, 0.026 |
| NIE | | -0.019 | -0.030, -0.007 |

CDE, controlled direct effect when psychotropic medication = 0; CI, confidence interval; GPS, genome-wide polygenic score; NDE, natural direct effect; NIE, natural indirect effect; TE, total effect.
Models were restricted to participants of white British genetic ancestry, and were adjusted for gender, educational attainment, English-speaking birth country, education/cognition GPS, bipolar disorder GPS, family history of dementia, family history of Parkinson's disease, maternal smoking around birth, childhood trauma, other psychiatric/neurological conditions, deprivation, and lifetime number of episodes of depressed mood/anhedonia.
a. Normal-based, from bootstrapped standard error (1000 replicates).
b. Estimate expressed as a standardized mean difference.
c. Estimate expressed as a risk difference for the probability of being correct.

#### Sensitivity Analyses

Rosenbaum bounds were calculated to check the sensitivity of the visuospatial memory total effects result to departures from exchangeability. The estimated effect crossed the null at a gamma value of 1.2, which corresponds to the probability of being in the exposed group being approximately 0.45 or 0.55, rather than 0.5 as would be the case if the groups were truly exchangeable. This indicates that the results would not be robust to an unmeasured confounder with even a weak association with group membership.

There was evidence of missing data bias on several of the outcome measures: much of the attenuation towards the null seen in the total effects models occurred prior to any multivariable adjustment/matching, simply by restricting the sample to participants with complete covariate data. When the multiple linear regression models for total effects were repeated after multiple imputation of the covariate values, the results showed less attenuation (eFigure 8 below). The effect size estimates in these reasoning, reaction time and numeric memory models were of small magnitude (point estimates approximately -0.10 to -0.15). The estimates in the visuospatial memory models remained similar regardless of which estimation method was used (approximate point estimate -0.19).

When the mediation models were repeated with imputation of missing mediator and covariate values, the proportion of the total effect transmitted indirectly via cardiometabolic disease remained negligible (eTable 10), whereas there was evidence of indirect effects via psychotropic medication for all the cognitive outcomes (eTable 11).

The results of the probabilistic analysis using `episens` indicated that the total effects estimate for visuospatial memory is likely to be sensitive to exposure misclassification. When dichotomized into impaired and unimpaired outcome categories, and assuming no exposure misclassification, the unadjusted relative risk of impairment was 1.70 in the mania/BD group (95% CI 1.45, 2.01). Assuming lower sensitivity to true mania/BD status among the cognitively impaired (sensitivity range 0.6 to 0.9) versus unimpaired participants (sensitivity range 0.7 to 1.0), the relative risk was estimated as 1.81 (0.22, 11.44).

The DAGitty algorithm determined that there were six other DAGs that were equivalent to the final DAG used in the present analyses. None of these alternative configurations was causally plausible, owing to temporal order constraints (e.g. it is not possible for parental genotype to be causally influenced by offspring genotype). DAGitty also generated minimum sufficient covariate adjustment sets for the equivalent DAGs, for the total effect of mania/BD on cognitive outcome. The same minimum adjustment set was valid for the analysed DAG and for the six equivalent DAGs. This indicates that the multivariable analyses reported here remain valid, regardless of which model within the equivalence class is correct.

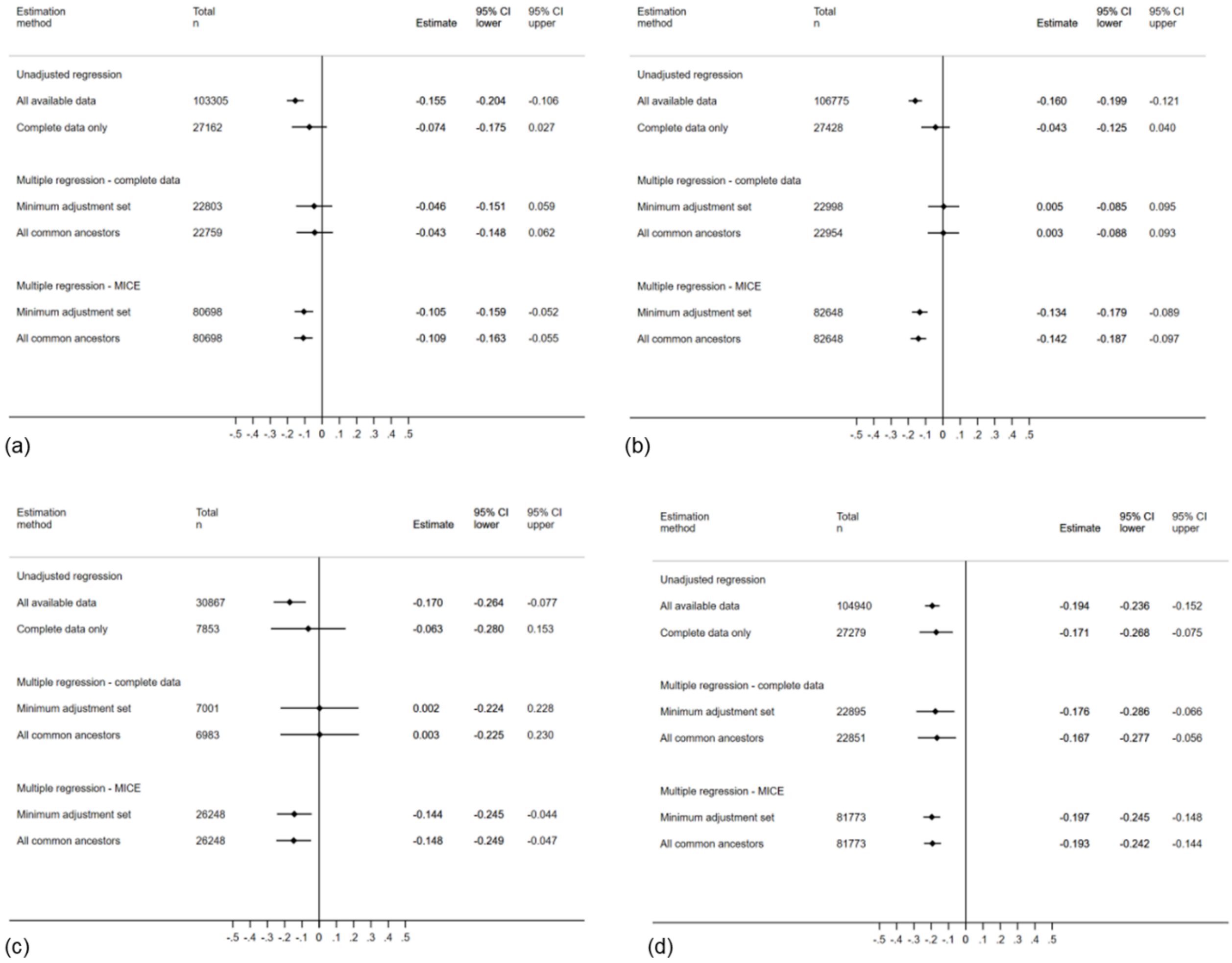

*eFigure 8 Comparison of missing data approaches in mania/bipolar disorder total effects analyses*

CI, confidence interval; MICE, multiple imputation with chained equations. Panels show: (a) reasoning, (b) reaction time, (c) numeric memory and (d) visuospatial memory. Prospective memory not shown as it was not possible to calculate risk differences.

*eTable 10 Mediation of the effect of mania/bipolar disorder on cognitive outcome via cardiometabolic disease, with missing data imputation*

| | N | Estimate | 95% CI[a] |
|---|---|---|---|
| **Reasoning**[b] | 80,698 | | |
| TE | | -0.092 | -0.158, -0.026 |
| CDE | | -0.087 | -0.152, -0.022 |
| NDE | | -0.085 | -0.151, -0.019 |
| NIE | | -0.007 | -0.017, 0.003 |
| **Reaction time**[b] | 82,648 | | |
| TE | | -0.094 | -0.150, -0.037 |
| CDE | | -0.091 | -0.148, -0.034 |
| NDE | | -0.091 | -0.148, -0.033 |
| NIE | | -0.003 | -0.014, 0.008 |
| **Numeric memory**[b] | 26,248 | | |
| TE | | -0.142 | -0.275, -0.008 |
| CDE | | -0.158 | -0.291, -0.024 |
| NDE | | -0.150 | -0.283, -0.018 |
| NIE | | 0.008 | -0.011, 0.027 |
| **Visuospatial memory**[b] | 81,773 | | |
| TE | | -0.206 | -0.267, -0.145 |
| CDE | | -0.200 | -0.261, -0.139 |
| NDE | | -0.210 | -0.272, -0.149 |
| NIE | | 0.005 | -0.007, 0.016 |
| **Prospective memory**[c] | 82,194 | | |
| TE | | -0.041 | -0.062, -0.021 |
| CDE | | -0.037 | -0.057, -0.017 |
| NDE | | -0.039 | -0.059, -0.018 |
| NIE | | -0.003 | -0.006, 0.001 |

CDE, controlled direct effect when cardiometabolic disease = 0; CI, confidence interval; GPS, genome-wide polygenic score; NDE, natural direct effect; NIE, natural indirect effect; $NO_2$, nitrogen dioxide; $PM_{10}$, particulate matter of up to 10μm diameter; TE, total effect. Models were restricted to participants of white British genetic ancestry, and were adjusted for age, gender, educational attainment, English-speaking birth country, education/cognition GPS, bipolar disorder GPS, family history of dementia, family history of Parkinson's disease, maternal smoking around birth, childhood trauma, other psychiatric/neurological conditions, deprivation, population density, road proximity, air pollution ($PM_{10}$ and $NO_2$), body mass index, alcohol frequency, smoking status, physical activity, and psychotropic medication. Missing mediator and covariate data were imputed via a single stochastic imputation using chained equations.

a. Normal-based, from bootstrapped standard error (1000 replicates).

b. Estimate expressed as a standardized mean difference.

c. Estimate expressed as a risk difference for the probability of being correct.

*eTable 11 Mediation of the effect of mania/bipolar disorder on cognitive outcome via psychotropic medication, with missing data imputation*

| | N | Estimate | 95% CI[a] |
|---|---|---|---|
| **Reasoning**[b] | 80,698 | | |
| TE | | -0.095 | -0.162, -0.027 |
| CDE | | -0.034 | -0.103, 0.035 |
| NDE | | -0.034 | -0.103, 0.035 |
| NIE | | -0.061 | -0.078, -0.044 |
| **Reaction time**[b] | 82,648 | | |
| TE | | -0.080 | -0.136, -0.024 |
| CDE | | -0.034 | -0.095, 0.028 |
| NDE | | -0.024 | -0.085, 0.037 |
| NIE | | -0.056 | -0.078, -0.033 |
| **Numeric memory**[b] | 26,248 | | |
| TE | | -0.147 | -0.276, -0.018 |
| CDE | | -0.079 | -0.210, 0.051 |
| NDE | | -0.082 | -0.212, 0.049 |
| NIE | | -0.065 | -0.098, -0.033 |
| **Visuospatial memory**[b] | 81,773 | | |
| TE | | -0.196 | -0.255, -0.137 |
| CDE | | -0.147 | -0.210, -0.084 |
| NDE | | -0.146 | -0.209, -0.083 |
| NIE | | -0.050 | -0.076, -0.024 |
| **Prospective memory**[c] | 82,194 | | |
| TE | | -0.025 | -0.045, -0.005 |
| CDE | | -0.010 | -0.029, 0.010 |
| NDE | | -0.012 | -0.031, 0.007 |
| NIE | | -0.014 | -0.019, -0.008 |

CDE, controlled direct effect when psychotropic medication = 0; CI, confidence interval; GPS, genome-wide polygenic score; NDE, natural direct effect; NIE, natural indirect effect; TE, total effect.
Models were restricted to participants of white British genetic ancestry, and were adjusted for gender, educational attainment, English-speaking birth country, education/cognition GPS, bipolar disorder GPS, family history of dementia, family history of Parkinson's disease, maternal smoking around birth, childhood trauma, other psychiatric/neurological conditions, deprivation, and lifetime number of episodes of depressed mood/anhedonia. Missing mediator and covariate data were imputed via a single stochastic imputation using chained equations.
a. Normal-based, from bootstrapped standard error (1000 replicates).
b. Estimate expressed as a standardized mean difference.
c. Estimate expressed as a risk difference for the probability of being correct.

### Cognitive Impairment in Major Depression

#### Characteristics of the Sample

eFigure 9 shows a flowchart of exclusions leading to the major depression analysis sample. As with the mania/BD analyses, a large number of participants were excluded due to missing data in at least one exposure information source, which meant they could not be classified in the comparison group.

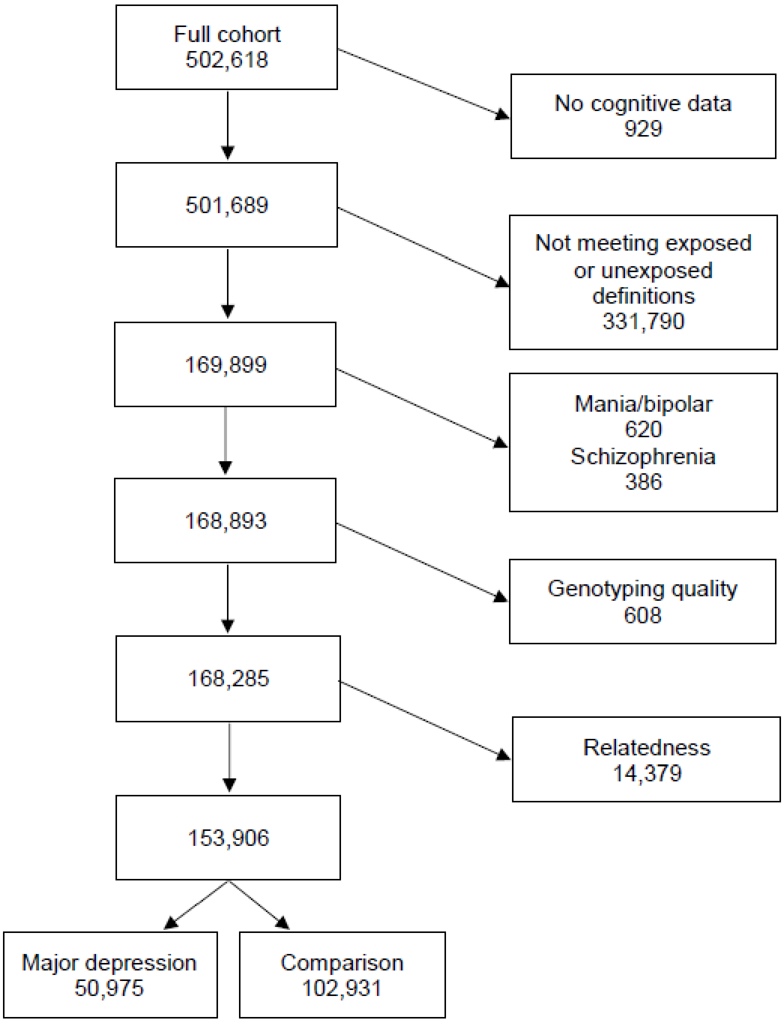

*eFigure 9 Major depression analysis sample flowchart*

eTable 12 describes the covariate data in both groups. As with the mania/BD analyses, the proportion of missing data was highest, by far, on the childhood trauma variable, and was also relatively high on the family history, current depressive symptoms and physical activity measures. The descriptive information indicated that the major depression group was younger, on average, than the comparison group, had a substantially higher proportion of women, and had higher proportions of current smokers, former drinkers, and participants living in deprived areas. The major depression group had higher proportions with frequent sleeplessness, obesity, cardiometabolic disease, comorbid neurological/psychiatric conditions, family history of severe depression, current psychotropic medication and history of childhood trauma, and they reported more depressed episodes and higher current depressive symptoms on average. The distribution of the major depression GPS score was skewed towards higher values in the major depression group. The subset of participants with complete covariate data appeared different from the full analysis sample across multiple measures, e.g. having a greater proportion of degree-holders and a smaller proportion from the most deprived areas.

*eTable 12 Summary of covariates in the major depression and comparison groups*

| | All available data | | Complete covariate data[a] | |
|---|---|---|---|---|
| | **Major depression** | **Comparison** | **Major depression** | **Comparison** |
| N | 50,975 | 102,931 | 50,975 | 102,931 |
| **Sociodemographic** | | | | |
| Age (years)[b] | | | | |
| Mean (SD) | 55.6 (7.9) | 57.0 (8.2) | 55.6 (7.9) | 57.0 (8.2) |
| Gender[b] | | | | |
| N (%) female | 33,090 (64.9) | 51,463 (50.0) | 33,090 (64.9) | 51,463 (50.0) |
| Ethnic group | | | | |
| N (%) missing | 220 (0.4) | 401 (0.4) | 220 (0.4) | 401 (0.4) |
| White, N (%)[c] | 48,345 (95.3) | 93,194 (90.9) | 48,345 (95.3) | 93,194 (90.9) |
| Asian/Asian British | 814 (1.6 ) | 3,663 (3.6) | 814 (1.6 ) | 3,663 (3.6) |
| Black/Black British | 641 (1.3) | 3,159 (3.1) | 641 (1.3) | 3,159 (3.1) |
| Chinese | 69 (0.1) | 473 (0.5) | 69 (0.1) | 473 (0.5) |
| Mixed & other background | 886 (1.8) | 2,041 (2.0) | 886 (1.8) | 2,041 (2.0) |
| White British genetic ancestry | | | | |
| N (%) missing | 1,518 (3.0) | 3,419 (3.3) | 1,518 (3.0) | 3,419 (3.3) |
| N (%)[c] | 41,509 (83.9) | 79,097 (79.5) | 41,509 (83.9) | 79,097 (79.5) |
| English-speaking country of birth | | | | |
| N (%) missing | 85 (0.2) | 149 (0.2) | 85 (0.2) | 149 (0.2) |
| N (%)[c] | 47,741 (93.8) | 91,524 (89.1) | 47,741 (93.8) | 91,524 (89.1) |
| Has a degree | | | | |
| N (%) missing | 493 (1.0) | 1,012 (1.0) | 493 (1.0) | 1,012 (1.0) |
| N (%)[c] | 16,713 (33.1) | 36,211 (35.5) | 16,713 (33.1) | 36,211 (35.5) |
| Townsend quintile[d] | | | | |
| N (%) missing | 93 (0.2) | 159 (0.2) | 93 (0.2) | 159 (0.2) |
| Qu1 (least deprived), N (%)[c] | 8,109 (15.9) | 17,672 (17.2) | 8,109 (15.9) | 17,672 (17.2) |
| Qu2 | 9,124 (17.9) | 20,807 (20.3) | 9,124 (17.9) | 20,807 (20.3) |
| Qu3 | 9,793 (19.3) | 21,291 (20.7) | 9,793 (19.3) | 21,291 (20.7) |
| Qu4 | 11,261 (22.1) | 23,193 (22.6) | 11,261 (22.1) | 23,193 (22.6) |
| Qu5 (most deprived) | 12,595 (24.8) | 19,809 (19.3) | 12,595 (24.8) | 19,809 (19.3) |
| **Local environment** | | | | |
| Home area population density[e] | | | | |
| N (%) missing | 596 (1.2) | 951 (0.9) | 596 (1.2) | 951 (0.9) |
| England/Wales urban, N (%)[c] | 42,462 (84.3) | 88,651 (86.9) | 42,462 (84.3) | 88,651 (86.9) |
| England/Wales town | 3,239 (6.4) | 6,392 (6.3) | 3,239 (6.4) | 6,392 (6.3) |
| England/Wales village | 2,031 (4.0) | 4,853 (4.8) | 2,031 (4.0) | 4,853 (4.8) |
| England/Wales hamlet/isolated | 842 (1.7) | 2,084 (2.0) | 842 (1.7) | 2,084 (2.0) |
| Scotland large urban | 1,376 (2.7) | 0 (0.0) | 1,376 (2.7) | 0 (0.0) |
| Scotland other urban | 300 (0.6) | 0 (0.0) | 300 (0.6) | 0 (0.0) |
| Scotland small town | 66 (0.1) | 0 (0.0) | 66 (0.1) | 0 (0.0) |
| Scotland rural | 63 (0.1) | 0 (0.0) | 63 (0.1) | 0 (0.0) |
| Proximity to major road (1/m) | | | | |
| N (%) missing | 772 (1.5) | 1,290 (1.3) | 772 (1.5) | 1,290 (1.3) |
| Mean (SD) | 0.006 (0.014) | 0.006 (0.013) | 0.006 (0.014) | 0.006 (0.013) |
| Particulate matter ≤10µm (µg/m$^3$) | | | | |
| N (%) missing | 938 (1.8) | 1,668 (1.6) | 938 (1.8) | 1,668 (1.6) |
| Mean (SD) | 22.4 (2.9) | 22.8 (3.1) | 22.4 (2.9) | 22.8 (3.1) |

| | All available data | | Complete covariate data[a] | |
|---|---|---|---|---|
| | **Major depression** | **Comparison** | **Major depression** | **Comparison** |
| Nitrogen dioxide (μg/m$^3$) | | | | |
| N (%) missing | 772 (1.5) | 1,290 (1.3) | 772 (1.5) | 1,290 (1.3) |
| Mean (SD) | 30.9 (10.1) | 31.9 (10.6) | 30.9 (10.1) | 31.9 (10.6) |
| **Lifestyle and physical** | | | | |
| Smoking status | | | | |
| N (%) missing | 170 (0.3) | 368 (0.4) | 170 (0.3) | 368 (0.4) |
| Never, N (%)[c] | 25,008 (49.2) | 58,955 (57.5) | 25,008 (49.2) | 58,955 (57.5) |
| Former | 18,320 (36.1) | 34,658 (33.8) | 18,320 (36.1) | 34,658 (33.8) |
| Current | 7,477 (14.7) | 8,950 (8.7) | 7,477 (14.7) | 8,950 (8.7) |
| Alcohol frequency | | | | |
| N (%) missing | 100 (0.2) | 80 (0.1) | 100 (0.2) | 80 (0.1) |
| Daily/almost daily, N (%)[c] | 9,839 (19.3) | 21,729 (21.1) | 9,839 (19.3) | 21,729 (21.1) |
| 3-4 times per week | 10,085 (19.8) | 24,196 (23.5) | 10,085 (19.8) | 24,196 (23.5) |
| 1-2 times per week | 11,765 (23.1) | 26,111 (25.4) | 11,765 (23.1) | 26,111 (25.4) |
| 1-3 times per month | 6,344 (12.5) | 11,118 (10.8) | 6,344 (12.5) | 11,118 (10.8) |
| Special occasions only | 7,333 (14.4) | 11,536 (11.2) | 7,333 (14.4) | 11,536 (11.2) |
| Never (former drinker) | 3,189 (6.3) | 3,096 (3.0) | 3,189 (6.3) | 3,096 (3.0) |
| Never (not former drinker) | 2,320 (4.6) | 5,065 (4.9) | 2,320 (4.6) | 5,065 (4.9) |
| Sleeplessness | | | | |
| N (%) missing | 58 (0.1) | 79 (0.1) | 58 (0.1) | 79 (0.1) |
| Never/rarely, N (%)[c] | 8,438 (16.6) | 28,617 (27.8) | 8,438 (16.6) | 28,617 (27.8) |
| Sometimes | 23,090 (45.4) | 49,193 (47.8) | 23,090 (45.4) | 49,193 (47.8) |
| Usually | 19,389 (38.1) | 25,042 (24.4) | 19,389 (38.1) | 25,042 (24.4) |
| Physical activity (MET h/week) | | | | |
| N (%) missing | 4,617 (9.1) | 6,690 (6.5) | 4,617 (9.1) | 6,690 (6.5) |
| Median (Q1, Q3) | 26.2 (11.6, 55.9) | 29.9 (13.8, 60.2) | 26.2 (11.6, 55.9) | 29.9 (13.8, 60.2) |
| Body mass index | | | | |
| N (%) missing | 354 (0.7) | 732 (0.7) | 354 (0.7) | 732 (0.7) |
| Underweight, N (%)[c] | 270 (0.5) | 496 (0.5) | 270 (0.5) | 496 (0.5) |
| Normal | 15,166 (30.0) | 34,192 (33.5) | 15,166 (30.0) | 34,192 (33.5) |
| Overweight | 20,289 (40.1) | 43,942 (43.0) | 20,289 (40.1) | 43,942 (43.0) |
| Obese class I | 9,781 (19.3) | 17,374 (17.0) | 9,781 (19.3) | 17,374 (17.0) |
| Obese class II | 3,443 (6.8) | 4,588 (4.5) | 3,443 (6.8) | 4,588 (4.5) |
| Obese class III | 1,672 (3.3) | 1,607 (1.6) | 1,672 (3.3) | 1,607 (1.6) |
| **Medical and family history** | | | | |
| Cardiometabolic disease | | | | |
| N (%) missing | 121 (0.2) | 177 (0.2) | 121 (0.2) | 177 (0.2) |
| N (%)[c] | 17,459 (34.3) | 31,361 (30.5) | 17,459 (34.3) | 31,361 (30.5) |
| Comorbid neurological or psychiatric condition[f] | | | | |
| N (%) | 11,542 (22.6) | 9,155 (8.9) | 11,542 (22.6) | 9,155 (8.9) |
| Family history of dementia | | | | |
| N (%) missing | 7,815 (15.3) | 14,476 (14.1) | 7,815 (15.3) | 14,476 (14.1) |
| N (%)[c] | 7,032 (16.3) | 15,330 (17.3) | 7,032 (16.3) | 15,330 (17.3) |
| Family history of Parkinson's disease | | | | |
| N (%) missing | 9,456 (18.6) | 16,090 (15.6) | 9,456 (18.6) | 16,090 (15.6) |
| N (%)[c] | 2,064 (5.0) | 4,161 (4.8) | 2,064 (5.0) | 4,161 (4.8) |

| | All available data | | Complete covariate data[a] | |
|---|---|---|---|---|
| | Major depression | Comparison | Major depression | Comparison |
| Family history of severe depression | | | | |
| N (%) missing | 7,871 (15.4) | 15,347 (14.9) | 7,871 (15.4) | 15,347 (14.9) |
| N (%)[c] | 12,370 (28.7) | 10,201 (11.7) | 12,370 (28.7) | 10,201 (11.7) |
| Maternal smoking around birth | | | | |
| N (%) missing | 6,933 (13.6) | 13,107 (12.7) | 6,933 (13.6) | 13,107 (12.7) |
| N (%)[c] | 14,521 (33.0) | 24,125 (26.9) | 14,521 (33.0) | 24,125 (26.9) |
| **Mental health** | | | | |
| Current depressive symptoms | | | | |
| N (%) missing | 5,028 (9.9) | 8,556 (8.3) | 5,028 (9.9) | 8,556 (8.3) |
| Mean (SD) | 3.1 (3.0) | 1.2 (1.7) | 3.1 (3.0) | 1.2 (1.7) |
| Any psychotropic medication | | | | |
| N (%) missing | 860 (1.7) | 1,194 (1.2) | 860 (1.7) | 1,194 (1.2) |
| N (%)[c] | 20,898 (41.7) | 2,577 (2.5) | 20,898 (41.7) | 2,577 (2.5) |
| Number of depressed episodes | | | | |
| N (%) missing | 1,495 (2.9) | 6,451 (6.3) | 1,495 (2.9) | 6,451 (6.3) |
| Median (Q1, Q3) | 1 (0, 4) | 0 (0, 1) | 1 (0, 4) | 0 (0, 1) |
| Any childhood trauma[g] | | | | |
| N (%) missing | 34,801 (68.3) | 67,160 (65.3) | 34,801 (68.3) | 67,160 (65.3) |
| N (%)[c] | 9,455 (58.5) | 15,651 (43.8) | 9,455 (58.5) | 15,651 (43.8) |
| **Genome-wide polygenic scores** | | | | |
| Education/cognition GPS decile[d] | | | | |
| N (%) missing | 1,518 (3.0) | 3,419 (3.3) | 1,518 (3.0) | 3,419 (3.3) |
| D1 (lowest), N (%)[c] | 5,169 (10.5) | 9,728 (9.8) | 5,169 (10.5) | 9,728 (9.8) |
| D2 | 5,089 (10.3) | 9,808 (9.9) | 5,089 (10.3) | 9,808 (9.9) |
| D3 | 5,002 (10.1) | 9,895 (9.9) | 5,002 (10.1) | 9,895 (9.9) |
| D4 | 4,931 (10.0) | 9,966 (10.0) | 4,931 (10.0) | 9,966 (10.0) |
| D5 | 4,960 (10.0) | 9,937 (10.0) | 4,960 (10.0) | 9,937 (10.0) |
| D6 | 4,890 (9.9) | 10,007 (10.1) | 4,890 (9.9) | 10,007 (10.1) |
| D7 | 4,914 (9.9) | 9,983 (10.0) | 4,914 (9.9) | 9,983 (10.0) |
| D8 | 4,830 (9.8) | 10,067 (10.1) | 4,830 (9.8) | 10,067 (10.1) |
| D9 | 4,815 (9.7) | 10,082 (10.1) | 4,815 (9.7) | 10,082 (10.1) |
| D10 (highest) | 4,857 (9.8) | 10,039 (10.1) | 4,857 (9.8) | 10,039 (10.1) |
| Major depression GPS decile[d] | | | | |
| N (%) missing | 1,518 (3.0) | 3,419 (3.3) | 1,518 (3.0) | 3,419 (3.3) |
| D1 (lowest), N (%)[c] | 4,271 (8.6) | 10,626 (10.7) | 4,271 (8.6) | 10,626 (10.7) |
| D2 | 4,645 (9.4) | 10,252 (10.3) | 4,645 (9.4) | 10,252 (10.3) |
| D3 | 4,690 (9.5) | 10,207 (10.3) | 4,690 (9.5) | 10,207 (10.3) |
| D4 | 4,693 (9.5) | 10,204 (10.3) | 4,693 (9.5) | 10,204 (10.3) |
| D5 | 4,890 (9.9) | 10,007 (10.1) | 4,890 (9.9) | 10,007 (10.1) |
| D6 | 4,887 (9.9) | 10,010 (10.1) | 4,887 (9.9) | 10,010 (10.1) |
| D7 | 5,057 (10.2) | 9,840 (9.9) | 5,057 (10.2) | 9,840 (9.9) |
| D8 | 5,247 (10.6) | 9,650 (9.7) | 5,247 (10.6) | 9,650 (9.7) |
| D9 | 5,313 (10.7) | 9,584 (9.6) | 5,313 (10.7) | 9,584 (9.6) |
| D10 (highest) | 5,764 (11.7) | 9,162 (9.2) | 5,764 (11.7) | 9,162 (9.2) |

D, decile; GPS, genome-wide polygenic score; MET, metabolic equivalent of task; Q, quartile; Qu, quintile; SD, standard deviation.
a. Participants with complete data on all the covariates that were entered into the maximally-adjusted total effects models (age, gender, white British genetic ancestry, English-speaking country of birth, degree, comorbid neurological/psychiatric condition, family history of

dementia, family history of Parkinson's disease, family history of severe depression, maternal smoking around birth, childhood trauma, education/cognition GPS, major depression GPS).
b. No missing data.
c. Missing excluded from denominator.
d. Based on data distribution in the whole UK Biobank cohort.
e. Scottish psychiatric hospital records were unavailable, which meant no Scotland-based participants could be classified in the comparison group; therefore all locations for comparison participants are in England/Wales.
f. Apart from mood disorder or schizophrenia; not possible to distinguish between missing data and self-report of no condition, therefore both classified as 'No'.
g. From the web-based questionnaire, which was completed by 157,366 (31.3%) of the cohort.

#### Evaluation of the Graphical Model

The different predicted independencies implied by the two specifications of the DAG (educational attainment as an antecedent, or a consequence, of major depression and other psychiatric/neurological conditions) were tested, and better fit was evident in the first specification. Fifteen percent (21 of 137) of the partial correlation coefficients were above |0.1|, but most of these were below |0.2| and the largest coefficient was |0.22|.

#### Total Effects

The best covariate balance was obtained using the first propensity score model with no interaction terms, as illustrated in eTable 13. This was used in all the outcome models that involved propensity score adjustment or matching, or inverse probability weighting.

eFigures 10 to 13 show the total effects results for reasoning, reaction time, numeric memory and prospective memory.

*eTable 13 Summary of covariates in matched major depression and comparison groups*

| | Major depression | Comparison |
|---|---|---|
| | Mean (SD) | |
| Age (years) | 55.1 (7.5) | 55.2 (7.5) |
| | % | |
| Female gender | 68.4 | 69.5 |
| English-speaking country of birth | 98.3 | 98.2 |
| Has a degree | 46.0 | 46.5 |
| Comorbid neurological or psychiatric condition | 19.3 | 19.3 |
| Family history of dementia | 16.4 | 16.4 |
| Family history of Parkinson's disease | 4.4 | 4.3 |
| Family history of severe depression | 25.2 | 24.7 |
| Maternal smoking around birth | 31.2 | 30.7 |
| Any childhood trauma | 54.8 | 54.7 |
| Education/cognition GPS | | |
| D1 (lowest) | 8.6 | 8.2 |
| D2 | 9.4 | 9.4 |
| D3 | 9.5 | 9.5 |
| D4 | 9.7 | 9.5 |
| D5 | 9.7 | 9.8 |
| D6 | 9.8 | 9.8 |
| D7 | 10.6 | 10.9 |
| D8 | 10.1 | 10.1 |
| D9 | 11.1 | 11.1 |
| D10 (highest) | 11.5 | 11.7 |
| Major depression GPS | | |
| D1 (lowest) | 8.6 | 8.8 |
| D2 | 9.6 | 9.7 |
| D3 | 9.9 | 9.8 |
| D4 | 9.0 | 8.8 |
| D5 | 9.8 | 10.0 |
| D6 | 9.8 | 9.3 |
| D7 | 10.2 | 10.6 |
| D8 | 10.6 | 10.7 |
| D9 | 10.8 | 10.9 |
| D10 (highest) | 11.7 | 11.4 |

D, decile; GPS, genome-wide polygenic score; SD, standard deviation.
These results are from the matched samples used in the 1:3 matched model for the total effect of major depression on reaction time (major depression N = 9,381; comparison N = 13,538).

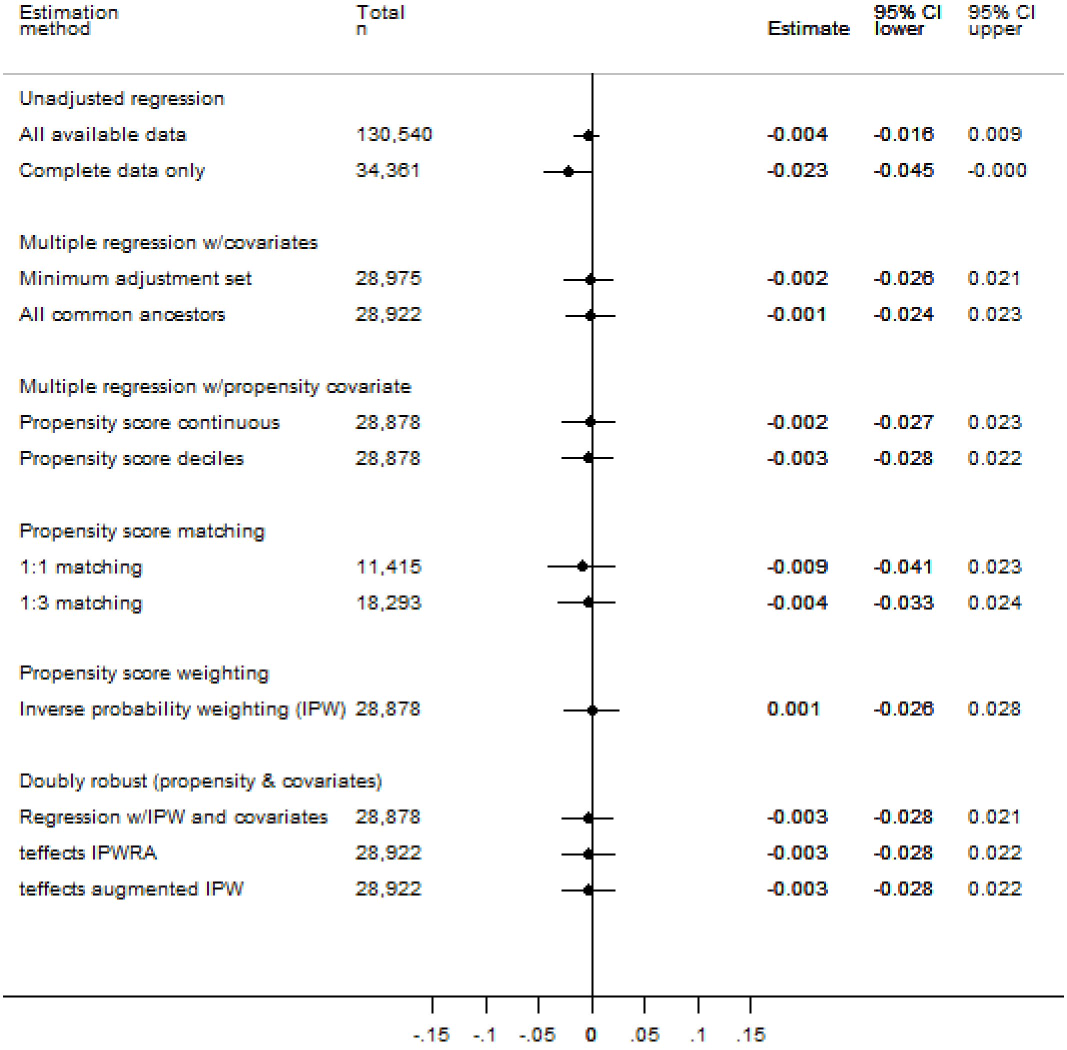

*eFigure 10 Total effect of major depression on reasoning*

CI, confidence interval; IPW, inverse probability weighting; IPWRA, inverse probability weighting with regression adjustment; teffects, Stata teffects package. Estimates are in z-score units and can be interpreted as standardized mean differences.

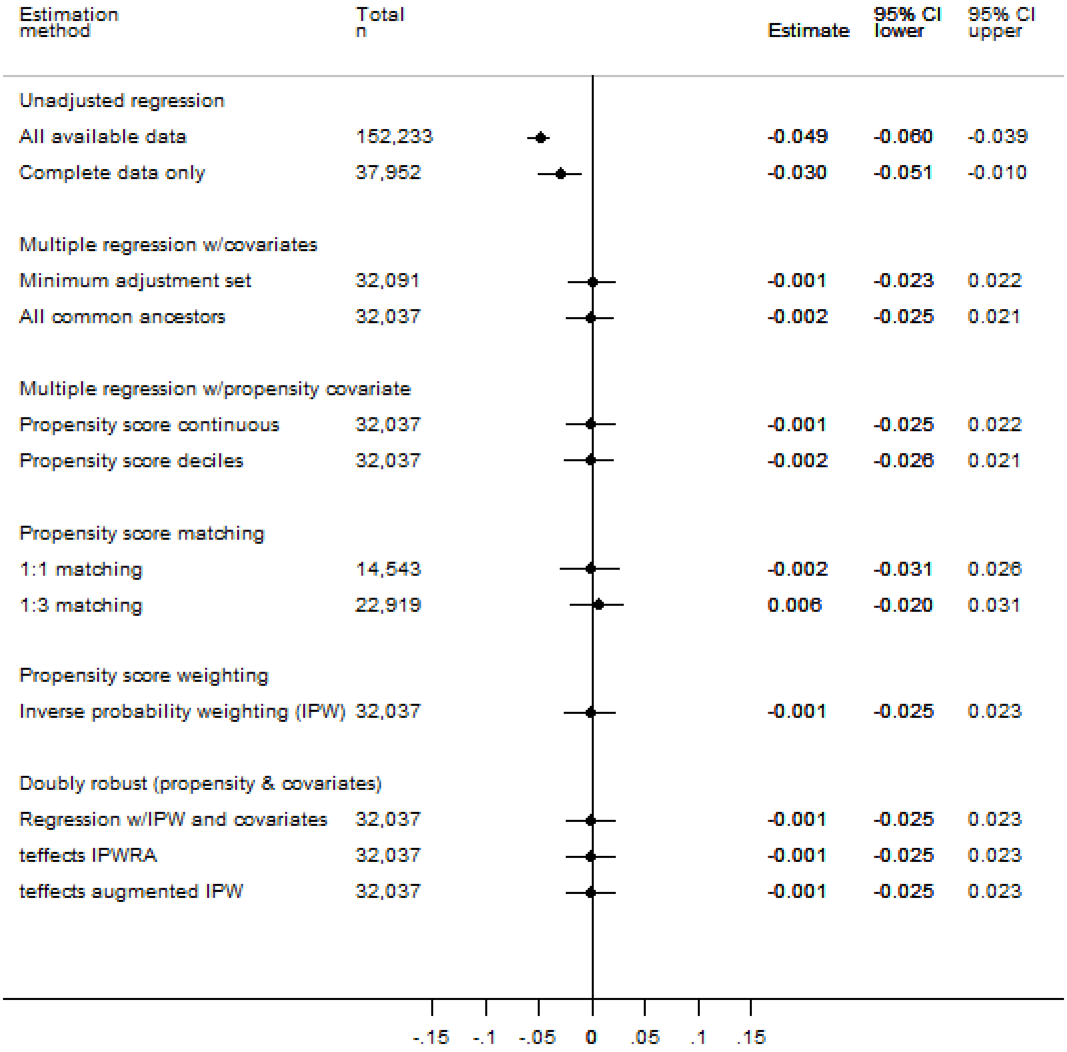

*eFigure 11 Total effect of major depression on reaction time*

CI, confidence interval; IPW, inverse probability weighting; IPWRA, inverse probability weighting with regression adjustment; teffects, Stata teffects package. Estimates are in z-score units and can be interpreted as standardized mean differences.

| Estimation method | Total n | Estimate | 95% CI lower | 95% CI upper |
|---|---|---|---|---|
| Unadjusted regression | | | | |
| All available data | 39,082 | -0.037 | -0.059 | -0.016 |
| Complete data only | 9,936 | -0.074 | -0.116 | -0.033 |
| Multiple regression w/covariates | | | | |
| Minimum adjustment set | 8,857 | -0.039 | -0.084 | 0.006 |
| All common ancestors | 8,835 | -0.034 | -0.079 | 0.012 |
| Multiple regression w/propensity covariate | | | | |
| Propensity score continuous | 8,821 | -0.035 | -0.081 | 0.011 |
| Propensity score deciles | 8,821 | -0.039 | -0.085 | 0.007 |
| Propensity score matching | | | | |
| 1:1 matching | 3,272 | -0.029 | -0.087 | 0.030 |
| 1:3 matching | 5,575 | -0.019 | -0.071 | 0.033 |
| Propensity score weighting | | | | |
| Inverse probability weighting (IPW) | 8,821 | -0.026 | -0.075 | 0.022 |
| Doubly robust (propensity & covariates) | | | | |
| Regression w/IPW and covariates | 8,821 | -0.029 | -0.076 | 0.018 |
| teffects IPWRA | 8,835 | -0.028 | -0.076 | 0.019 |
| teffects augmented IPW | 8,835 | -0.028 | -0.076 | 0.019 |

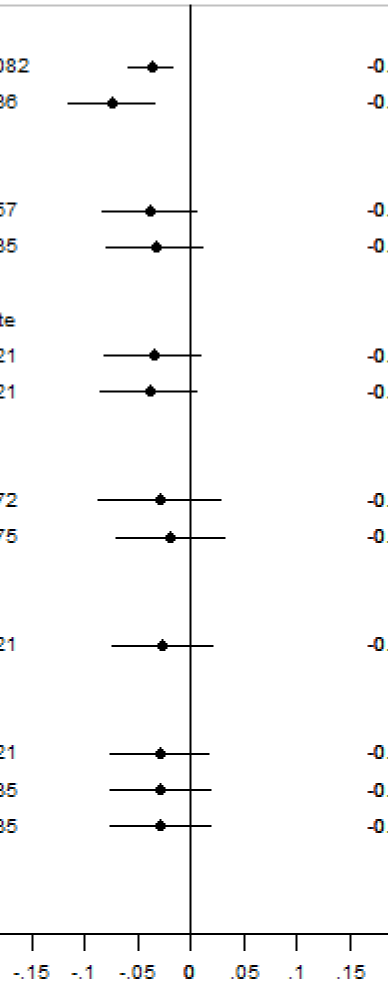

*eFigure 12 Total effect of major depression on numeric memory*

CI, confidence interval; IPW, inverse probability weighting; IPWRA, inverse probability weighting with regression adjustment; teffects, Stata teffects package. Estimates are in z-score units and can be interpreted as standardized mean differences.

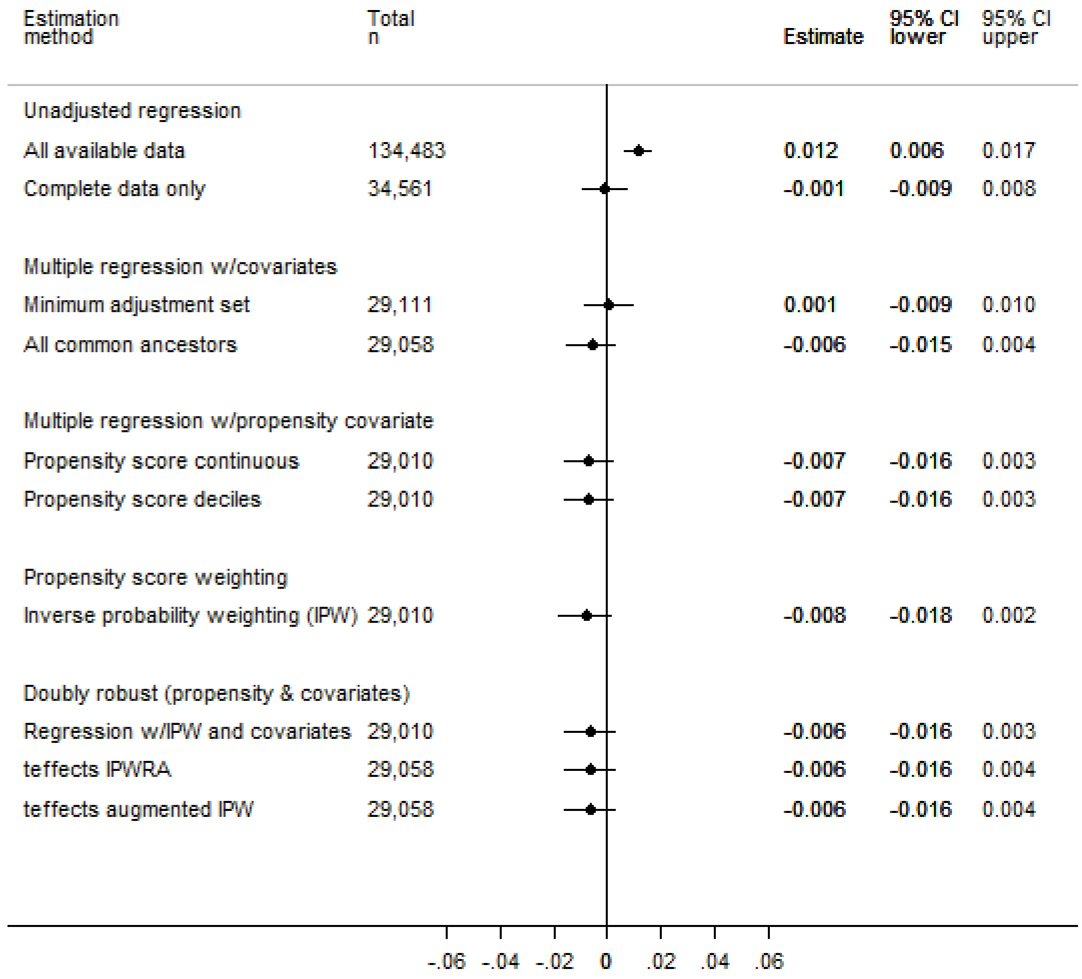

*eFigure 13 Total effect of major depression on prospective memory*

CI, confidence interval; IPW, inverse probability weighting; IPWRA, inverse probability weighting with regression adjustment; teffects, Stata teffects package. Estimates are proportions and can be interpreted as risk differences. No estimates are provided from propensity-score matched models as it was not possible to express these as risk differences.

#### Mediation Analyses

In both sets of mediation analyses, the indirect pathways were affected by intermediate confounding, and so an identifying assumption was firstly made of no interaction between major depression and either cardiometabolic disease or psychotropic medication.[61] This assumption was checked by conducting a regression model of each cognitive outcome on major depression exposure status, the mediator and all the covariates, including a product term for major depression * mediator. There was no evidence of interaction in the cardiometabolic disease mediation models, but some evidence of interaction in the psychotropic medication models (eTable 14 below). The alternative identifying assumption proposed by Peterson et al.[62] was therefore made for the psychotropic medication models; following De Stavola et al.[59] this was checked by testing for interactions between major depression and each of the intermediate confounders (deprivation, and lifetime number of episodes of depressed mood/anhedonia), in regression models that included major depression, psychotropic medication and their product, along with all the other model covariates. There was little evidence of interaction between major depression and the intermediate confounders (eTable 15), and so this identifying assumption was deemed reasonable and these mediation models were estimated with a product term included between major depression and psychotropic medication.

*eTable 14 Tests of interactions between exposure and mediators in the major depression analyses*

| | Coefficient for major depression * mediator | 95% CI | *P* |
|---|---|---|---|
| **Mediator: Cardiometabolic disease** | | | |
| Reasoning[a] | 0.029 | -0.027, 0.085 | 0.304 |
| Reaction time[a] | 0.053 | -0.001, 0.107 | 0.056 |
| Numeric memory[a] | 0.057 | -0.052, 0.166 | 0.306 |
| Visuospatial memory[a] | 0.036 | -0.024, 0.095 | 0.241 |
| Prospective memory[b] | 1.157 | 0.952, 1.407 | 0.142 |
| **Mediator: Psychotropic medication** | | | |
| Reasoning[a] | 0.244 | 0.135, 0.354 | <0.001 |
| Reaction time[a] | 0.080 | -0.025, 0.185 | 0.134 |
| Numeric memory[a] | 0.203 | 0.013, 0.393 | 0.036 |
| Visuospatial memory[a] | 0.082 | -0.037, 0.200 | 0.176 |
| Prospective memory[b] | 1.454 | 1.057, 2.000 | 0.021 |

CI, confidence interval.
All models included major depression, the mediator and their product, as well as all the covariates entered into the gformula mediation models.
a. Linear regression model with outcome measured in z-score units.
b. Logistic regression model with outcome measured as correct or not; estimate expressed as odds ratio.

*eTable 15 Tests of interactions between exposure and intermediate confounders*

| | **Coefficient for major depression * deprivation** | **95% CI** | ***P*** | **Coefficient for major depression * lifetime number of episodes of depressed mood/anhedonia** | **95% CI** | ***P*** |
|---|---|---|---|---|---|---|
| Reasoning[a] | 0.018 | -0.030, 0.066 | 0.458 | -0.020 | -0.074, 0.035 | 0.481 |
| Reaction time[a] | -0.017 | -0.064, 0.030 | 0.475 | 0.033 | -0.020, 0.085 | 0.224 |
| Numeric memory[a] | 0.041 | -0.057, 0.138 | 0.413 | -0.002 | -0.106, 0.103 | 0.974 |
| Visuospatial memory[a] | 0.056 | 0.005, 0.108 | 0.033 | -0.023 | -0.081, 0.035 | 0.433 |
| Prospective memory[b] | 0.989 | 0.836, 1.171 | 0.900 | 1.000 | 0.834, 1.212 | 0.996 |

CI, confidence interval.
Deprivation was entered as a dichotomous indicator for the two most deprived Townsend quintiles versus the three least deprived quintiles. Lifetime number of episodes of depressed mood/anhedonia was entered as a dichotomous indicator for ≥2 episodes versus <2 episodes. All models included the product terms indicated in the table above, as well as major depression, deprivation, lifetime number of episodes of depressed mood/anhedonia, psychotropic medication, major depression * psychotropic medication, and all the other covariates entered into the gformula mediation models.
a. Linear regression model with outcome measured in z-score units.
b. Logistic regression model with outcome measured as correct or not; estimate expressed as odds ratio.

*eTable 16 Mediation of the effect of major depression on cognitive outcome via cardiometabolic disease*

| | N | Estimate | 95% CI[a] |
|---|---|---|---|
| **Reasoning**[b] | 26,679 | | |
| TE | | -0.009 | -0.038, 0.019 |
| CDE | | -0.029 | -0.056, -0.001 |
| NDE | | 0.005 | -0.023, 0.033 |
| NIE | | -0.014 | -0.028, 0.000 |
| **Reaction time**[b] | 29,422 | | |
| TE | | -0.015 | -0.042, 0.013 |
| CDE | | -0.006 | -0.036, 0.024 |
| NDE | | -0.009 | -0.037, 0.020 |
| NIE | | -0.006 | -0.021, 0.009 |
| **Numeric memory**[b] | 8,085 | | |
| TE | | -0.034 | -0.087, 0.019 |
| CDE | | -0.032 | -0.085, 0.021 |
| NDE | | -0.035 | -0.089, 0.018 |
| NIE | | 0.001 | -0.027, 0.029 |
| **Visuospatial memory**[b] | 29,284 | | |
| TE | | -0.074 | -0.107, -0.042 |
| CDE | | -0.077 | -0.109, -0.046 |
| NDE | | -0.079 | -0.111, -0.047 |
| NIE | | 0.005 | -0.012, 0.022 |
| **Prospective memory**[c] | 26,789 | | |
| TE | | -0.010 | -0.022, 0.001 |
| CDE | | -0.001 | -0.012, 0.010 |
| NDE | | -0.011 | -0.022, 0.000 |
| NIE | | 0.001 | -0.012, 0.010 |

CDE, controlled direct effect when cardiometabolic disease = 0; CI, confidence interval; GPS, genome-wide polygenic score; NDE, natural direct effect; NIE, natural indirect effect; $NO_2$, nitrogen dioxide; $PM_{10}$, particulate matter of up to 10μm diameter; TE, total effect. Models were restricted to participants of white British genetic ancestry, and were adjusted for age, gender, educational attainment, English-speaking birth country, education/cognition GPS, major depression GPS, family history of dementia, family history of Parkinson's disease, maternal smoking around birth, childhood trauma, other psychiatric/neurological conditions, deprivation, population density, road proximity, air pollution ($PM_{10}$ and $NO_2$), body mass index, alcohol frequency, smoking status, physical activity, and psychotropic medication.

a. Normal-based, from bootstrapped standard error (1000 replicates).

b. Estimate expressed as a standardized mean difference.

c. Estimate expressed as a risk difference for the probability of being correct.

*eTable 17 Mediation of the effect of major depression on cognitive outcome via psychotropic medication*

| | N | Estimate | 95% CI[a] |
|---|---|---|---|
| **Reasoning**[b] | 27,263 | | |
| TE | | -0.009 | -0.037, 0.019 |
| CDE | | -0.011 | -0.040, 0.018 |
| NDE | | -0.010 | -0.039, 0.020 |
| NIE | | 0.001 | -0.016, 0.017 |
| **Reaction time**[b] | 30,189 | | |
| TE | | 0.002 | -0.026, 0.029 |
| CDE | | 0.005 | -0.025, 0.036 |
| NDE | | 0.026 | -0.004, 0.055 |
| NIE | | -0.024 | -0.042, -0.005 |
| **Numeric memory**[b] | 8,338 | | |
| TE | | -0.023 | -0.078, 0.032 |
| CDE | | -0.021 | -0.077, 0.035 |
| NDE | | -0.019 | -0.076, 0.038 |
| NIE | | -0.004 | -0.035, 0.028 |
| **Visuospatial memory**[b] | 30,038 | | |
| TE | | -0.058 | -0.088, -0.028 |
| CDE | | -0.066 | -0.100, -0.031 |
| NDE | | -0.039 | -0.073, -0.006 |
| NIE | | -0.019 | -0.040, 0.003 |
| **Prospective memory**[c] | 27,381 | | |
| TE | | 0.001 | -0.011, 0.012 |
| CDE | | 0.001 | -0.010, 0.013 |
| NDE | | 0.001 | -0.011, 0.012 |
| NIE | | 0.000 | -0.006, 0.006 |

CDE, controlled direct effect when psychotropic medication = 0; CI, confidence interval; GPS, genome-wide polygenic score; NDE, natural direct effect; NIE, natural indirect effect; TE, total effect.
Models were restricted to participants of white British genetic ancestry, and were adjusted for gender, educational attainment, English-speaking birth country, education/cognition GPS, major depression GPS, family history of dementia, family history of Parkinson's disease, maternal smoking around birth, childhood trauma, other psychiatric/neurological conditions, deprivation, and lifetime number of episodes of depressed mood/anhedonia. All models included a product term for major depression * psychotropic medication.
a. Normal-based, from bootstrapped standard error (1000 replicates).
b. Estimate expressed as a standardized mean difference.
c. Estimate expressed as a risk difference for the probability of being correct.

#### Sensitivity Analyses

Rosenbaum bounds were calculated to check the sensitivity of the visuospatial memory total effect result to departures from exchangeability. The estimated effect crossed the null at a gamma value of 1.07, i.e. the point where the probability of being in the exposed group is approximately 0.48 or 0.52. The results would therefore not be robust to an unmeasured confounder with even a very weak association with group membership.

There was evidence of missing data bias, in that the unadjusted total effects estimates shifted (towards or away from the null) when the sample was restricted to participants with complete covariate data. When the multiple linear regression models for total effects were repeated using imputed covariate values, the estimates for reaction time indicated a very small detrimental effect (point estimate approximately -0.02 to -0.03) in the major depression group (eFigure 14(b) below), which was not evident in the complete case analyses. The estimates for the other cognitive outcomes were similar between the complete case analyses and those using multiple imputation.

When the mediation models were repeated with imputation of missing mediator and covariate values, there remained no evidence of indirect effects via cardiometabolic disease (eTable 18). There was evidence of an indirect effect via psychotropic medication on visuospatial memory performance (eTable 19), accounting for approximately 18% of the total effect.

The results of the probabilistic analysis using `episens` indicated that the total effects estimate for visuospatial memory would be biased away from the null if there were differential misclassification of the exposure. When dichotomized into impaired and unimpaired outcome categories, and assuming no exposure misclassification, the unadjusted relative risk of impairment was 1.14 in the major depression group (95% CI 1.07, 1.20). Assuming lower sensitivity to true major depression status among the cognitively impaired (sensitivity range 0.6 to 0.9) versus unimpaired participants (sensitivity range 0.7 to 1.0), the relative risk was estimated as 1.37 (1.06, 1.75).

DAGitty determined that there were six other DAGs that were equivalent to the DAG used in the analyses. None of these alternative configurations was causally plausible, owing to temporal order constraints. The same minimum adjustment set was valid for the analysed DAG and for the six equivalent DAGs, for estimating the total effect of major depression on cognitive outcome.

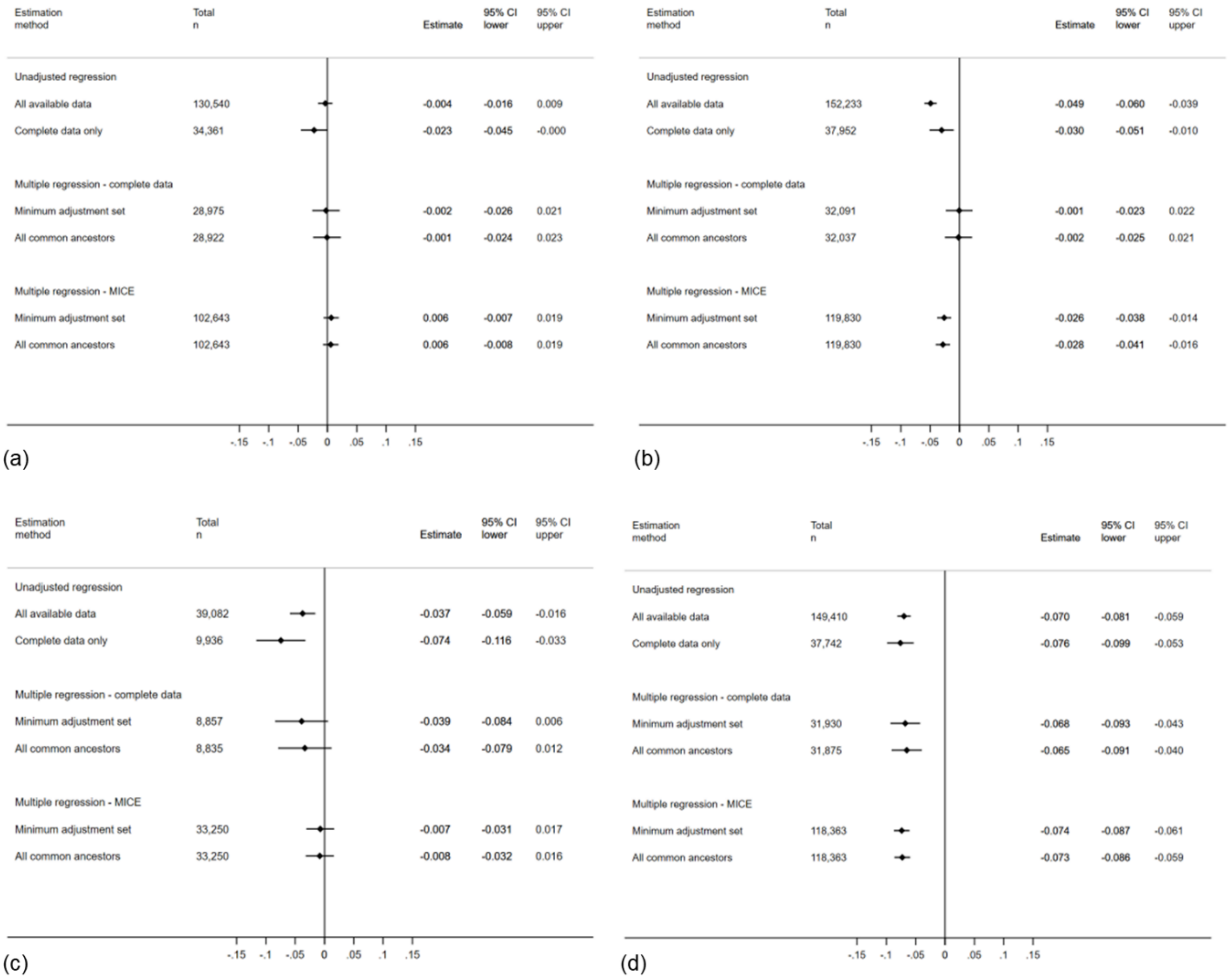

*eFigure 14 Comparison of missing data approaches in major depression total effects analyses*

CI, confidence interval; MICE, multiple imputation with chained equations. Panels show: (a) reasoning, (b) reaction time, (c) numeric memory and (d) visuospatial memory. Prospective memory not shown as it was not possible to calculate risk differences.

*eTable 18 Mediation of the effect of major depression on cognitive outcome via cardiometabolic disease, with missing data imputation*

| | N | Estimate | 95% CI[a] |
|---|---|---|---|
| **Reasoning**[b] | 102,643 | | |
| TE | | 0.001 | -0.017, 0.019 |
| CDE | | -0.005 | -0.023, 0.013 |
| NDE | | -0.009 | -0.026, 0.009 |
| NIE | | 0.009 | -0.023, 0.013 |
| **Reaction time**[b] | 119,830 | | |
| TE | | -0.024 | -0.042, -0.006 |
| CDE | | -0.022 | -0.039, -0.004 |
| NDE | | -0.032 | -0.050, -0.014 |
| NIE | | 0.008 | -0.002, 0.019 |
| **Numeric memory**[b] | 33,250 | | |
| TE | | -0.006 | -0.039, 0.027 |
| CDE | | -0.013 | -0.047, 0.020 |
| NDE | | -0.006 | -0.039, 0.027 |
| NIE | | 0.000 | -0.016, 0.017 |
| **Visuospatial memory**[b] | 118,363 | | |
| TE | | -0.072 | -0.091, -0.054 |
| CDE | | -0.074 | -0.093, -0.056 |
| NDE | | -0.072 | -0.091 -0.053 |
| NIE | | -0.001 | -0.011, 0.010 |
| **Prospective memory**[c] | 104,509 | | |
| TE | | -0.004 | -0.009, 0.001 |
| CDE | | -0.002 | -0.007, 0.003 |
| NDE | | -0.004 | -0.009, 0.002 |
| NIE | | -0.000 | -0.003, 0.002 |

CDE, controlled direct effect when cardiometabolic disease = 0; CI, confidence interval; GPS, genome-wide polygenic score; NDE, natural direct effect; NIE, natural indirect effect; $NO_2$, nitrogen dioxide; $PM_{10}$, particulate matter of up to 10μm diameter; TE, total effect. Models were restricted to participants of white British genetic ancestry, and were adjusted for age, gender, educational attainment, English-speaking birth country, education/cognition GPS, major depression GPS, family history of dementia, family history of Parkinson's disease, maternal smoking around birth, childhood trauma, other psychiatric/neurological conditions, deprivation, population density, road proximity, air pollution ($PM_{10}$ and $NO_2$), body mass index, alcohol frequency, smoking status, physical activity, and psychotropic medication. Missing mediator and covariate data were imputed via a single stochastic imputation using chained equations.

a. Normal-based, from bootstrapped standard error (1000 replicates).

b. Estimate expressed as a standardized mean difference.

c. Estimate expressed as a risk difference for the probability of being correct.

*eTable 19 Mediation of the effect of major depression on cognitive outcome via psychotropic medication, with missing data imputation*

| | N | Estimate | 95% CI[a] |
|---|---|---|---|
| **Reasoning**[b] | 102,643 | | |
| TE | | -0.004 | -0.022, 0.014 |
| CDE | | 0.003 | -0.016, 0.021 |
| NDE | | -0.010 | -0.028, 0.008 |
| NIE | | 0.006 | -0.003, 0.015 |
| **Reaction time**[b] | 119,830 | | |
| TE | | -0.009 | -0.026, 0.008 |
| CDE | | 0.011 | -0.009, 0.030 |
| NDE | | -0.008 | -0.026, 0.009 |
| NIE | | -0.001 | -0.011, 0.010 |
| **Numeric memory**[b] | 33,250 | | |
| TE | | -0.022 | -0.056, 0.011 |
| CDE | | -0.001 | -0.035, 0.034 |
| NDE | | -0.005 | -0.040, 0.030 |
| NIE | | -0.017 | -0.036, 0.002 |
| **Visuospatial memory**[b] | 118,363 | | |
| TE | | -0.068 | -0.087, -0.049 |
| CDE | | -0.035 | -0.057, -0.014 |
| NDE | | -0.056 | -0.074, -0.037 |
| NIE | | -0.012 | -0.023, -0.001 |
| **Prospective memory**[c] | 104,509 | | |
| TE | | 0.002 | -0.003, 0.008 |
| CDE | | 0.007 | 0.001, 0.013 |
| NDE | | 0.001 | -0.004, 0.006 |
| NIE | | 0.001 | -0.002, 0.004 |

CDE, controlled direct effect when psychotropic medication = 0; CI, confidence interval; GPS, genome-wide polygenic score; NDE, natural direct effect; NIE, natural indirect effect; TE, total effect.
Models were restricted to participants of white British genetic ancestry, and were adjusted for gender, educational attainment, English-speaking birth country, education/cognition GPS, major depression GPS, family history of dementia, family history of Parkinson's disease, maternal smoking around birth, childhood trauma, other psychiatric/neurological conditions, deprivation, and lifetime number of episodes of depressed mood/anhedonia. All models included a product term for major depression * psychotropic medication. Missing mediator and covariate data were imputed via a single stochastic imputation using chained equations.
a. Normal-based, from bootstrapped standard error (1000 replicates).
b. Estimate expressed as a standardized mean difference.
c. Estimate expressed as a risk difference for the probability of being correct.

### eReferences

1. Smith DJ, Nicholl BI, Cullen B, et al. Prevalence and Characteristics of Probable Major Depression and Bipolar Disorder within UK Biobank: Cross-Sectional Study of 172,751 Participants. *PLoS One* 2013; **8**(11): e75362.
2. Townsend P. Deprivation. *J Soc Policy* 1987; **16**: 125-46.
3. Vienneau D, de Hoogh K, Bechle MJ, et al. Western European Land Use Regression Incorporating Satellite- and Ground-Based Measurements of NO2 and PM10. *Environmental Science & Technology* 2013; **47**(23): 13555-64.
4. Booth M. Assessment of physical activity: An international perspective. *Res Q Exercise Sport* 2000; **71**(2): S114-S20.
5. Kroenke K, Spitzer RL, Williams JB. The PHQ-9: validity of a brief depression severity measure. *J Gen Intern Med* 2001; **16**(9): 606-13.
6. Bernstein DP, Stein JA, Newcomb MD, et al. Development and validation of a brief screening version of the Childhood Trauma Questionnaire. *Child Abuse Neglect* 2003; **27**(2): 169-90.
7. Walker EA, Gelfand A, Katon WJ, et al. Adult health status of women with histories of childhood abuse and neglect. *Am J Med* 1999; **107**(4): 332-9.
8. Bycroft C, Freeman C, Petkova D, et al. Genome-wide genetic data on ~500,000 UK Biobank participants. *bioRXiv* 2017.
9. Okbay A, Beauchamp JP, Fontana MA, et al. Genome-wide association study identifies 74 loci associated with educational attainment. *Nature* 2016; **533**(7604): 539-42.
10. Hill WD, Marioni RE, Maghzian O, et al. A combined analysis of genetically correlated traits identifies 187 loci and a role for neurogenesis and myelination in intelligence. *Molecular Psychiatry* 2018; **[Epub ahead of print]**.
11. Plomin R, von Stumm S. The new genetics of intelligence. *Nature Reviews Genetics* 2018; **19**(3): 148-59.
12. Sklar P, Ripke S, Scott LJ, et al. Large-scale genome-wide association analysis of bipolar disorder identifies a new susceptibility locus near ODZ4. *Nature Genetics* 2011; **43**(10): 977-83.
13. Wray NR, Ripke S, Mattheisen M, et al. Genome-wide association analyses identify 44 risk variants and refine the genetic architecture of major depression. *Nature Genetics* 2018; **50**(5): 668-81.
14. de Zeeuw EL, van Beijsterveldt CEM, Glasner TJ, et al. Polygenic Scores Associated With Educational Attainment in Adults Predict Educational Achievement and ADHD Symptoms in Children. *American Journal of Medical Genetics Part B-Neuropsychiatric Genetics* 2014; **165**(6): 510-20.
15. Krapohl E, Plomin R. Genetic link between family socioeconomic status and children's educational achievement estimated from genome-wide SNPs. *Molecular Psychiatry* 2016; **21**(3): 437-43.
16. Krapohl E, Euesden J, Zabaneh D, et al. Phenome-wide analysis of genome-wide polygenic scores. *Molecular Psychiatry* 2016; **21**(9): 1188-93.
17. Selzam S, Krapohl E, von Stumm S, et al. Predicting educational achievement from DNA. *Molecular Psychiatry* 2017; **22**(2): 267-72.
18. Aminoff SR, Tesli M, Bettella F, et al. Polygenic risk scores in bipolar disorder subgroups. *J Affect Disord* 2015; **183**: 310-4.
19. Elwert F. Graphical causal models. In: Morgan SL, ed. Handbook of Causal Analysis for Social Research. New York: Springer; 2013: 245-73.
20. Pearl J, Glymour M, Jewell NP. Causal Inference in Statistics: A Primer. Chichester, West Sussex: Wiley; 2016.
21. Deary IJ, Corley J, Gow AJ, et al. Age-associated cognitive decline. *British Medical Bulletin* 2009; **92**(1): 135-52.
22. Miller DI, Halpern DF. The new science of cognitive sex differences. *Trends in Cognitive Sciences* 2014; **18**(1): 37-45.
23. Blanco C, Compton WM, Saha TD, et al. Epidemiology of DSM-5 bipolar I disorder: Results from the National Epidemiologic Survey on Alcohol and Related Conditions - III. *J Psychiatr Res* 2017; **84**: 310-7.
24. Lloyd T, Kennedy N, Fearon P, et al. Incidence of bipolar affective disorder in three UK cities: results from the AESOP study. *Br J Psychiatry* 2005; **186**: 126-31.
25. Hill WD, Davies G, Liewald DC, McIntosh AM, Deary IJ, CHARGE Cognitive Working Group. Age-Dependent Pleiotropy Between General Cognitive Function and Major Psychiatric Disorders. *Biol Psychiatry* 2016; **80**(4): 266-73.
26. Hagenaars SP, Harris SE, Davies G, et al. Shared genetic aetiology between cognitive functions and physical and mental health in UK Biobank (N=112 151) and 24 GWAS consortia. *Molecular Psychiatry* 2016; **21**(11): 1624-32.
27. Talati A, Bao Y, Kaufman J, Shen L, Schaefer CA, Brown AS. Maternal smoking during pregnancy and bipolar disorder in offspring. *American Journal of Psychiatry* 2013; **170**(10): 1178-85.
28. Aas M, Henry C, Andreassen OA, Bellivier F, Melle I, Etain B. The role of childhood trauma in bipolar disorders. *International Journal of Bipolar Disorders* 2016; **4**: 2.
29. Mortensen PB, Pedersen CB, Melbye M, Mors O, Ewald H. Individual and familial risk factors for bipolar affective disorders in Denmark. *Arch Gen Psychiatry* 2003; **60**(12): 1209-15.

30. Scott EM, Hermens DF, White D, et al. Body mass, cardiovascular risk and metabolic characteristics of young persons presenting for mental healthcare in Sydney, Australia. *BMJ Open* 2015; **5**(3): e007066.
31. Baumgart M, Snyder HM, Carrillo MC, Fazio S, Kim H, Johns H. Summary of the evidence on modifiable risk factors for cognitive decline and dementia: A population-based perspective. *Alzheimers Dement* 2015; **11**(6): 718-26.
32. Kemp DE, Fan JB. Cardiometabolic Health in Bipolar Disorder. *Psychiatric Annals* 2012; **42**(5): 179-83.
33. Fiedorowicz JG, He J, Merikangas KR. The association between mood and anxiety disorders with vascular diseases and risk factors in a nationally representative sample. *Journal of Psychosomatic Research* 2011; **70**(2): 145-54.
34. Knopman D, Boland LL, Mosley T, et al. Cardiovascular risk factors and cognitive decline in middle-aged adults. *Neurology* 2001; **56**(1): 42-8.
35. Qiu C, Fratiglioni L. A major role for cardiovascular burden in age-related cognitive decline. *Nature Reviews Cardiology* 2015; **12**(5): 267-77.
36. Gitlin MJ, Miklowitz DJ. The difficult lives of individuals with bipolar disorder: A review of functional outcomes and their implications for treatment. *J Affect Disord* 2017; **209**: 147-54.
37. Lyu J, Burr JA. Socioeconomic Status Across the Life Course and Cognitive Function Among Older Adults: An Examination of the Latency, Pathways, and Accumulation Hypotheses. *J Aging Health* 2016; **28**(1): 40-67.
38. Marioni RE, Davies G, Hayward C, et al. Molecular genetic contributions to socioeconomic status and intelligence. *Intelligence* 2014; **44**: 26-32.
39. Killin LOJ, Starr JM, Shiue IJ, Russ TC. Environmental risk factors for dementia: a systematic review. *BMC Geriatrics* 2016; **16**: 175.
40. Power MC, Adar SD, Yanosky JD, Weuve J. Exposure to air pollution as a potential contributor to cognitive function, cognitive decline, brain imaging, and dementia: A systematic review of epidemiologic research. *Neurotoxicology* 2016; **56**: 235-53.
41. Bourne C, Aydemir O, Balanza-Martinez V, et al. Neuropsychological testing of cognitive impairment in euthymic bipolar disorder: an individual patient data meta-analysis. *Acta Psychiatr Scand* 2013; **128**(3): 149-62.
42. Ganguli M, Du YC, Dodge HH, Ratcliff GG, Chang CCH. Depressive symptoms and cognitive decline in late life - A prospective epidemiological study. *Arch Gen Psychiatry* 2006; **63**(2): 153-60.
43. Harvey AG, Schmidt DA, Scarna A, Semler CN, Goodwin GM. Sleep-related functioning in euthymic patients with bipolar disorder, patients with insomnia, and subjects without sleep problems. *American Journal of Psychiatry* 2005; **162**(1): 50-7.
44. Fortier-Brochu E, Morin CM. Cognitive Impairment in Individuals with Insomnia: Clinical Significance and Correlates. *Sleep* 2014; **37**(11): 1789-800.
45. Goldman-Mellor S, Caspi A, Gregory AM, Harrington H, Poulton R, Moffitt TE. Is Insomnia Associated with Deficits in Neuropsychological Functioning? Evidence from a Population-Based Study. *Sleep* 2015; **38**(4): 623-31.
46. Robinson LJ, Ferrier IN. Evolution of cognitive impairment in bipolar disorder: a systematic review of cross-sectional evidence. *Bipolar Disord* 2006; **8**(2): 103-16.
47. Balanzá-Martínez V, Selva G, Martinez-Aran A, et al. Neurocognition in bipolar disorders-A closer look at comorbidities and medications. *European Journal of Pharmacology* 2010; **626**(1): 87-96.
48. Lim CS, Baldessarini RJ, Vieta E, Yucel M, Bora E, Sim K. Longitudinal neuroimaging and neuropsychological changes in bipolar disorder patients: Review of the evidence. *Neuroscience and Biobehavioral Reviews* 2013; **37**(3): 418-35.
49. Miskowiak KW, Kjaerstad HL, Meluken I, et al. The search for neuroimaging and cognitive endophenotypes: A critical systematic review of studies involving unaffected first-degree relatives of individuals with bipolar disorder. *Neuroscience and Biobehavioral Reviews* 2017; **73**: 1-22.
50. Textor J, van der Zander B, Gilthorpe MS, Liskiewicz M, Ellison GTH. Robust causal inference using directed acyclic graphs: the R package 'dagitty'. *International Journal of Epidemiology* 2016; **45**(6): 1887-94.
51. Brookhart MA, Schneeweiss S, Rothman KJ, Glynn RJ, Avorn J, Sturmer T. Variable selection for propensity score models. *Am J Epidemiol* 2006; **163**(12): 1149-56.
52. Friedman JH. Greedy function approximation: A gradient boosting machine. *Ann Stat* 2001; **29**(5): 1189-232.
53. Garrido MM, Kelley AS, Paris J, et al. Methods for Constructing and Assessing Propensity Scores. *Health Services Research* 2014; **49**(5): 1701-20.
54. StataCorp. Stata Statistical Software: Release 15. College Station, TX: StataCorp LP; 2017.
55. Austin PC. An Introduction to Propensity Score Methods for Reducing the Effects of Confounding in Observational Studies. *Multivar Behav Res* 2011; **46**(3): 399-424.
56. Rosenbaum PR. Observational Studies. 2nd ed. New York: Springer; 2002.
57. White IR, Royston P, Wood AM. Multiple imputation using chained equations: Issues and guidance for practice. *Stat Med* 2011; **30**(4): 377-99.
58. Cullen B, Smith DJ, Deary IJ, Evans JJ, Pell JP. The 'cognitive footprint' of psychiatric and neurological conditions: cross-sectional study in the UK Biobank cohort. *Acta Psychiatr Scand* 2017; **135**(6): 593-605.
59. De Stavola BL, Daniel RM, Ploubidis GB, Micali N. Mediation Analysis With Intermediate Confounding: Structural Equation Modeling Viewed Through the Causal Inference Lens. *Am J Epidemiol* 2015; **181**(1): 64-80.

60. VanderWeele TJ, Vansteelandt S, Robins JM. Effect Decomposition in the Presence of an Exposure-Induced Mediator-Outcome Confounder. *Epidemiology* 2014; **25**(2): 300-6.
61. Robins JM, Greenland S. Identifiability and Exchangeability for Direct and Indirect Effects. *Epidemiology* 1992; **3**(2): 143-55.
62. Petersen ML, Sinisi SE, van der Laan MJ. Estimation of direct causal effects. *Epidemiology* 2006; **17**(3): 276-84.
